# Discovery of a Cellular Mechanism Regulating Transcriptional Noise

**DOI:** 10.1101/2020.06.29.128439

**Authors:** Ravi V. Desai, Maike M.K. Hansen, Benjamin Martin, Chen Yu, Sheng Ding, Matt Thomson, Leor S. Weinberger

## Abstract

Stochastic fluctuations in gene expression (‘noise’) are often considered detrimental but, in other fields, fluctuations are harnessed for benefit (e.g., ‘dither’ or amplification of thermal fluctuations to accelerate chemical reactions). Here, we find that DNA base-excision repair amplifies transcriptional noise, generating increased cellular plasticity and facilitating reprogramming. The DNA-repair protein Apex1 recognizes modified nucleoside substrates to amplify expression noise—while homeostatically maintaining mean levels of expression— for virtually all genes across the transcriptome. This noise amplification occurs for both naturally occurring base modifications and unnatural base analogs. Single-molecule imaging shows amplified noise originates from shorter, but more intense, transcriptional bursts that occur via increased DNA supercoiling which first impedes and then accelerates transcription, thereby maintaining mean levels. Strikingly, homeostatic noise amplification potentiates fate-conversion signals during cellular reprogramming. These data suggest a functional role for the observed occurrence of modified bases within DNA in embryonic development and disease.

## Main Text

From Brownian motion to electrical ‘shot’ noise, fluctuations are fundamental to physical processes. Since the 1800s (*1*), fluctuations have been recognized to dynamically shape the distribution of microstates a system adopts, and modulation of fluctuations has been harnessed throughout engineering and the sciences. For example, in chemistry, thermal fluctuations— amplified via temperature increase (e.g., Bunsen Burners)—accelerate reactions (*2*); in engineering, amplification of electrical, acoustic, or mechanical fluctuations (i.e., ‘dither’, from the Middle English “didderen” meaning to “tremble”) is used for signal recovery (*3*), and in neuroscience, electrophysiological fluctuations—first reported in the 1950s (*4*)—are clinically amplified to improve sensorimotor function (*5-7*). Such ‘dither’ approaches break Poisson dependency so that Δvariance ≠ Δmean.

Evolutionary theories dating to the 1960s (*8-10*) proposed that biological organisms maximized fitness by harnessing putative fluctuations to enable probabilistic ‘bet-hedging’ decisions. Subsequent studies showed that intrinsic molecular fluctuations in gene expression (i.e., stochastic ‘noise’), modulated by gene-regulatory circuits, enabled probabilistic fate selection (Fig. 1A) in diverse biological systems (*11-13*). Open questions remain as to whether cellular noise control is limited to inherently locus-specific gene-regulatory circuits or if generalized noise-modulation mechanisms exist, if and how such mechanisms might orthogonally tune noise independent of mean, and, given the detrimental effects of noise, if such putative mechanisms might be regulated ‘on-demand’ to potentiate cell-fate specification.

**Figure 1:**
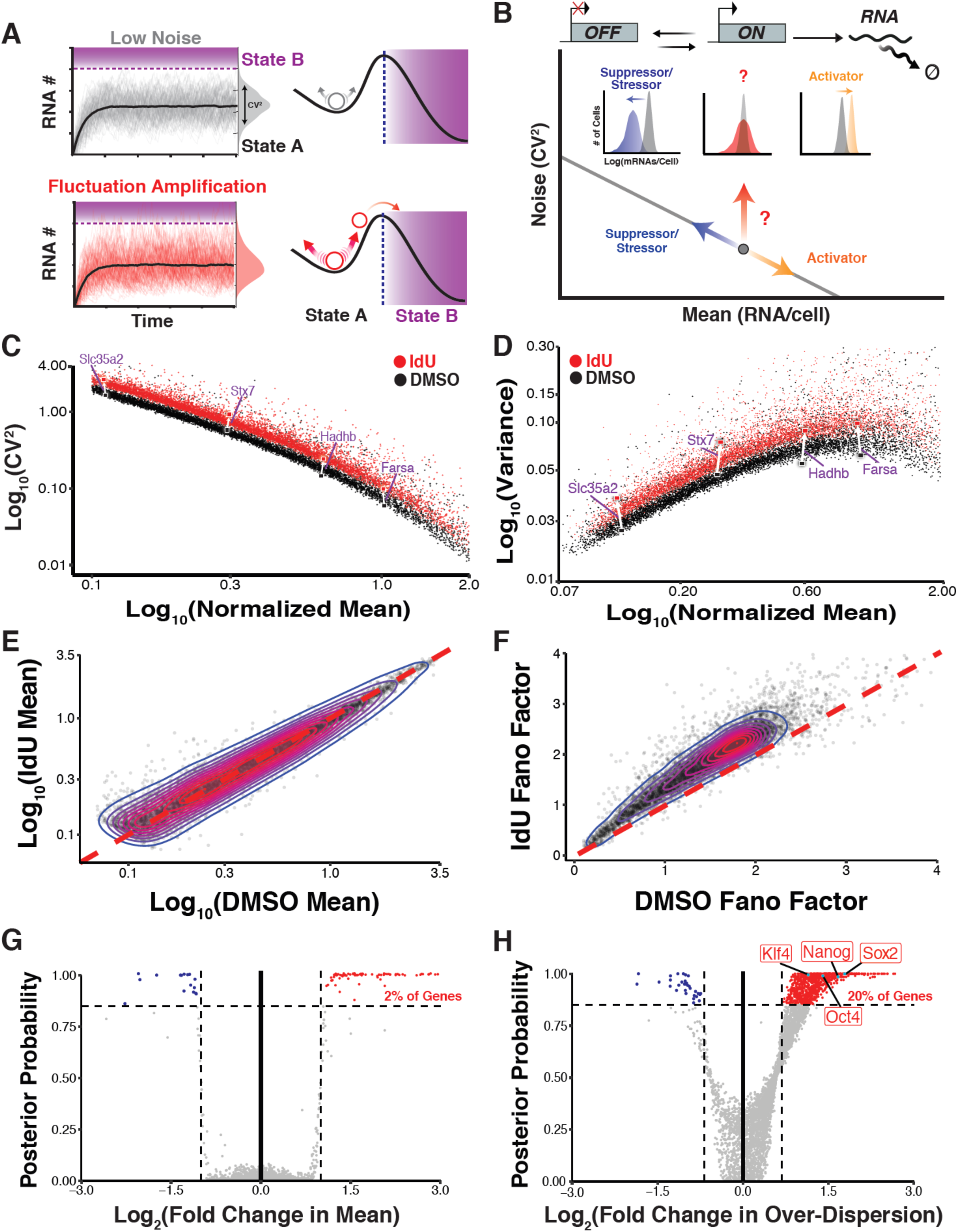
Genome-wide amplification of cell-to-cell mRNA variability (i.e., ‘noise’) independent of mean. **A**. (Left) Monte-Carlo simulations of the two-state Random-Telegraph model of transcription showing low noise and higher noise trajectories with matched mean expression levels. Coefficient of Variation (σ^2^/*μ*^2^, CV^2^) quantifies magnitude of fluctuations. (Right) The predicted facilitation of state transitions through ‘dithering’. **B**. (Top) Schematic of two-state Random-Telegraph model of transcription. (Bottom) Schematic of mean vs. CV^2^ for mRNA abundance with solid gray line representing Poisson, inverse scaling of CV^2^ as a function of mean. Question mark symbolizes unknown noise-control mechanisms that amplify fluctuations independently of mean. Histograms depict expected shift in mRNA copy number distributions. **C-F**. scRNA-seq of mESCs treated with DMSO (black) or 10µM IdU (red) for 24h. 812 and 744 transcriptomes (filtered and normalized with Seurat) from DMSO and IdU treatments, respectively, were analyzed. (C) Mean expression vs. CV^2^ and (D) mean vs. variance for 4,578 genes. Four examples of housekeeping genes (purple) demonstrate how IdU increases expression fluctuations with minimal change in mean (white arrows). (E) Mean expression and (F) Fano factor (σ^2^/*μ*) of 4,578 genes in DMSO vs. IdU treatments. Overlay of density contours reveals how center of mass lies on diagonal for mean values while lying above the diagonal for Fano factor measurements. **G-H**. BASiCS analysis of scRNA-seq data for 4,578 genes. (G) Fold change in mean vs. certainty (posterior probability) that gene is up- or down-regulated. With IdU treatment, 113 genes (red) were classified as differentially expressed (>2-fold change in mean with >85% probability). (H) Fold change in over-dispersion vs. certainty (posterior probability) that gene is highly- or lowly-variable. 945 genes (red) were classified as highly variable (>1.5-fold change in over-dispersion with >85% probability).

Non-genetic variability or noise in gene expression, often quantified by measurement of cell-to-cell variability in reporter expression, can arise from multiple sources, both intrinsic and extrinsic. In mammalian cells, intrinsic noise originates from episodic transcriptional ‘bursts’ (*14-17*) initiated by promoter toggling between ON and OFF states (Fig. 1B). The two-state random-telegraph model describes this bursting via two parameters: (i) the fraction of time a promoter is active (*K*_*ON*_*/[K*_*ON*_*+K*_*OFF*_]), and (ii) the number of transcripts produced during the ON state (burst size, *K*_*TX*_/*K*_*OFF*_) (*18-20*). These bursting parameters are tuned by regulatory machinery (*21*) like histone acetyltransferases, which can increase burst frequency by facilitating nucleosome clearance from promoters thereby increasing *mean* transcriptional levels (*22*). Increases in mean expression (μ) are typically accompanied by a stereotypical reduction in noise measured by coefficient of variation, CV, (σ/μ) (Fig. 1B), whereas stressors that decrease mean are typically accompanied by an increase in noise (*23-25*). This 1/μ scaling of noise can be broken by gene-regulatory circuits such as feedback and feedforward loops (*26*), and some small-molecule pharmaceuticals can modulate transcriptional fluctuations/noise (σ/μ) independent of change in mean (μ) (*27, 28*). Since some molecules can amplify expression noise of diverse unrelated promoters (*27, 29*), we asked if these molecules might be functioning via disruption or enhancement of a putative cellular noise-control mechanism.

A series of screens (Fig. S1) identified one compound, 5’-iodo-2’-deoxyuridine (IdU), which consistently increased expression noise of multiple transcriptional reporter constructs in diverse cell types. To test the generality of this noise amplification effect we focused on mouse embryonic stem cells (mESCs), due to extensive characterization of their developmental and transcriptional characteristics (*30-34*). Strikingly, single-cell RNA sequencing (scRNA-seq) of mESCs maintained in 2i/LIF media—after filtering and normalization using Seurat (*35*)—showed that IdU amplified cell-to-cell variability in transcript levels (i.e., transcript noise) for virtually all genes across the genome—4,578 genes analyzed (Fig. 1C)—with little alteration in mean transcript levels for most genes, as analyzed by either CV^2^ (σ^2^/µ^2^) or variance (σ^2^) versus mean (Figs. 1C– D). To account for the Poisson scaling of variance on mean, transcript noise was also quantified using the Fano factor (σ^2^/μ), which measures how noise deviates from Poisson scaling (σ^2^/μ = 1) (*20, 36, 37*). Despite mean-expression levels exhibiting minimal changes (Fig. 1E), the Fano factor increased for > 90% of genes (Fig. 1F) with lowly expressed genes showing a slightly greater change in Fano (Fig. S2). These results of a global increase in transcript noise with little change in mean levels are in stark contrast to the effects of transcriptional activators or cellular stressors that alter noise in a stereotypic manner together with changes in mean (*23, 38*).

To account for technical noise and quantify statistical significance of changes in noise and mean, we used an established Bayesian hierarchical model (*39, 40*) to create probabilistic, gene-specific estimates of both mean expression and cell-to-cell transcript variability. Of the 4,578 genes, the algorithm classified 945 genes (∼20%) as highly variable, whereas 113 genes (∼2%) showed a significant change in mean expression (Figs. 1G–H). Bulk RNA-seq measurements of mean abundances—performed using ERCC spike-ins for normalization—confirmed the scRNA-seq findings (Fig. S3). Thus, analyses from two methods (Seurat and BASiCS) show that IdU induces a significant increase in transcript variability (expression noise) but comparatively little change in mean expression in mESCs.

To examine if certain characteristics could explain a gene’s potential for noise enhancement we examined (i) gene length, (ii) promoter and (iii) gene-body AT content, (iv) number of exons, (v) TATA-box inclusion and (vi) strand orientation. None of these characteristics exhibited predictive power or correlated with a gene’s potential for noise enhancement (Fig. S4). However, genes susceptible to high noise enhancement were preferentially located within the interior of topologically associated domains (TADs), suggesting gene topology influences potency of noise enhancement (Fig. S4). Ontology analysis of highly variable genes showed enrichment of house-keeping pathways along with pluripotency maintenance factors, particularly Sox2, Oct4, Nanog and Klf4 (Figs. 1H, S5). As these pluripotency maintenance factors are key influencers of cell-fate specification, we next focused on the molecular mechanisms driving their amplified transcript noise.

We first tested whether the enhanced variability arose from extrinsic factors, which include cell-cycle phase and cell-type identity (*16, 41-44*). Cells within the scRNA-seq dataset were computationally assigned a cycle stage (G1, S, G2/M) (*45*) which showed that Nanog, Oct4, Sox2 and Klf4 were highly variable in each cell-cycle phase, indicating that their variability is not cell-cycle dependent (Fig. S6). Moreover, pseudo-time analysis showed no bifurcations, indicating transcriptional variability was not due to a differentiation-induced mixture of cell-types (Fig. S7).

Extrinsic variability may also arise from the coordinated propagation of noise through gene-regulatory networks (*46, 47*) and can be measured by gene-to-gene correlation matrices (*48, 49*). If the increase in global transcript noise is extrinsic, expression correlation between network partners would increase or remain unchanged. Analysis of gene-to-gene correlation matrices showed that ∼80% of gene-gene pairs *lost* correlation strength following IdU treatment (Figs. 2A and S8), indicating that enhanced expression noise is uncorrelated and not consistent with an extrinsic noise source. Exclusion of these extrinsic noise sources suggested that IdU amplifies intrinsic noise arising from stochastic fluctuations in transcript birth (promoter toggling) or death (degradation).

**Figure 2:**
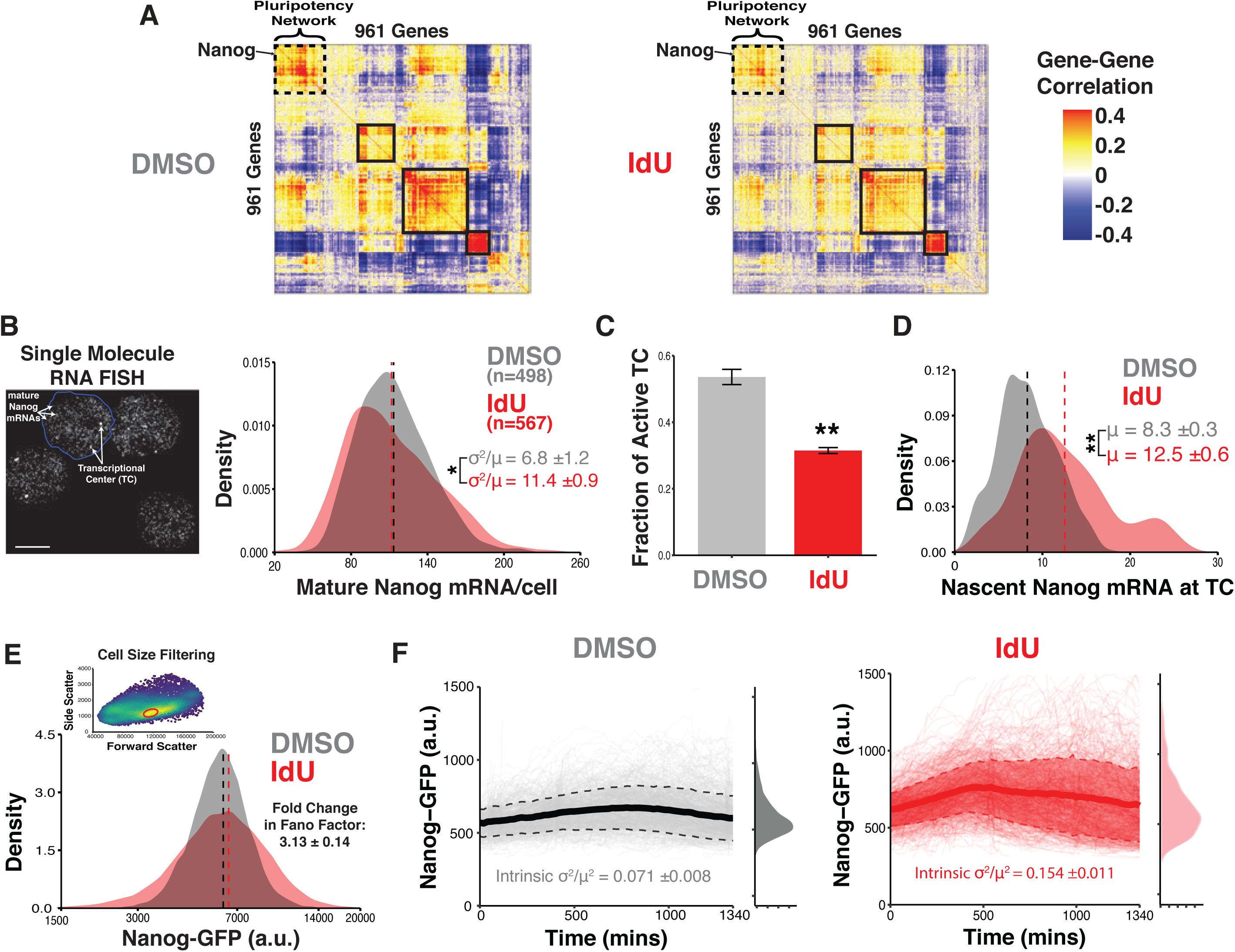
Amplification of mRNA noise is not due to extrinsic sources, results from shorter but more intense transcriptional bursts, and propagates to protein levels. **A**. Pearson correlations of expression for gene pairs in scRNA-seq dataset. Hierarchical clustering reveals networks of genes (highlighted in black rectangles) sharing similar correlation patterns. Dashed rectangle highlights network enriched with pluripotency factors like Nanog. IdU treatment causes a fading of heatmap, indicating weakened expression correlations. **B-D**. Results of smRNA-FISH used to count nascent and mature Nanog mRNA in Nanog-GFP mESCs treated with DMSO or 10µM IdU for 24 hours in 2i/LIF media. Data are from four biological replicates. (B) (Left) Representative micrograph (maximum intensity projection) in which Nanog transcripts are labelled with probe-set for eGFP. Bright foci correspond to transcriptional centers as verified by intron probe set. Scale bar is 5µm. (Right) Distributions of mature Nanog transcripts/cell. Dashed lines represent mean. IdU treatment increases cell-to-cell variability of transcript abundance as reported by averaged Fano factors (± SD), ^*^p =0.0011 by a two-tailed, unpaired Student’s t test. (C) Fraction of possible transcriptional centers that are active as detected by overlap of signal in exon and intron probe channels. Each cell is assumed to have 2 possible transcriptional centers (TCs). Data represent mean and SD. With IdU, the fraction of possible of TCs that are active decreases, ^**^p =6.9 x 10^−5^ by a two-tailed, unpaired Student’s t test. (D) Distributions of nascent Nanog mRNA per TC. With IdU, active TCs have more nascent mRNAs, ^**^p =1.0 x 10^−4^ by a two-tailed, unpaired Student’s t test. **E**. Representative flow cytometry distribution of Nanog-GFP expression in mESCs treated with DMSO or 10µM IdU for 24h in 2i/LIF. Dashed lines represent mean. Fold change in Fano factor (± SD) obtained from three biological replicates. IdU increases cell-to-cell variability in Nanog protein expression. Inset: Representative flow cytometry dot-plot showing conservative gating on forward and side scatter to filter extrinsic noise arising from cell size heterogeneity. **F**. Time-lapse imaging of Nanog-GFP mESCs treated with either DMSO (n = 1513) or 10µM IdU (n = 1414) in 2i/LIF. Image acquisition began immediately after addition of compounds. Trajectories from two replicates of each condition are pooled, with solid and dashed lines representing mean and standard deviation of trajectories respectively. Distributions of Nanog-GFP represent expression at final time-point. Intrinsic-CV^2^ of each detrended trajectory was calculated, with the average (± SD) of all trajectories reported.

To test whether a change in promoter toggling could account for IdU-enhanced noise, we used single-molecule RNA FISH (smRNA-FISH) to count both nascent and mature transcripts of Nanog, a master regulator of pluripotency. Spot counting was performed on a mESC line in which both endogenous alleles of Nanog are fused to eGFP. This fusion does not alter mRNA or protein half-life or impair differentiation potential (*50*). To target mature transcripts, smRNA-FISH probes to eGFP were used, and to minimize extrinsic noise, analyses were limited to cells of similar sizes (Fig. S9A). Consistent with scRNA-seq, smRNA-FISH showed a large increase in cell-to-cell variability of mature Nanog transcripts (∼2-fold increase in Fano) with little change in mean Nanog levels (Fig. 2B). Fewer IdU-treated cells exhibited active transcriptional centers (TCs), but the number of nascent mRNAs at each TC increased (Figs. 2C–D). Fitting of the two-state random-telegraph model to smRNA-FISH data revealed that increased variability was due to a shortened burst duration (increased *k*_*OFF*_) and amplified transcription rate (higher *k*_*TX*_) (Fig. S9B, Table S2). These results represent direct validation of previous predictions (*20, 27*) that enhanced noise could arise from reciprocal changes in transcriptional burst duration (1/*k*_*OFF*_) and intensity (*k*_*TX*_).

To test if enhanced transcript variability transmitted to the protein level, we performed flow-cytometric analysis of Nanog-GFP reporter protein. In IdU-treated cells, the Nanog protein Fano factor increased by ∼3-fold, with little change in mean, indicating that mRNA variability from altered promoter toggling indeed resulted in changes to protein noise (Fig. 2E). The increase in protein noise showed no dependency on cell-cycle (Fig. S10C–D) despite G1-to-S cell-cycle progression being slightly slowed by IdU treatment (Fig. S10A–B). Consistent with the extrinsic noise analysis above, there was no evidence of aneuploidy following IdU treatment (Fig. S10A), precluding the possibility that increased noise results from a sub-population of cells with non-physiologic gene-copy numbers.

Given that Nanog noise was intrinsic and transmitted to the protein level, we next tested a previous theoretical prediction about Nanog. When cultured in 2i/LIF, mESCs exhibit Nanog protein expression that is unimodal and high, but when cultured in serum/LIF, mESCs exhibit bimodal Nanog expression with both a low Nanog state and a high Nanog state (Fig. S11) (*34*). Theories predicted that increased transcriptional noise would drive greater excursions from the high Nanog state into the low Nanog state (*51*). We found that in mESCs cultured in serum/LIF, IdU-induced amplification of Nanog noise did indeed generate greater excursions into the low Nanog state (Fig. S11), verifying theoretical predictions. This result demonstrates how promoter toggling can drive Nanog state-switching thereby altering differentiation potential.

To verify that enhanced noise is not a population-level phenomenon brought on by differential responses to IdU in distinct cellular subpopulations (i.e., verify ‘ergodicity’ and that individual cells exhibit increased fluctuations), we used live-cell time-lapse imaging to quantify both the magnitude (intrinsic-CV^2^) and frequency content (1/half-autocorrelation time) of Nanog fluctuations. Single-cell tracking of individual cells showed that IdU induced a 2-fold increase in the magnitude (intrinsic-CV^2^) of fluctuations (Figs. 2F, S12A), and auto-correlation analysis of detrended trajectories showed a broadening of the frequency distribution to higher spectra, indicating reduced memory of protein state (Fig. S12B). These higher frequency fluctuations are consistent with amplification of a non-genetic, intrinsic source of noise (*52, 53*) because genetic sources of cellular heterogeneity, such as promoter mutations, would lead to longer retention of protein states (increased memory) (*54*). *In silico* sorting of cells based on starting Nanog expression verified that noise enhancement was not dependent on memory of initial state (Fig. S13). Fluctuations in promoter toggling therefore drive individual cells to dynamically explore a larger state-space of Nanog expression. To further validate that IdU perturbs an intrinsic source of noise, we used a mESC line in which the two endogenous alleles of Sox2 are tagged with P2A-mClover and P2A-tdTomato, respectively, to enable quantification of the intrinsic and extrinsic components of noise. Treatment with IdU increased Sox2 intrinsic noise greater than 2-fold across all expression levels (Fig. S14) further validating that IdU enhances intrinsic noise.

To test if a generalized stress response could explain the noise enhancement induced by IdU, we subjected Nanog-GFP mESCs to UV radiation for 15, 30, or 60 minutes. Both the mean and Fano factor of Nanog expression decreased for all timepoints, which markedly differs from IdU treatment (Fig. S15). This result further indicates that IdU does not perturb an extrinsic or global noise source; rather, it perturbs an intrinsic source of noise (i.e., promoter toggling).

To pinpoint the molecular mechanism, 14 additional nucleoside analogs (Table S3) were screened for noise enhancement effects. 5’-bromo-2’-deoxyuridine (BrdU), 5-hydroxymethylcytosine (hmC), and 5-hydroxymethyluridine (hmU) also increased Nanog Fano factor to varying degrees (Fig. 3A). Intriguingly, hmU and hmC are naturally produced by the Ten Eleven Translocation (Tet) family of enzymes during oxidation of thymine and methylated cytosine respectively (*55-58*). Given that these base modifications are removed via base-excision repair (BER), we surmised that their incorporation and removal from genomic DNA may be responsible for noise enhancement (Fig. 3B) (*59, 60*). To test this, we knocked down 25 genes (3 gRNAs/gene, Table S4) involved in nucleoside metabolism and DNA repair using CRISPRi, and quantified how these knockdowns affected IdU’s noise enhancement. We identified 2 genes: AP Endonuclease 1 (Apex1) and thymidine kinase 1 (Tk1) whose knockdown abrogated noise enhancement (Fig. 3C). Knockdown was confirmed via RT-qPCR (Fig. S16).

**Figure 3:**
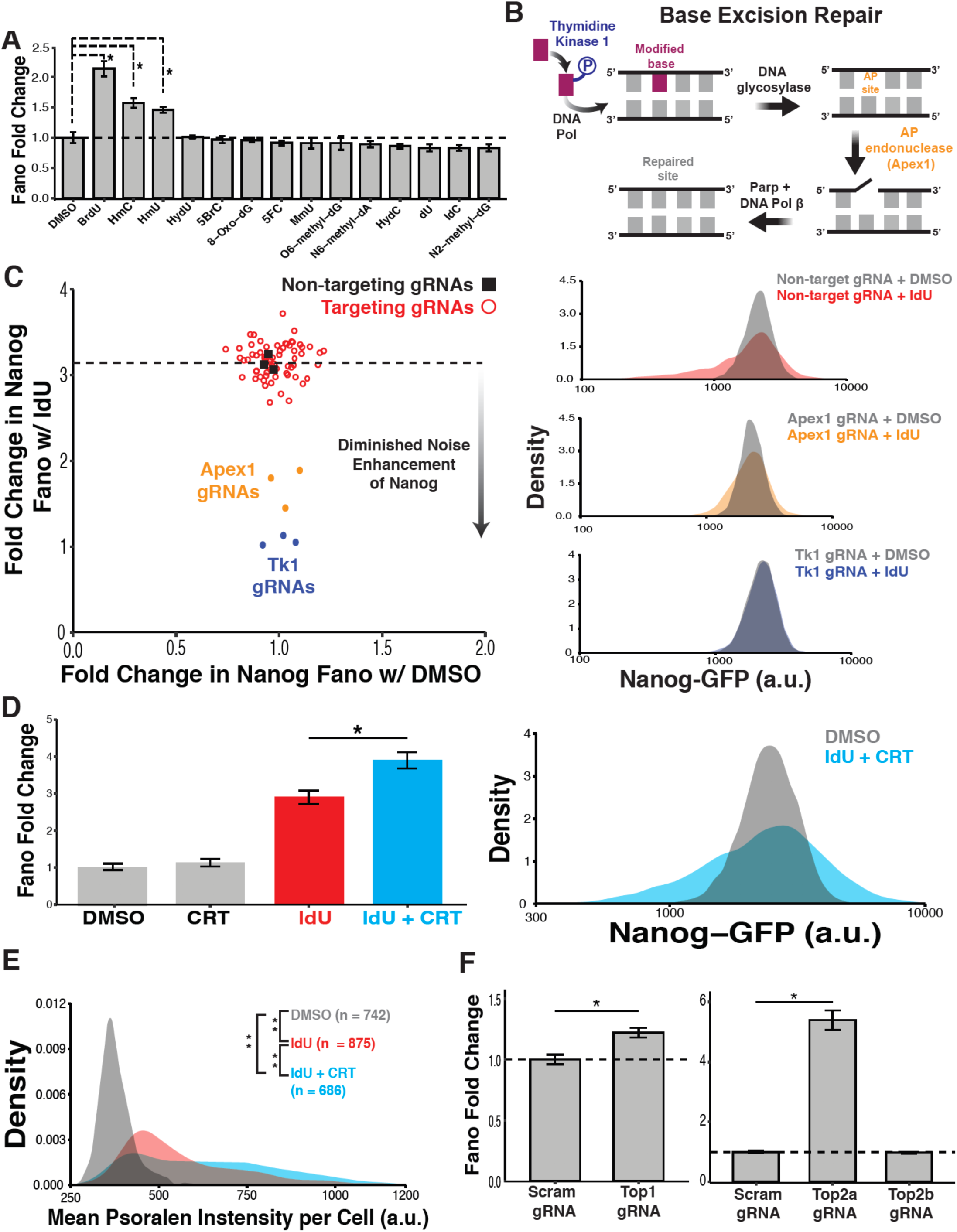
Noise amplification independent of mean is due to Apex1-mediated DNA repair. **A**. Screening of 14 additional nucleoside analogs. Nanog-GFP mESCs grown in 2i/LIF were supplemented with 10µM of nucleoside analog for 24h. Fano factor for Nanog protein expression was normalized to DMSO. Data represent mean (± SD) of two biological replicates. BrdU, hmC and hmU increase Nanog expression variability as compared to DMSO, ^*^p < 0.01 by a Kruskal-Wallis test followed by Tukey’s multiple comparison test. **B**. Schematic of nucleoside analog incorporation into genomic DNA and removal via base excision repair pathway. **C**. (Left) CRISPRi screening for genetic dependencies of IdU noise enhancement. Nanog-GFP mESCs stably expressing dCas9-KRAB-p2A-mCherry were transduced with a single gRNA expression vector with BFP reporter. 75 gRNAs (25 genes, 3 gRNAs/gene) were tested in addition to 3 non-targeting control gRNAs. Two days following transduction, each gRNA-expressing population of mESCs was treated with DMSO or 10µM IdU for 24h in 2i/LIF media. Nanog-GFP protein expression was measured for mCherry/BFP double positive cells. Nanog Fano factor for DMSO and IdU treatment of each gRNA population was normalized to Nanog Fano factor of non-targeting gRNA+DMSO population. Each point represents a gRNA. Dashed horizontal line represents average noise enhancement of Nanog from IdU in the background of non-targeting gRNA expression (black squares). Knockdown of Apex1 and Tk1 diminishes noise enhancement of Nanog from IdU. (Right) Representative flow cytometry distributions of Nanog expression for mESCs expressing non-targeting (top-right), Apex1 (middle-right), or Tk1 (bottom-right) gRNAs and treated with DMSO or 10µM IdU. **D**. Combination of IdU and small-molecule inhibitor of the Apex1 endonuclease domain (CRT0044876). (Left) mESCs were treated with DMSO, 100µM CRT0044876, 10µM IdU or 10µM IdU + 100µM CRT0044876 for 24h in 2i/LIF. Nanog Fano factor for each treatment was normalized to DMSO control. Data represent mean (± SD) of three biological replicates. Inhibition of Apex1 endonuclease domain in combination with IdU synergistically increases cell-to-cell variability of Nanog expression, ^*^p = 0.0028 by a two-tailed, unpaired Student’s t test. (Right) Representative flow cytometry distributions of Nanog expression for mESCs treated with DMSO or 10µM IdU + 100µM CRT0044876. **E**. Single-cell quantification of negative supercoiling levels using psoralen-crosslinking assay. mESCs were treated with DMSO, 10µM IdU or 10µM + 100µM CRT0044876 for 24h in 2i/LIF. 1µM aphidicolin was added to cultures 2h prior to incubation with biotinylated-trimethylpsoralen (bTMP). Following UV-crosslinking, cells were stained with streptavidin-Alexa594 conjugate and DAPI. Distributions for nuclear intensities of bTMP staining are shown. Data are pooled from two biological replicates of each treatment. IdU treatment increases negative supercoiling as compared to DMSO control, ^**^p < 0.0001. IdU in combination with CRT0044876 further increases supercoiling levels as compared to DMSO (^**^p < 0.0001) and IdU alone (^**^p < 0.0001). P values were calculated using Kruskal-Wallis test followed by Tukey’s multiple comparison test. **F**. CRISPRi Knockdown of Topoisomerases involved in relaxation of DNA supercoiling. Nanog Fano factor was normalized to scrambled gRNA population. Data represent mean (± SD) of three biological replicates. Knockdown of Top1 (*p = 0.002) and Top2a (*p = 0.003) increases Nanog expression variability. P values were calculated by two-tailed, unpaired Student’s t test.

Tk1 adds a requisite gamma-phosphate group to diphosphate nucleotides prior to genomic incorporation (Fig. 3B) (*61*). Our knockdown results indicated that phosphorylation of IdU by Tk1 and subsequent incorporation of phosphorylated IdU into the genome may be necessary for noise enhancement. To validate this, we tested the effect of treatment with 10µM IdU combined with excess thymidine, a competitive substrate of Tk1. Competitive inclusion of thymidine returned Nanog noise to baseline levels (Fig. S17), indicating that noise enhancement is dependent on IdU incorporation. The *reduction* in Nanog noise with addition of exogenous thymidine also suggests that IdU-induced noise amplification is not a generic effect of nucleotide imbalances within the cell.

Apex1 (a.k.a., Ref-1, Ape1) plays a pivotal role in the BER pathway as it incises DNA at apurinic/apyrimidinic sites via an endonuclease domain, allowing for subsequent removal of the sugar backbone and patching of the gap (*62, 63*). To confirm the knockdown results, we attempted to knockout Apex1 in mESCs. However, the knockout was lethal, in agreement with previous reports (*64*). As an alternative, we used a small-molecule inhibitor (CRT0044876) specific for the Apex1 endonuclease domain (*65*). Unexpectedly, the combination of CRT0044876 with IdU synergistically increased Nanog protein variability, without significantly changing the mean (Fig. 3D).

The contrasting effects of Apex1 knockdown and catalytic inhibition implied that a physical rather than enzymatic quality of the protein is responsible for modulation of transcriptional bursting. In support of this, Apex1 induces helical distortions and local supercoiling to identify mismatched bases (*66, 67*). Furthermore, catalytically inactive Apex1 binds DNA with higher affinity (*68*). Therefore, CRT0044876 may lengthen Apex1 residence times on DNA, thus synergistically amplifying topological reformations. Taken together with evidence that supercoiling sets mechanical bounds on transcriptional bursting (*69-71*), we next asked whether Apex1 recruitment impacts supercoiling levels.

To assay supercoiling, we used a psoralen-crosslinking assay in which mESCs are incubated with biotinylated-trimethylpsoralen (bTMP), which preferentially intercalates into negatively supercoiled DNA (*72, 73*). To eliminate DNA replication as a contributor of supercoiling, aphidicolin is added to inhibit DNA polymerases prior to bTMP incubation (*74*). IdU treatment significantly increased genomic supercoiling as demonstrated by a ∼2-fold increase in bTMP intercalation (Fig. 3E). The combination of IdU and CRT0044876 further increased intercalation, suggesting that supercoiling levels are correlated with noise enhancement through increased Apex1-DNA interactions (Fig. 3E). IdU treatment followed by a short incubation with bleomycin (which decreases supercoiling through double-stranded breaks) reduced bTMP intercalation below the DMSO control level, indicating IdU alone in uncoiled DNA does not increase intercalation (Fig. S18).

If DNA topology influences transcriptional bursting, additional modifiers of supercoiling should also affect Nanog noise. Topoisomerase 1 and 2a (Top1 and Top2a, respectively) relax coiled DNA through the introduction of single- and double-stranded breaks, respectively. Knockdown of Top1 and Top2a via CRISPRi increased Nanog protein variability (Fig. 3F). Furthermore, inhibition of topoisomerase activity with the small-molecule inhibitors topotecan and etoposide recapitulated these effects (Fig. S19). Taken together with psoralen-crosslinking data, these results suggest that Apex1-induced supercoiling tunes gene-expression fluctuations without altering mean expression levels.

To understand the mechanism by which Apex1 might increase transcriptional noise without altering mean expression levels, we developed a series of minimalist computational models to account for the experimental data (supplementary text 2). We ran Monte Carlo simulations of each model using smRNA-FISH data for parameterization (Table S2); this revealed that a transcription-coupled-repair (TCR) mechanism best accounts for the noise-without-mean amplification (Figs. 4A-B, S20-21, supplementary text 5). Importantly, Apex1 binding triggers a transcriptionally non-productive, negatively supercoiled state, whereas unbinding of Apex1 allows mRNA production to resume with an amplified transcription rate. This amplification may originate from the increased negative supercoiling during repair which can facilitate upstream binding of transcriptional machinery in preparation for when repair is complete (*75-82*). The ability to render a gene transcriptionally non-productive while also stimulating recruitment of transcriptional resources points to a homeostatic mechanism: the BER pathway maintains gene-expression homeostasis (i.e., mean) by amplifying transcriptional fluctuations through reciprocal modulation of burst intensity and duration (Fig. 4B).

**Figure 4:**
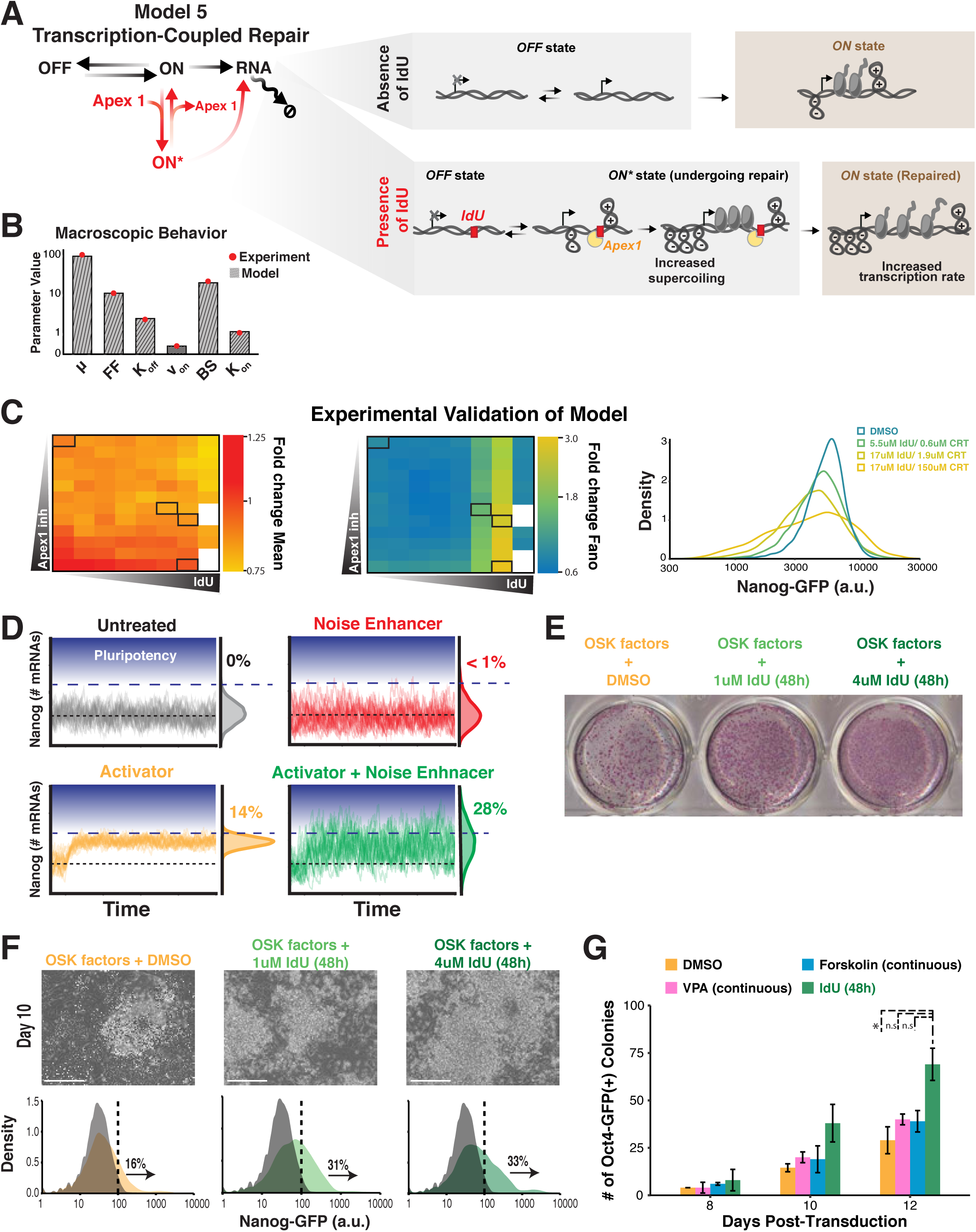
Transcription-coupled DNA repair tunes transcriptional bursting across a broad parameter regime, generating amplified noise which potentiates reprogramming of cellular identity. **A**. Detailed schematic of Model 5 (see Fig. S20 for schematic of models 1-4), which represents transcription-coupled repair (TCR). In the presence of IdU (bottom panel), Apex1 binding occurs when gene is transcriptionally permissive (ON state). Binding induces negative supercoiling which lengthens the time that a gene is transcriptionally non-productive (ON* state) while also facilitating recruitment of transcriptional resources. Upon repair completion, a higher transcriptional rate compensates for lost productivity. Mean expression is maintained with larger transcriptional fluctuations. **B**. The macroscopic behavior (mean Nanog mRNA [µ], Fano factor [FF], *K*_*off*_, fraction of time active [v_on_], burst size [BS], *K*_*on*_) of model 5 simulations are compared to experimentally derived values of each parameter (red dots) obtained from smRNA-FISH data. Absolute percentage error (APE) is calculated as described in supplementary text 5.2.2. Model 5 (TCR model) best matches experimental data. **C**. Testing of 96 concentration combinations of IdU and CRT0044876 to validate tunability of Nanog variability. IdU and CRT0044876 were used to increase binding and decrease unbinding of Apex1 respectively. Nanog-GFP mESCs grown in 96-well plates were treated with 12 concentrations of CRT0044876 ranging from 0 to 150µM in combination with 8 concentrations of IdU ranging from 0 to 50µM. Data represent average of two biological replicates. (Leftmost and Center Panels) 96-well heatmaps displaying fold change in Nanog mean and Fano factor for each drug combination as compared to DMSO (top-leftmost well). Insufficient number of cells (<50,000) for extrinsic noise filtering were recorded from white wells. (Rightmost Panel) Representative flow cytometry distributions from highlighted wells (black rectangles). Nanog variability increases independently of the mean. **D**. Simulations of the TCR model for Nanog gene expression in the presence of DMSO (top left), IdU (top right), an activator (increased *K*_*ON*,_ decreased *K*_*OFF*_) of promoter activity (bottom left) and an activator combined with IdU (bottom right). Homeostatic noise amplification potentiates responsiveness to an activator of gene expression as demonstrated by increased threshold crossing (14% to 28%). **E**. Nanog-GFP secondary MEFs (seeded at 10,000 cells/cm^2^) harboring stably-integrated, doxycycline-inducible cassettes for Oct4, Sox2, and Klf4 (OSK) were subjected to 10 days of doxycycline treatment in combination with DMSO (first well), 1µM IdU (second well), or 4µM IdU (third well) for the first 48 hours of reprogramming. Alkaline phosphatase staining for pluripotent colonies of cells demonstrates how IdU treatment potentiates pluripotency induction. **F**. (Top) Micrographs of Nanog-GFP secondary MEFs at day 10 of doxycycline-induced reprogramming (scale bar = 100 µm). (Bottom) Flow cytometric analysis of Nanog-GFP activation at day 10 of reprogramming. Data are pooled from two replicates. **G**. Oct4-GFP primary MEFs (seeded at 10,000 cells/cm^2^) were retrovirally transduced with cDNAs encoding Oct4, Sox2, Klf4, and c-Myc. 24 hours after transduction, infected cells were treated with DMSO (continuously), 1mM valproic acid (VPA, continuously), 10µM forskolin (continuously), or 4µM IdU (first 48 hours). VPA and forskolin are established enhancers of cellular reprogramming. The number of Oct4-GFP(+) stem cell colonies were counted 8, 10, and 12 days from the start of drug treatment. Data represent mean and SD of 2 biological replicates. Treatment of transduced MEFs with IdU during early stages of reprogramming increases the number of Oct4-GFP(+) colonies that form as compared to DMSO control, *p =0.039 by one-way ANOVA with Bonferroni post hoc test.

Sensitivity analysis of the Apex1 TCR model revealed that orthogonal modulation of Nanog mean and noise is possible within a large portion of the parameter space (Fig. S22A-B, supplementary text 6). As validation, we tested the effect of 96 concentration combinations (Table S6) of IdU and CRT0044876 to perturb the rates of Apex1 binding and unbinding respectively. The experimental results confirmed model predictions, showing that Nanog noise could be tuned independently of the mean (Fig. 4C). Testing of BrdU and hmU further validated that parameter regimes exist where noise can be regulated independent of mean (Fig. S23). The hmU data in particular showed that the BER pathway can amplify noise while maintaining mean expression when removing a naturally occurring base modification. Additionally, sensitivity analysis indicated that for genes whose *K*_*OFF*_ >> *K*_*ON*_ (i.e., lowly expressed genes), IdU treatment would increase mean abundance (Fig 22E). This prediction was verified experimentally with bulk RNA-seq measurements of transcript abundance in mESCs treated with IdU (Fig S3), as all 98 of the up-regulated genes reside within the lowest expression regime.

To test whether the homeostatic mechanism of BER applies to additional genes, mRNA distributions from the scRNA-seq dataset were fit to a Poisson-beta model (two-state model) allowing for estimation of *K*_*ON*_, *K*_*OFF*_, and *K*_*TX*_ (*83, 84*). A consistent pattern emerged for genes classified as highly variable: 80% exhibited increased rates of promoter inactivation (*K*_*OFF*_) and 84% had increased transcription rates (*K*_*TX*_) (Fig. S24). Alignment of these rate estimates with predictions from the TCR model revealed that the BER pathway can alter noise while maintaining transcriptional homeostasis by individually tailoring expression fluctuations for genes with vastly different bursting kinetics (Fig. S25, supplementary text 7).

We next asked if this amplification of transcriptional variability acted to enhance cellular plasticity as previously suggested (*85*). Using a neural-network approach, we reconstructed the Waddington landscape based on a predictive model of gene-gene interactions inferred from the scRNA-seq data (*86*). In this approach, each cell has a characteristic energy determined by its proximity to an attractor state, with lower energy values corresponding to greater stability. The analysis indicated that cells exposed to IdU lie at a higher altitude on this landscape, indicating destabilization of cellular identity and greater developmental plasticity (Fig. S26). Numerical simulations of the TCR model then verified that IdU-mediated amplification of transcriptional noise has the potential to increase responsiveness to activation stimuli (Fig. 4D). The complementary abilities of IdU-mediated noise amplification to destabilize cellular identity and potentiate responsiveness to fate signals suggested that IdU might facilitate cellular reprogramming.

To experimentally verify these predictions, we tested if IdU could potentiate conversion of differentiated cells into pluripotent stem cells using two cellular reprogramming systems. The first assay utilized mouse embryonic fibroblasts (MEFs) that express GFP from the endogenous Nanog locus and harbor stably integrated, doxycycline-inducible cassettes for three of the Yamanaka factors: Oct4, Sox2, and Klf4 (OSK). As confirmation that IdU acts as a noise-enhancer in this system, treatment of secondary MEFs with IdU for 48 hours in standard MEF media caused increased variability in Nanog protein expression (Fig. S27A) with no changes in cell-cycle progression (Fig. S27B). Strikingly, IdU supplementation for the first 48 hours of a 10-day reprogramming course enhanced the formation of pluripotent colonies as measured by alkaline phosphatase staining (Fig. 4E). Bulk RNA-seq at days 2 and 5 of reprogramming (Fig. S27C) and flow-cytometric analysis at day 10 (Fig. 4F) demonstrate that early-stage noise-enhancement accelerates activation of the pluripotency program. To confirm the results in an orthogonal reprogramming assay, Oct4-GFP primary MEFs were transduced with retroviral vectors expressing Oct4, Sox2, Klf4, and c-Myc. IdU supplementation for the 48 hours immediately following transduction caused a ∼2.4-fold increase in the number of Oct4-GFP(+) colonies (Fig. 4G), further demonstrating how amplification of intrinsic gene expression fluctuations can potentiate cell-fate conversion.

Overall, these data reveal that a DNA-surveillance pathway exploits the biomechanical link between supercoiling and transcription to homeostatically enhance noise without altering mean-expression levels. This homeostatic noise-without-mean amplification appears to increase cellular plasticity, thus facilitating reprogramming of cellular identity. This raises intriguing implications for the role of naturally occurring oxidized nucleobases (e.g., hmU) in cell-fate determination, particularly since these base modifications are found at higher frequencies in embryonic stem-cell DNA (*57*). Mechanistic insight from modeling and experimental perturbation of Apex1 suggest that homeostatic (i.e., orthogonal) noise amplification may also apply to other DNA-processing activities that interrupt transcription. It is important to note that homeostatic noise amplification cannot occur for all promoters (i.e., promoters with *K*_*OFF*_ >> *K*_*ON*_ are precluded as they will exhibit increased mean) and propagation of transcriptional variability to the protein level likely depends on protein half-lives and thus may not occur for a large swath of proteins. The proteins monitored in this study either have naturally short half-lives (Nanog) or PEST tags (e.g. d_2_GFP) which minimizes the buffering of transcriptional bursts conferred by longer protein half-lives (*87*). The ability to independently control the mean and variance of gene expression may indicate that cells have the ability to amplify transcriptional noise for fate exploration and specification.

## Acknowledgements

We thank Michael Simpson, Benoit Bruneau, Jonathon Weissman and the Weinberger lab for thoughtful discussions and suggestions. We thank Kathryn Claiborn for editing and Giovanni Maki for graphics support. We would like to acknowledge the technical assistance of Nandhini Raman in the Gladstone Institute Flow Cytometry Facility (NIH S10 RR028962, P30 AI027763, DARPA, and the James B. Pendleton Charitable Trust) and the Gladstone Assay Development and Drug Discovery Core. We also acknowledge Kurt Thorn and DeLaine Larson in the UCSF Nikon Imaging Center (NIH S10 1S10OD017993-01A1). We are grateful to the Gladstone Institute Genomics Core for assistance with single-cell RNA-sequencing experiments. The dual-tagged Sox2 mESCs were a kind donation from Benoit Bruneau and Elphege Nora. We thank Noam Vardi for the EF-1*α* d_2_GFP and UBC d_2_GFP isoclones. We thank Marco Jost and Jonathon Weissman for CRISPRi reagents.

## Funding

R.V.D. is supported by an NIH/NICHD F30 fellowship (HD095614-03). L.S.W. acknowledges support from the Bowes Distinguished Professorship, Alfred P. Sloan Research Fellowship, Pew Scholars in the Biomedical Sciences Program, NIH awards R01AI109593, P01AI090935, and the NIH Director’s New Innovator Award (OD006677) and Pioneer Award (OD17181) programs.

## Author contributions

R.V.D. and L.S.W. conceived and designed the study. R.V.D., C.U., S.D., and L.S.W conceived and designed the cellular reprogramming experiments. R.V.D. and C.U. performed the experiments. R.V.D., M.M.K.H., and B.M. analyzed data. R.V.D., M.M.K.H., B.M. and L.S.W. constructed and analyzed the mathematical models. R.V.D. and L.S.W. wrote the manuscript.

## Competing interests

Authors declare no competing interests.

## Data and materials availability

Sequencing data from bulk RNA-seq and single-cell RNA-seq will be deposited onto GEO. Custom code for analysis of sequencing data and mathematical modeling will be made available on GitHub. Reagents, including plasmids and cell-lines, are available from the corresponding author upon request.

## Supplementary Materials

Materials and Methods

Supplementary Text

Figures S1-S27

Tables S1-S6

References (*88-96*)

## Supplementary Materials for

## This PDF file includes

Materials and Methods

Supplementary Text

Figs. S1 to S27

Captions for Tables S1 to S6

## Materials and Methods

### Cell Culture and Growth Conditions

Mouse E14 embryonic stem cells (male) were routinely cultured in feeder-free conditions on gelatin-coated plates with ESGRO-2i medium (Millipore, cat:SF016-200) at 37°C, 5% CO_2_, in humidified conditions *(36)*. Jurkat T Lymphocytes (male) were cultured in RPMI-1640 medium (supplemented with L-glutamine, 10% fetal bovine serum, and 1% penicillin-streptomycin), at 37°C, 5% CO_2_, in humidified conditions at 0.1 × 10^6^ to 1 × 10^6^ cells/mL. Isoclonal Jurkat T Lymphocytes with lentivirally integrated EF1α-d_2_GFP construct were previously described *(36)*. Human immortalized myelogenous leukemia (K652, female) cells were cultured in RPMI-1640 medium (supplemented with L-glutamine, 10% fetal bovine serum, and 1% penicillin-streptomycin), at 37°C, 5% CO_2_, in humidified conditions at 2 × 10^5^ to 2 × 10^6^ cells/mL. Isoclonal K562 cells with lentivirally integrated EF1α-d_2_GFP and UBC-d_2_GFP constructs were previously described *(36)*. Human embryonic kidney (HEK293, female) cells were cultured in DMEM (supplemented with 10% fetal bovine serum, 1% penicillin-streptomycin, 25mM HEPES, and 2mM L-glutamine) at 37°C, 5% CO_2_, in humidified conditions at 30 to 90% confluency.

### Noise Enhancer Testing on Isoclonal Jurkat and K562 Cells

Jurkat and K562 cells were seeded into 12-well plates at densities of 0.2 × 10^6^ and 0.4 × 10^6^ cells/mL respectively in media containing 20 µM IdU (Sigma, cat:I7125, dissolved in DMSO) or equivalent volume of DMSO for 24 hours. Flow cytometry was performed using a BD LSRII cytometer. Treated cells were run unfixed and live to avoid additional sources of variability from fixation. 50k live cells were collected from each sample for noise measurements. Conservative gating for a live subset of approximately 3k cells of similar size, volume, and state, was applied on the FSC vs. SSC to reduce extrinsic noise contributions as previously described *(25,27)*.

### Single-Cell RNA Sequencing Preparation and Analysis

1×10^6^ mESCs were seeded in a gelatin-coated, 10cm dish in 2i/LIF media. 24 hours following seeding, cultures were replenished with 2i/LIF media containing 10µM IdU or an equivalent volume of DMSO for 24 hours. After treatment, cells were trypsinized with TrypLE and spun down for 5 minutes at 200 x g. Single-cell suspensions were prepared in DPBS at a concentration of 83,000 cells/ml. Approximately 3000 cells from each sample were loaded into a chip and processed with the Chromium Single Cell Controller (10x Genomics). To generate single-cell gel beads in emulsion (GEMs), DMSO- and IdU-treated samples were assigned unique indexes using Single Cell 3 ′ Library and Gel Bead Kit V2 (10x Genomics, cat:120237). Sequencing was performed on an Illumina HiSeq4000 with a paired-end setup specific for 10x libraries.

Data were aligned to mm10 reference genome using 10x Cell Ranger v2. Quality control, normalization and analysis were carried out using two packages: Seurat and BASiCS. For analysis in Seurat, gene-barcode matrices were filtered and normalized using the “LogNormalize” method, resulting in 812 and 744 transcriptomes from DMSO and IdU samples. Transcript variability was quantified using variance (*σ* ^2^), coefficient of variation 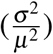, and Fano factor 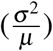. During the normalization procedure in Seurat, counts for the *i*^*th*^ gene in the *j*^*th*^ cell (*x*_*i j*_) are multiplied by the following scaling factor: 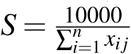, where n is the number of genes in the dataset. The scaling factor is therefore dependent on the number of UMIs detected per cell. The coefficient of variation is insensitive to this scaling factor as it is a dimensionless quantity (i.e, *σ* and *µ* are scaled by the same factor and thus cancel out when calculating coefficient of variation). However, the Fano factor, which has units, must be re-scaled to account for the differential effect that this normalization procedure has on *σ* ^2^ vs. *µ* (i.e., *σ* ^2^ gets scaled by *S*^2^ while *µ* gets scaled by S). To negate the carryover of this scaling factor, calculated Fano factors from the Seurat-normalized dataset were multiplied by 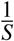 where S is a unique value for the DMSO and IdU samples: 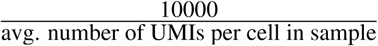. On average, 4151.3 and 4191.4 UMIs were detected per cell in DMSO and IdU samples respectively.

For analysis using BASiCS, quality control and filtering was performed using the BASiCS Filter function resulting in an identical number of transcriptomes (812 and 744) as produced by Seurat. Posterior estimates of mean and over-dispersion for each gene were computed using a Markov Chain Monte Carlo (MCMC) simulation with 40,000 iterations and a log-normal prior. For differential mean testing, a threshold of fold change *>*2 with an FDR cutoff of 0.05 was used. Differential variability was tested with a threshold of fold change *>*1.5 with an FDR cutoff of 0.05. Only genes with no change in mean expression (4,458 of 4,578) were considered for interpreting changes in variability.

Gene features and sequences from the GRCm38 reference were used for analysis of gene characteristics that potentiate noise enhancement. TAD boundary locations in mESCs were taken from Hi-C maps produced by Elphège et al *(89)*. DAVID v6.8 was used to test for gene ontology (GO) enrichment among highly variable genes. All tested genes (4,458) from BASiCS were used as background. Bonferroni-corrected p-values (adjusted p-values) were used to visualize GO enrichment. Cell cycle determination was performed using *cyclone* as implemented in scran *(45)*. The default set of cell cycle marker genes for mESCs (mouse cycle markers.rds) was used. Cells were assigned to G1, S, and G2/M phases using their normalized genes counts produced by Seurat. Pseudotime analysis was conducted using *destiny (90)* with the Seurat-normalized cell-gene matrix as input. Gene-gene correlation matrices were assembled by first filtering out genes from the Seurat-normalized matrix whose mean abundance *<*1 in each treatment group to avoid spurious correlations that may emerge from low expression. 961 genes remained for downstream analysis. Pearson correlation for each gene pair was calculated. Clustering of gene-pairs based on similarity in correlation patterns was performed using the hierarchical clustering method within the *seriation* package. Change in correlation strength was calculated by subtracting absolute value of gene-pair correlation in DMSO condition from IdU condition.

### Bulk RNA Sequencing Preparation and Analysis

2×10^5^ mESCs were seeded in each well of a gelatin-coated, 6-well plate in 2i/LIF media. 24 hours following seeding, cultures were replenished with 2i/LIF media containing 10µM IdU, 5µM IdU or an equivalent volume of DMSO in triplicate for 24 hours. After treatment, cells were trypsinized with TrypLE and RNA was extracted using a RNeasy minikit (Qiagen) according to manufacturer’s instructions. ERCC spike-in RNA (2µl diluted at 1:100) was added to each RNA extraction (Ambion, cat:4456740). A total of 9 cDNA libraries were prepared with an NEBNext Ultra II RNA Library Prep kit (NEB, cat:E7770S) and sequenced with an Illumina HiSeq4000. Sequencing yielded a median of ∼ 40 million single-end reads per library. Read quality was checked via FASTQC. Reads were aligned to an edited version of the mm10 reference genome containing the ERCC spike-in sequences using TopHat with default parameters. Transcript level quantification was performed using Cufflinks with default parameters. The quantification matrix was then imported into R and analyzed via DESeq2. Samples were normalized using ERCC transcripts as controls for size factor estimation. Differential mean testing was conducted with a threshold of fold change *>*2 and an FDR cutoff of 0.05.

### Single Molecule RNA FISH

Probes for detection of nascent and mature Nanog transcripts were developed using the designer tool from Stellaris (LGC Biosearch Technologies) (Table S1). 30 probes (TAMRA conjugated) for mature Nanog mRNA were targeted towards the 3′ GFP segment of transcripts. 48 probes (Quasar 670 conjugated) for nascent Nanog mRNA were targeted towards the first intronic sequence as taken from the mm10 genome reference. Probes were designed using a masking level of 5, and at least 2 base pair spacing between single probes.

1×10^5^ Nanog-GFP mESCs were seeded into each well of a gelatin-coated, 35mm Ibidi dish (quad-chambered, cat:80416) in 2i/LIF media. 24 hours following seeding, media was replaced with 2i/LIF containing 10µM IdU or equivalent volume DMSO. After 24 hours of treatment, cells were then fixed with DPBS in 4% paraformaldehyde for 10 minutes. Fixed cells were washed with DPBS and stored in 70% EtOH at 4°C for one hour to permeabilize the cell membranes. Probes were diluted 200-fold and allowed to hybridize at 37°C overnight. Wash steps and DAPI (Thermo) staining were performed as described (https://www.biosearchtech.com/support/resources/stellaris-protocols).

To minimize photo-bleaching, cells were imaged in a buffer containing 50% glycerol (Thermo), 75 µg/mL glucose oxidase (Sigma Aldrich), 520 µg/mL catalase (Sigma Aldrich), and 0.5 mg/mL Trolox (Sigma Aldrich). Images were taken on a Zeiss Axio Observer Z1 microscope equipped with a Yokogawa CSU-X1 spinning disk unit and 100x/1.4 oil objective. Approximately 20 xy locations were randomly selected for each condition. For each xy location, Nyquist sampling was performed by taking 30, 0.4µM steps along the z-plane.

Image analysis and spot counting was performed using FISH-quant *(88)*. Cells were manually segmented and analysis was conducted on cells of a similar size to minimize extrinsic noise. Transcriptional centers (TCs) were identified by signal overlap in exon, intron and DAPI channels. The amount of nascent mRNA at TCs was quantified through a weighted superposition of point spread functions.

### Rate calculations for random-telegraph model

From smRNA-FISH data for Nanog, the kinetic parameters of the random-telegraph model were inferred using the empirically derived values of mRNA mean (*µ*), mRNA Fano factor (*Fano*), transcriptional center frequency (*f*_*ON*_) and transcriptional center size (*TC*_*mRNA*_) *(36)*. The transcription rate (*k*_*tx*_) is calculated as:

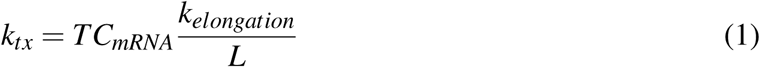

where *k*_*elongation*_ is the elongation rate of RNAPII (1.9 kb/min) *(96)* and *L* is the length of the transcribed region of Nanog. The degradation rate (*k*_*decay*_) is calculated as:

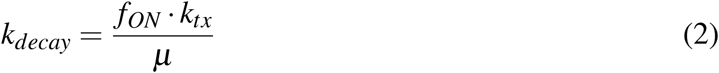

The rate of promoter activation (*k*_*ON*_) is given by:

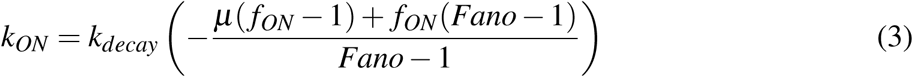

The rate of promoter inactivation (*k*_*OFF*_) is given by:

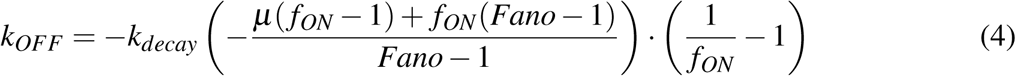

### Extrinsic Noise Filtering on Flow Cytometry Data

All flow cytometry data were collected on BD FACSCalibur, LSRII or LSRFortessa X-20 with 488-nm laser used to detect GFP. For all measurements of Nanog-GFP mean and variability, *>*50k cells are collected per sample. Gating of cytometry data was performed with FlowJo. Prior to quantification of Nanog-GFP mean and variability, the smallest possible forward- and side-scatter region containing at least 3k cells was used to isolate cells of similar size and shape. This filters out gene expression variability arising from cell-size heterogeneity as previously established *(25,27,36)*.

### Cell-cycle Analysis by Propidium-Iodide Staining

2×10^5^ Nanog-GFP mESCs were seeded in each well of a gelatin-coated, 6-well plate in 2i/LIF media. 24 hours following seeding, media was replaced with 2i/LIF media containing 10µM IdU or an equivalent volume of DMSO in triplicate for 24 hours. After treatment, cells were washed with DPBS, dissociated with TrypLE, pelleted, washed with DPBS, and resuspended in ice-cold 70% ethanol. Samples were stored overnight at -20°C and pelleted the following day at 200g for 5 minutes at 4°C. Cells were washed twice with DPBS supplemented with 0.5% BSA to prevent cell loss. Pellets were resuspended in 150µL of DPBS supplemented with 0.1mg/ml RNAse A (Thermo) and 30 µg/ml Propidium Iodide (Thermo). After overnight incubation at 4°C, cells were directly analyzed on a BD LSRII cytometer.

### Noise Enhancer Testing in Serum/LIF culture

Serum/LIF media was prepared with 85% DMEM (supplemented with 2mM of L-glutamine), 15% FBS, 0.1mM 2-mercaptoethanol, and 1000U/ml of LIF (Sigma Aldrich). Nanog-GFP mESCs grown feeder-free in 2i/LIF were passaged and seeded onto gelatin-coated 10cm dishes in serum/LIF media. Cells were passaged twice in serum/LIF media prior to noise enhancer testing. 4×10^5^ Nanog-GFP mESCs were seeded into each well of a gelatin-coated 6-well plate in serum/LIF media. 24 hours following seeding, media was replaced with serum/LIF supplemented with either 10µM IdU or equivalent volume DMSO in triplicate. After 24 hours of treatment, cells were run unfixed and live on BD LSRII flow cytometer.

### Sox2 two-color reporter assay

The endogenous alleles of Sox2 are tagged with P2A-mClover and P2A-tdTomato. Both fluorophores have a PEST tag, thus shortening their half-lives to approximately 2.5 hours. 2×10^5^ Sox2-dual-tag mESCs were seeded in each well of a gelatin-coated, 6-well plate in 2i/LIF media. 24 hours following seeding, cultures were replenished with 2i/LIF media containing 10µM IdU or an equivalent volume of DMSO in triplicate for 24 hours. Cells were run unfixed and live on BD LSRII flow cytometer. Intrinsic noise was calculated as in Elowitz et. al *(42)*. Data from all three replicates were pooled together. No cell-size gating was performed as assay allows for separation of extrinsic noise. To align fluorescence values of mClover and tdTomato on the same scale, each cell’s fluorescence intensity was normalized to the mean expression level of that fluorophore for the population. Since Sox2 expression spans several orders of magnitude, cells were binned according to their total Sox2 expression (mClover + tdTomato). Bins with fewer than 100 cells were discarded. Intrinsic noise (CV^2^) of Sox2 expression for each bin was calculated using the following formula:

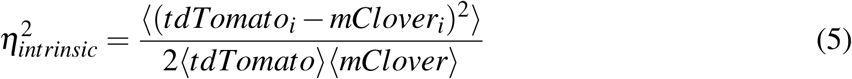

This value was then multiplied by the mean Sox2 expression for each bin to obtain the Fano factor. Given that the number of cells in each bin differs and variance estimates are affected by sample size, we calculated 95% confidence intervals around the Fano factor for each bin through bootstrapping. Bin populations were resampled 10,000 times with replacement.

### UV stress assay

1×10^5^ Nanog-GFP mESCs were seeded in each well of gelatin-coated, 12-well plates in 2i/LIF media. 24 hours following seeding, cultures were exposed to 3kJ of 365nM light (Fotodyne UV Transilluminator 3-3000 with 15W bulbs) for 15, 30 or 60 minutes at room temperature in the dark. Control plates were left at room temperature in the dark for equivalent periods of time. Cells from UV-exposed and control plates were run unfixed and live on BD FACS Calibur cytometer 1,2,4,8, and 12 hours post-exposure in replicate. Extrinsic noise filtering via cell-size gating was performed prior to calculation of Nanog Fano factor.

### Live-cell time-lapse microscopy

1×10^5^ Nanog-GFP mESCs were seeded into each well of a gelatin-coated, 35mm Ibidi dish (quad-chambered, cat:80416) in 2i/LIF media. 24 hours following seeding, media was replenished with 2i/LIF containing 10µM IdU or equivalent volume DMSO in replicate. Time-lapse imaging commenced immediately after addition of compounds with IdU- and DMSO-treated cells imaged in the same experiment (neighboring wells). Imaging was performed on a Zeiss Axio Observer Z1 microscope equipped with Yokogawa CSU-X1 spinning disk unit and a Cool-SNAP HQ2 14-bit camera (PhotoMetrics). 488nM laser line (50% laser power, 500-ms excitation) was used for GFP imaging. Samples were kept in an enclosed stage that maintained humidified conditions at 37°C and 5% CO_2_. Images were captured every 20 minutes for 24 hours. For each xy location, three z-planes were sampled at 4-µm intervals. The objective used was 40x oil, 1.3 N.A.

Cell segmentation, tracking and GFP quantification were carried out using CellProfiler *(95)*. Tracking of cells was manually verified. Segmented cells tracked for less than 4 hours were discarded. Cell division triggered the start of 2 new trajectories. After illumination correction and background subtraction, the mean GFP fluorescence intensity of a segmented cell was taken from each z-plane and averaged over the entire z-stack. For each trajectory, noise autocorrelation (*τ*_1*/*2_) and noise magnitude (intrinsic-CV^2^) were calculated as previously described *(52)*. Fluorescence trajectories were first detrended (normalized) by subtracting the population time-dependent average fluorescence to isolate intrinsic noise. Distributions of noise frequency ranges (*F*_*N*_) were extracted from normalized autocorrelation functions (ACFs) of individual trajectories, where 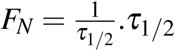 is the value of *τ* (lag time) where the normalized ACF reaches a value of 0.5.

### Nucleoside analog screening

14 nucleoside analogs (compound names and sources listed in Table S3) were resuspended in DMSO. 1×10^5^ Nanog-GFP mESCs were seeded in gelatin-coated 12-well plates in 2i/LIF media. 24 hours after seeding, media was swapped with 2i/LIF containing 10µM of nucleoside analog or equivalent volume DMSO in replicate. After 24 hours of treatment, cells were run unfixed and live on BD LSRII cytometer. Extrinsic noise filtering via cell-size gating was performed prior to calculation of Nanog Fano factor. Fano factor for Nanog-GFP expression for each treatment was normalized to DMSO control.

### Generation of stable CRISPRi Nanog-GFP mESC line

To stably integrate the CRISPRi machinery into the ROSA26 locus of Nanog-GFP mESCs, AAVS1 homology arms of the CRISPRi knockin construct (krab-dCas9-p2a-mCherry, Addgene:73497) were swapped with ROSA26 homology arms. The dox-inducible promoter of this construct was replaced with a constitutive CAGGS promoter and the kanamycin resistance cassette was replaced with puromycin resistance. Two million Nanog-GFP mESCs were nucleofected with the CRISPRi knockin construct and left to recover for 48 hours. Puromycin (1µg/ml) selection was run until single colonies could be picked. Clonal CRISPRi Nanog-GFP mESC lines were assessed for mCherry expression and ability to knockdown Nanog. We selected the clone with the highest percentage of mCherry-positive cells.

### CRISPRi gRNA design and cloning

gRNA sequences were were taken from the mCRISPRi-v2 library *(91)*. gRNA oligos were annealed and cloned into the pU6-sgRNA EF1Alpha-puro-T2A-BFP lentiviral vector (Addgene:60955) using the BstXI/BlpI ligation strategy *(91)*.

### CRISPRi screening for genetic dependencies of noise enhancer

25 genes involved in nucleotide metabolism, DNA repair, and chromatin remodeling were screened for their potential role in noise enhancement from IdU. Three gRNAs were designed per gene (gene names and gRNA sequences listed in Table S4). Three non-targeting controls (scrambled gRNAs) were taken from the mCRISPRi-v2 library *(91)*. Each gRNA expression plasmid was separately packaged into lentivirus in HEK293T cells as previously described *(91)*. For each gRNA lentivirus, 1.5×10^5^ CRISPRi Nanog-GFP mESCs were spinoculated with filtered viral supernatant for 90 minutes at 200 x g in replicate. Following spinoculation, infected cells were seeded into gelatin-coated, 6-well plates in 2i/LIF media. 48 hours following seeding, media was swapped with 2i/LIF supplemented with either 10µM IdU or equivalent volume DMSO. Consequently, for every knockdown there is a DMSO and IdU treatment group. After 24 hours of treatment, cells were run unfixed and live on a BD LSRII flow cytometer. To minimize technical variability, analysis was restricted to cells with homogeneous levels of dCas9-KRAB and gRNA expression through stringent gating on mCherry/BFP double-positive cells. Extrinsic noise filtering through cell-size gating was then applied. For each gRNA, Nanog Fano factor for the DMSO and IdU treatments were normalized to the Nanog Fano factor of the non-targeting controls treated with DMSO.

### qPCR verification of CRISPRi knockdown

To verify CRISPRi knockdown of Apex1 and Tk1, each of the six gRNA-expression plasmids targeting these two genes along with a non-targeting control and empty vector were packaged into lentivirus. 1.5×10^5^ CRISPRi Nanog-GFP mESCs were spinoculated with filtered viral supernatant for 90 minutes at 200 x g in replicate. Following spinoculation, infected cells were seeded into gelatin-coated, 6-well plates in 2i/LIF media. 72 hours following seeding, 1×10^6^ mCherry/BFP double-positive cells from each infected cell population were sorted on a FACSAria II. Total RNA was extracted using an RNeasy Mini Kit (QIAGEN cat:74104) and reverse-transcribed using a QuantiTect Reverse Transcription Kit (QIAGEN cat:205311). cDNA from each independent biological replicate was plated in triplicate and run on a 7900HT Fast Real-Time PCR System (Thermo) using designed primers (Table S5) and Fast SYBR Green Master Mix (Applied Biosystems, cat:4385612). Expression of GAPDH was used for normalization. Relative mRNA levels of Apex1 and Tk1 were calculated by the ΔΔ*C*_*t*_ method using the empty-vector populations as the control. All reported levels of repression are relative to the non-targeting control.

### Tk1 competition assay

1×10^5^ Nanog-GFP mESCs were seeded in each well of gelatin-coated, 12-well plates in 2i/LIF media. 24 hours following seeding, media was replaced with 2i/LIF supplemented with 10µM IdU in combination with thymidine (Sigma cat:T1895) or uridine (Sigma cat:U3003) at concentrations ranging from 0 to 100µM. Concentration combinations were done in triplicate. After 24 hours of treatment, cells were run unfixed and live on BD FACS Calibur cytometer. Extrinsic noise filtering via cell-size gating was performed prior to calculation of Nanog Fano factor.

### Biotinylated-trimethylpsoralen (bTMP) supercoiling assay

1×10^5^ Nanog-GFP mESCs were seeded into each well of a gelatin-coated, 35mm Ibidi dish (quad-chambered, cat:80416) in 2i/LIF media. 24 hours following seeding, media was replaced with 2i/LIF supplemented with 10µM IdU, 10µM IdU & 100µM CRT0044876, or equivalent volume DMSO in replicate. After 24 hours of treatment, media was replaced with 2i/LIF supplemented with 1µM aphidicolin for two hours. For control experiments, Nanog-GFP mESCs were cultured with or without 10µM IdU for 24 hours followed by treatment with 100µM bleomycin for one hour. Cells were then washed 1xDPBS and then permeabilized with 0.1% Tween-20 in DPBS for 15 minutes. Cells were then incubated with 0.3mg/ml EZ-Link Psoarlen-PEG3-Biotin (Thermo cat:29986) for 15 minutes. Cultures were then exposed to 365nM light (AlphaImager HP with 15W bulbs, ProteinSimple) for 15 minutes at room temperature. Cells were then washed 2xDPBS, fixed with cold 70% ethanol for 30 minutes at 4°C, and then washed 2xDPBS. Cells were then incubated with Alexa Fluor 594 Streptavidin (Thermo cat:S32356) for one hour at room temperature in the dark, washed 2xDPBS, and stained with DAPI for 10 minutes at room temperature in the dark. Cells were imaged in a buffer containing 50% glycerol (Thermo), 75 µg/mL glucose oxidase (Sigma Aldrich), 520 µg/mL catalase (Sigma Aldrich), and 0.5 mg/mL Trolox (Sigma Aldrich). Images were taken on a Zeiss Axio Observer Z1 microscope equipped with a Yokogawa CSU-X1 spinning disk unit and 63x/1.4 oil objective. Approximately 20 xy locations were randomly selected for each condition. For each xy location, three z-planes were sampled at 4-µm intervals. Nuclear segmentation using DAPI signal and quantification of psoralen staining intensity were carried out using CellProfiler. After illumination correction and background subtraction, the mean psoralen fluorescence intensity of a segmented nucleus was taken from each z-plane and averaged over the entire z-stack.

### 96 dose combination for testing of noise phase space

Compound plates containing 96 concentration combinations of IdU, BrdU, or HmU with CRT0044876 were prepared by the Gladstone Assay Development and Drug Discovery Core using an Agilent Bravo liquid handling system. All wells contained equivalent volumes of DMSO. Compound mixtures were suspended in 200µL of 2i/LIF media. 1×10^4^ Nanog-GFP mESCs were seeded into each well of a gelatin-coated, 96-well dish in 200µL of 2i/LIF media. 24 hours after seeding, 100µL of media was removed from each well and 100µL of compound-containing 2i/LIF was added in replicate. Layout and final concentrations of treatments are listed in Table S6. IdU and BrdU concentrations ranged from 0 to 50µM while HmU concentrations ranged from 0 to 10µM. CRT0044876 ranged from 0 to 150µM. After 24 hours of treatment, cells were detached using TrypLE and plates were run on BD LSRFortessa high-throughput system. After extrinsic noise filtering via cell-size gating, Nanog mean and Fano factor for each treatment were normalized to DMSO control well. Reported fold changes in mean and Fano factor are the average of two replicates.

### Estimation of promoter toggling kinetics from scRNA-seq data

Gene expression data from the scRNA-seq dataset were fit to the 2-state model using the D3E algorithm, allowing for estimation of *k*_*ON*_, *k*_*OFF*_, *andk*_*tx*_ in proportion to the rate of mRNA degradation which is the lone parameter that is not estimable from this dataset alone *(83)*. Parameter estimation was conducted using the methods of moments approach with the normalise and removeZeros options. Analysis was run for the 945 genes classified as highly variable according to the BASiCS algorithm. The Cramer-von Mises test was used for goodness-of-fit testing. Values of *k*_*decay*_ were then retrieved from an existing dataset of mRNA degradation rates in mESCs *(92)*, with the assumption that degradation rates are unchanged between DMSO and IdU conditions. Parameter estimates were then verified against experimental values of mean mRNA counts using the following relationship: 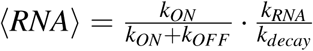. Genes whose predicted mean was within 10% of experimental value were used for downstream analysis. 314 genes passed this filtering process based on availability of mRNA degradation rates and alignment of parameter estimates with expected mean mRNA counts.

### Mathematical modeling and simulations

Detailed descriptions of model development, parameterization, evaluation, and sensitivity analysis can be found in the supplementary text along with details of Gillespie simulations.

### Estimation of cellular developmental potential and Waddington Landscape reconstruction

The HopLand algorithm (continuous Hopfield network) was used to create a predictive model of gene-gene interactions from scRNA-seq data *(86)*. As input, raw count data for a total of 800 randomly chosen cells (400 from DMSO and 400 from IdU treatment groups) were used. Each neuron of the network corresponds to a gene. Genes whose variance fell within in the top 10% were used for construction of a neural network resulting in 512 nodes. The weight matrix describing pair-wise interactions between nodes was initialized using the gene-gene Pearson correlation matrix. A Gaussian process latent variable model (GP-LVM) was used for dimensionality reduction to create a 2-D map of cell clustering (x and y coordinates on Waddington Landscape). Energy values (z coordinate on landscape) were calculated using the Lyapunov function which is a measure of stability. Lower energy values indicate greater proximity to an equilibrium point (attractor state) and thus less developmental potential.

### Cellular reprogramming assays

Two cellular reprogramming systems were tested in this study: (1) Nanog-GFP secondary mouse embryonic fibroblasts (MEFs) harboring stably integrated, doxycycline-inducible cassettes for Oct4, Sox2, and Klf4. GFP is expressed from the endogenous Nanog locus. (2) Oct4-GFP primary MEFs that express GFP from the endogenous Oct4 locus.

Secondary MEFs were seeded onto gelatin-coated, 12-well plates at a density of 10,000 cells/cm^2^ in MEF medium (DMEM supplemented with 10% FBS and 0.1mM non-essential amino acid, and 2mM Glutamax). 24 hours after seeding, wells were washed with DPBS and media was switched to ESC media (knockout DMEM, 10% FBS, 10% KSR, 2mM Glutamax, 0.1mM non-essential amino acid, 0.1mM 2-mercaptoethanol, 10^3^ units/ml leukemia inhibitory factor) supplemented with 1µg/ml doxycycline. Additionally, IdU (1uM or 4uM) or equivalent volume DMSO (Day 0) were added to media. 48 hours after the start of IdU treatment, wells were washed with DPBS and media was replaced with ESC media supplemented with 1µg/ml doxycycline alone. Media was refreshed every other day until day 10 of reprogramming. Alkaline phosphatase staining was performed according to manufacturer’s instructions using the Alkaline Phosphatase Detected Kit (Millipore). For flow cytometric analysis of Nanog-GFP expression, cells were dissociated with TrypLE and run unfixed on BD FACS Calibur cytometer.

Oct4-GFP primary MEFs were transduced with lentiviral vectors encoding Oct4, Sox2, Klf4, and c-Myc. Lentiviruses encoding these factors were individually packaged in PLAT-E cells (ATCC) using pMX-based vectors. 48 hours after transfection of lentiviral vectors, viral supernatant was collected and filtered. For infection, Oct4-GFP primary MEFs were seeded on gelatin-coated, 6-well plates at a density of 10,000 cells/cm^2^ in MEF medium 24 hours prior to transduction (Day -2). Oct4, Sox2, Klf4, and c-Myc viruses were mixed in equal volume along with 5µg/ml polybrene and incubated with primary MEFs for 24 hours in MEF medium (Day -1). Following infection, wells were washed with ESC media and cells were incubated with ESC media supplemented with 10µM Forskolin, 1mM Valproic Acid, 4µM IdU or equivalent volume DMSO (Day 0). ESC media was refreshed every other day. IdU supplementation was discontinued after 48 hours while Forskolin and Valproic Acid were kept in media continuously. Oct4-GFP(+) colonies were counted on days 8, 10 and 12.

### Bulk RNA-seq of secondary MEFs undergoing reprogramming

Secondary MEFs were seeded onto gelatin-coated, 6-well plates at a density of 10,000 cells/cm^2^ in MEF medium. For each timepoint (2- and 5-day), 4 wells were seeded (2 replicates for standard reprogramming and 2 replicates for IdU-assisted reprogramming). 24 hours after seeding, wells were washed with DPBS and media was switched to ESC media supplemented with 1µg/ml doxycycline. Additionally, 4µM IdU or equivalent volume DMSO (Day 0) were added to media. 48 hours after the start of reprogramming, cells for the 2-day timepoint in DMSO and IdU conditions were dissociated with TrypLE, pelleted, and snap frozen with liquid nitrogen. Media in the wells for the 5-day timepoint was refreshed with ESC media supplemented with 1µg/ml doxycycline alone. This was repeated on day 4. On day 5, remaining cells were dissociated and frozen identically to that of the 2-day timepoint.

RNA was extracted from each cell pellet using a RNeasy minikit (Qiagen) according to manufacturer’s instructions. A total of 8 cDNA libraries were prepared with an NEBNext Ultra II RNA Library Prep kit (NEB, cat:E7770S) and sequenced with an Illumina HiSeq4000. Sequencing yielded a median of ≈ 50 million single-end reads per library. Read quality was checked via FASTQC. Reads were aligned to the mm10 reference genome using TopHat with default parameters. Transcript level quantification was performed using Cufflinks with default parameters.

## Supplementary Text

### 1 Overview of tested models

To delineate how Apex1 alters transcriptional dynamics of Nanog gene expression in a way that increases noise without changing mean, we developed a series of models that allow for Apex1 interaction at different stages of the transcription process. Through stochastic simulation of these models and comparison to experimental data, the aim is to develop greater mechanistic insight into what stages of the transcription process Apex1 affects. The tested models listed in supplementary figure 20 are adapted from the two-state random telegraph model. We assume in each of the models that IdU incorporation into the genome leads to recruitment of Apex1 (*k*_*incorpo*_). The resulting interaction results in a transcriptionally non-productive state. Unbinding of Apex1 is triggered by completion of repair (*k*_*repair*_).

### 2 Detailed mathematics and derivation of parameter constraints

We derive here some relations, at equilibrium, between the kinetic rates of the diverse models. These relationships are then used to constrain the parameter phase space for a given set of data.

#### 2.1 Model 0

This is the *null* model and consists of only the canonical two-state random-telegraph. This model is used as a *null* hypothesis, in particular for *log-likelihood* and *AIC*-based model selection.

#### 2.2 Model 1

We assume that IdU incorporation and subsequent interaction of Apex1 with the chromatin, occurs only in the OFF state of the promoter. Biologically, this may occur if control mechanisms inhibit DNA repair during active transcription.

The differential equations governing the gene fractions in the different states and the mRNA counts are as follows:

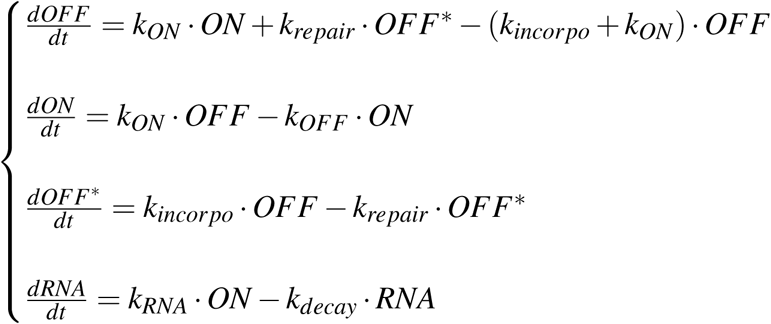

Using the fact that:

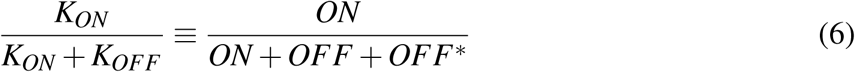

we derive the following constraint between *k*_*repair*_ and *k*_*incorpo*_, where *K*_*OFF*_ represents the transition rate to the macroscopic OFF state consisting of the OFF (*k*_*OFF*_) and OFF*(*k*_*incorpo*_) states in Model 1:

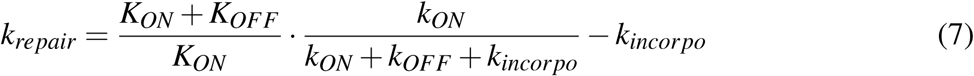

#### 2.3 Model 2

In this model, Apex1 can interact with chromatin in both the ON and OFF states of the promoter. If Apex1 interacts with the chromatin in the ON state, this leads to a turning off of the system. Molecularly this may be seen as a strong inhibitory effect mediated by Apex1: the interaction may recruit chromatin modifiers (e.g. histone deacetylases, histone methyltransferases) that silence gene expression. In the same way, stalled polymerases, at both the promoter proximal region and further in the gene, may unbind DNA.

This model can be described by the following set of ODEs:

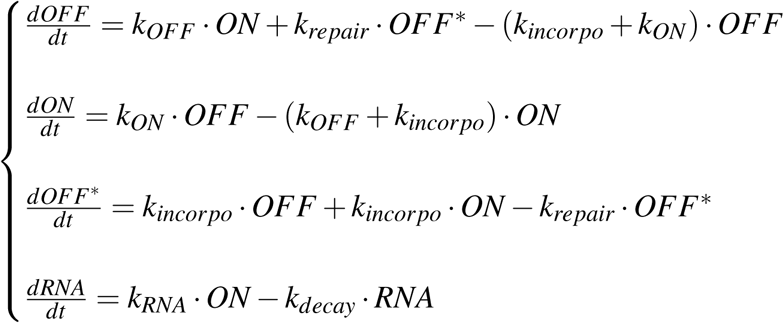

Using equation (6) we derive the following constraint between *k*_*repair*_ and *k*_*incorpo*_:

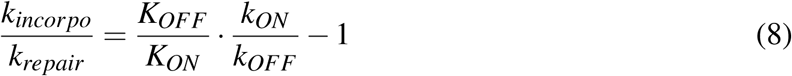

#### 2.4 Model 3

Here we assume that Apex1 can still interact in the ON state but that does not alter the “primed” characteristic of the gene expression system. The system is thus in a *transcriptionally non-productive* ON* state. In other words transcription can not be achieved when Apex1 interacts with the chromatin but the transcriptionally permissive chromatin and molecular context is not altered: primed polymerases remain, transcription enhancing epigenetic marks are not erased, etc.

This model can be described by the following set of ODEs:

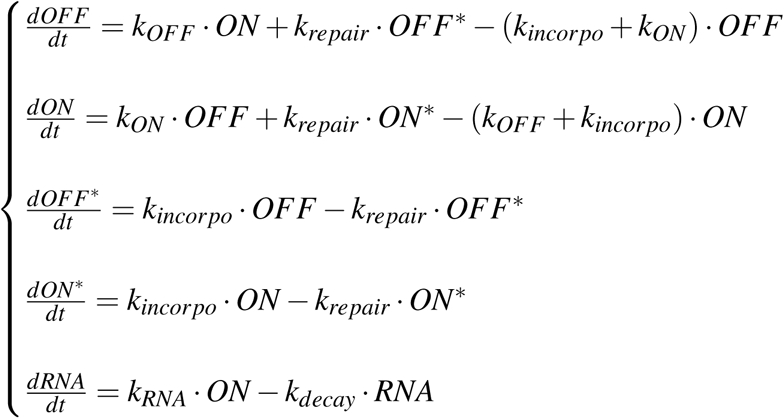

We can thus derive the following constraint:

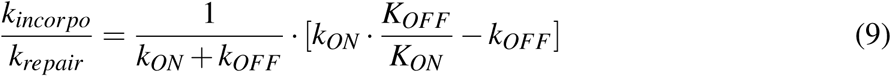

The details of this derivation are described below for Model 4.

#### 2.5 Model 4

Model 4 is based on Model 3, but an amplification step was added. When the system transitions from ON* to ON, the basal transcription rate 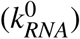 increased by a multiplicative factor: 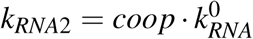. Thus we assume that there is molecular memory of the repair event. This may be rooted in a modification of supercoiling, polymerase accumulation, or chromatin remodeling.

This model can be described by the following set of ODEs :

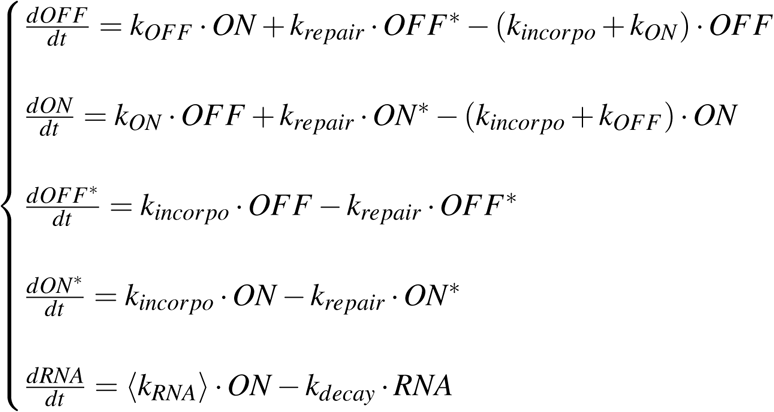

Where 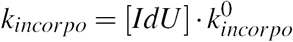. We assume that IdU is in excess and thus [IdU] remains constant. At equilibrium :

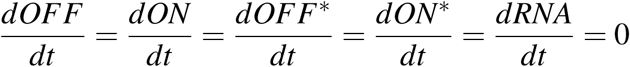

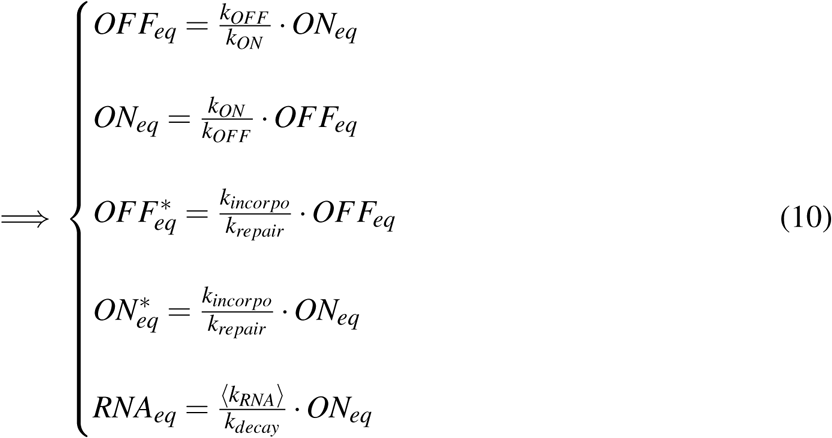

By definition:

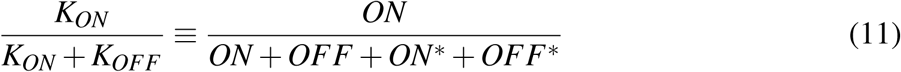

where *K*_*OFF*_ represents the transition rate to the macroscopic OFF state (defined by no transcription) consisting of the OFF, OFF*, and ON* states. Using the set of equations in (10) and (11) we obtain:

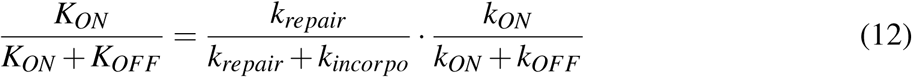

Equation (12) can be seen as :

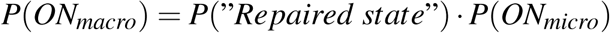

Because *P*(*ON*_*macro*_) ≡ *P*(“*Repaired state*” ∩ *ON*_*micro*_) we can deduce that “*Repaired state*” and *ON*_*micro*_ are independent probabilistic events.

Equation (12) can be rewritten as:

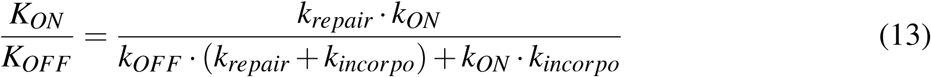

Which gives us:

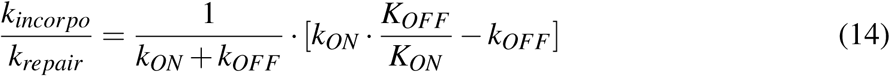

Let us define ⟨*k*_*RNA*_⟩ and 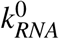 as the mean transcription rate in the presence of IdU and the transcription rate in the control condition (DMSO), respectively. By construction of the model:

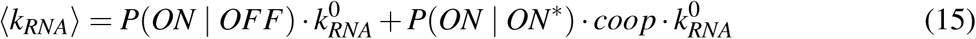

The *coop* term represents the amplification of the transcription rate following completion of repair. *P*(*ON* | *OFF*) and *P*(*ON* | *ON*^∗^) represent the probability that the gene transitioned to the ON state from the OFF and ON* states respectively.

Or :

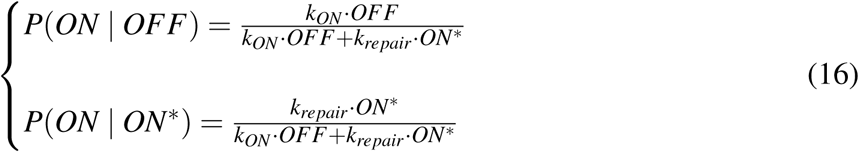

Combining equations (14) and (16) we get:

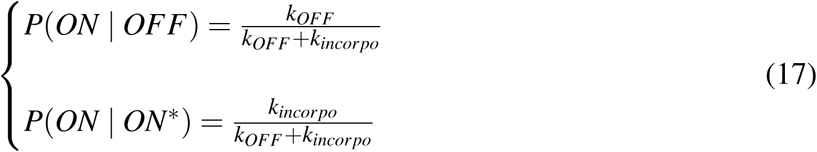

Rewriting (15) using (17) we obtain :

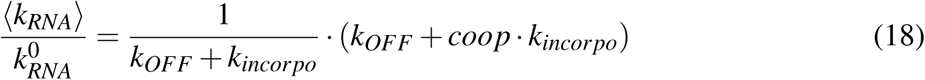

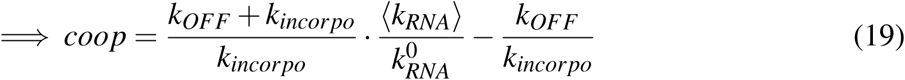

Equation (19) can be rewritten to explicitly take into account [IdU] :

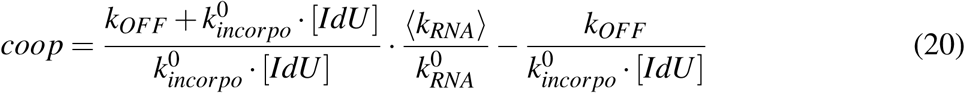

#### 2.6 Model 5

In this last model we make the same assumptions as in model 4 except that Apex1 interaction with chromatin occurs *only* in the ON state. This does not imply that IdU incorporation and subsequent Apex1 interaction only occurs in the ON state. Our assumption supposes that the interaction of Apex1 in the OFF state is negligible quantitatively speaking compared to that in the ON state.

The ODEs describing this model are as follows:

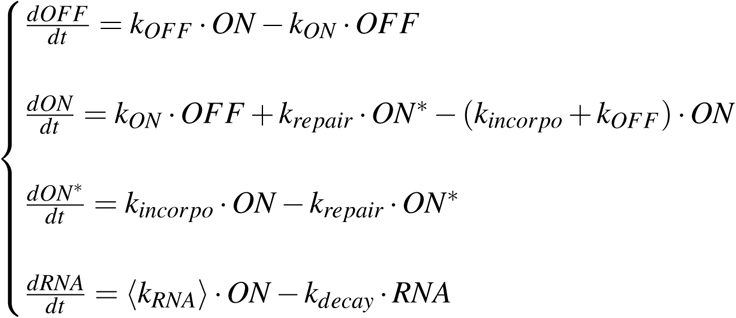

The *coop* expression remains unchanged from Model 4 but the ratio between *k*_*repair*_ and *k*_*incorpo*_ changes as follows:

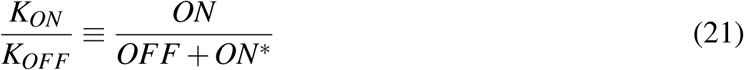

Thus we obtain:

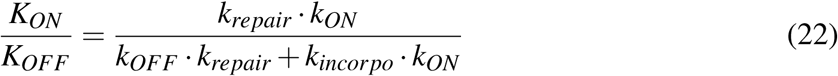

And :

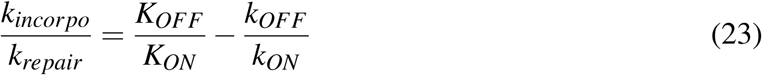

### 3 Chemical Master Equation

For all models we constructed an associated stochastic scheme. As an example, Model 5 can be rewritten using the following scheme:

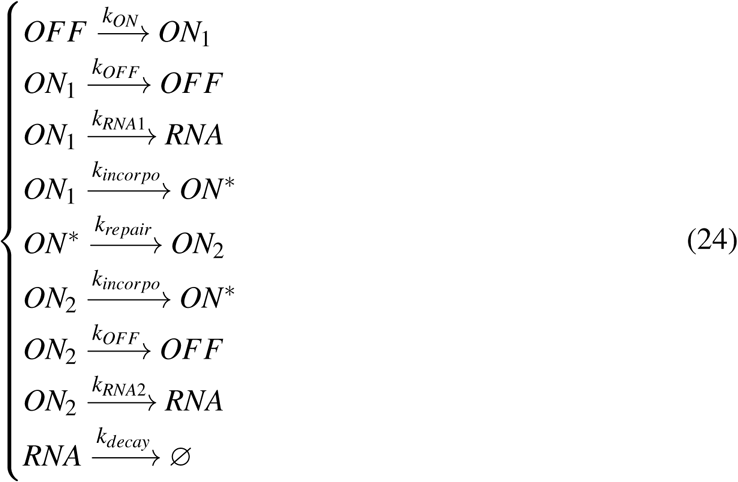

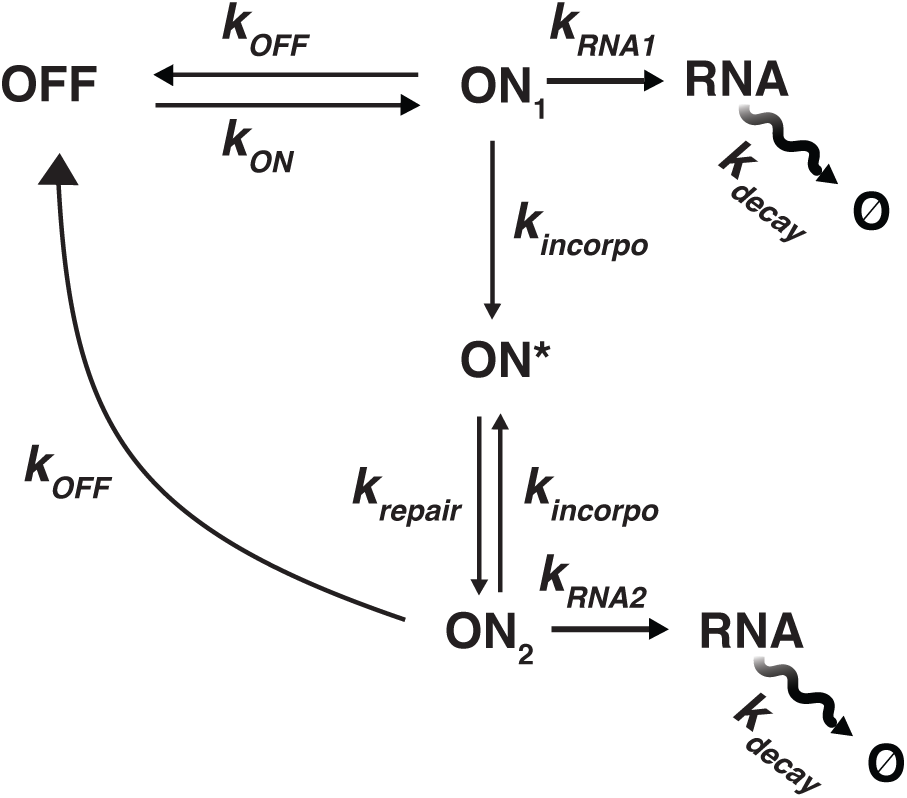

*ON*_2_ represents the repaired state of the gene, which results in a higher transcription rate (*k*_*RNA*2_). Using this scheme, we can construct the chemical master equation (CME) describing the time dependent distributions of mRNA copy number:

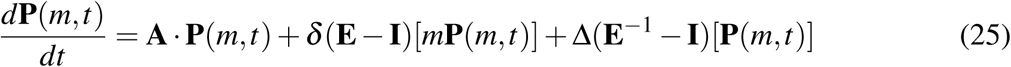

Where **A**, Δ, and *δ* are the transition, transcription, and degradation matrices respectively:

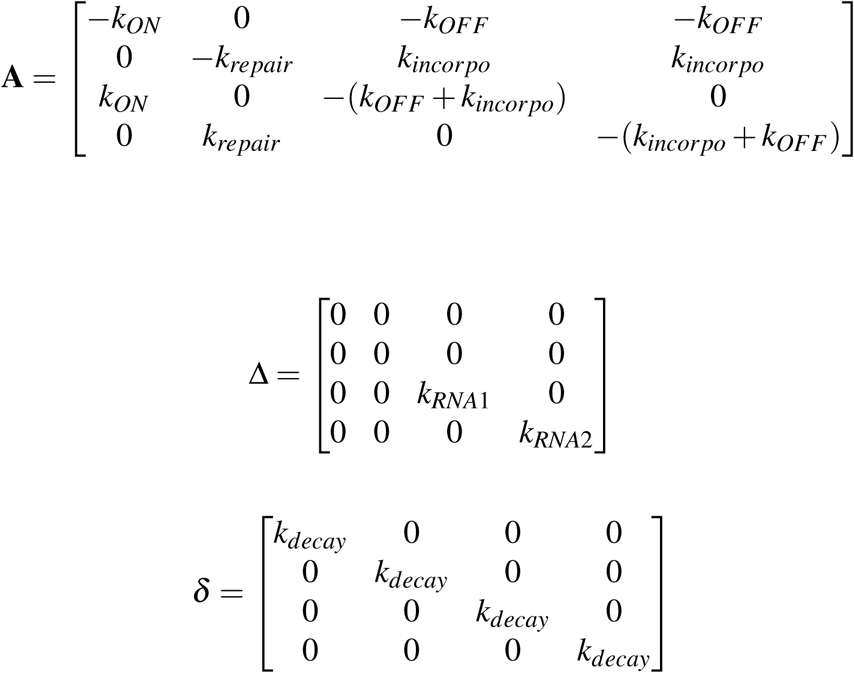

**P**(*m, t*) is a four-element column vector consisting of the time-dependent mRNA probability distributions while in the *OFF, ON*^∗^, *ON*_1_, *andON*_2_ states respectively. **E** and **E**^−1^ are the forward and backward shift operators while **I** is the identity matrix.

At steady-state, the mRNA probability distribution can be reconstructed as a sum of the binomial moments *(93)*

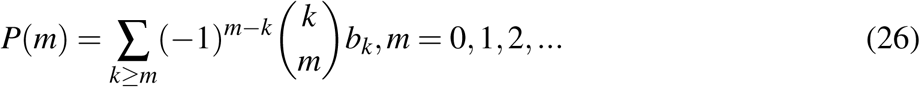

Where *b*_*k*_ is the *k*^*th*^ binomial moment of the distribution given by:

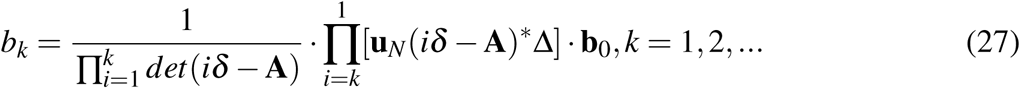

where **u**_*N*_ = [1,1,1,1]. (*iδ* − **A**)^∗^ and *det*(*iδ* − **A**) are the adjugate and the determinant of matrix (*iδ* − **A**) respectively. *b*_1_ is equivalent to the mean mRNA abundance of the system at equilibrium. **b**_0_ is the corresponding eigenvector for the zero eigenvalue of **A**. The 4 elements of **b**_0_ therefore represent the fraction of time spent in the *OFF, ON*^∗^, *ON*_1_, *and ON*_2_ states at equilibrium. The *n*^*th*^ component of **b**_0_ is given by:

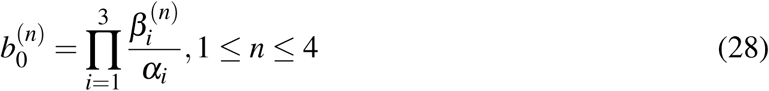

where *α*_1_, *α*_2_, *α*_3_ are the three non-zero eigenvalues of **A**. 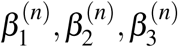 are the three eigen-values of the sub-matrix **M**_*n*_ which is constructed by removing the *n*^*th*^ row and *n*^*th*^ column of **A**.

The Fano factor for mRNA counts at equilibrium is then given by:

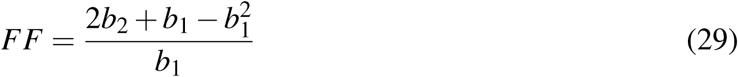

An exact simulation of such a stochastic process is given by the Gillespie algorithm. We implemented the algorithm using a homemade script in Julia 1.1.1. For each model, 1500 simulations were run for a virtual duration of 200h. Since the time is not discrete we used a “parsing” algorithm, based on recursive binary search, to align all the traces on a common time scale of 2000 intervals. For each interval of the discretized time we computed the average and variance of the number of mRNAs, taking into account all traces. The analytic relationships described above are used to verify the inferred kinetic rates from stochastic simulations.

### 4 Estimation of model parameters from experimental data

From the smRNA FISH data, we can infer the *macroscopic* kinetic rates (Table S2). The kinetic rates computed in the control condition (DMSO in Table S2) are at the basis of the simulations and have to be considered as constant for all the models and associated results.

Use of these macroscopic rates along with the relationships developed in section 2 results in one remaining degree of freedom for our models: *k*_*repair*_ (or *k*_*incorpo*_ for Model 1).

The confrontation between experimental data and the results of stochastic simulations for each of the models should allow us to define the consistency of our models and thus gain mechanistic insight into how Apex1 affects transcriptional dynamics.

### 5 Model selection: Comparison of simulation results to experimental data

#### 5.1 Information theory-based approach: MLE and Akaike’s criterion

##### 5.1.1 Workflow

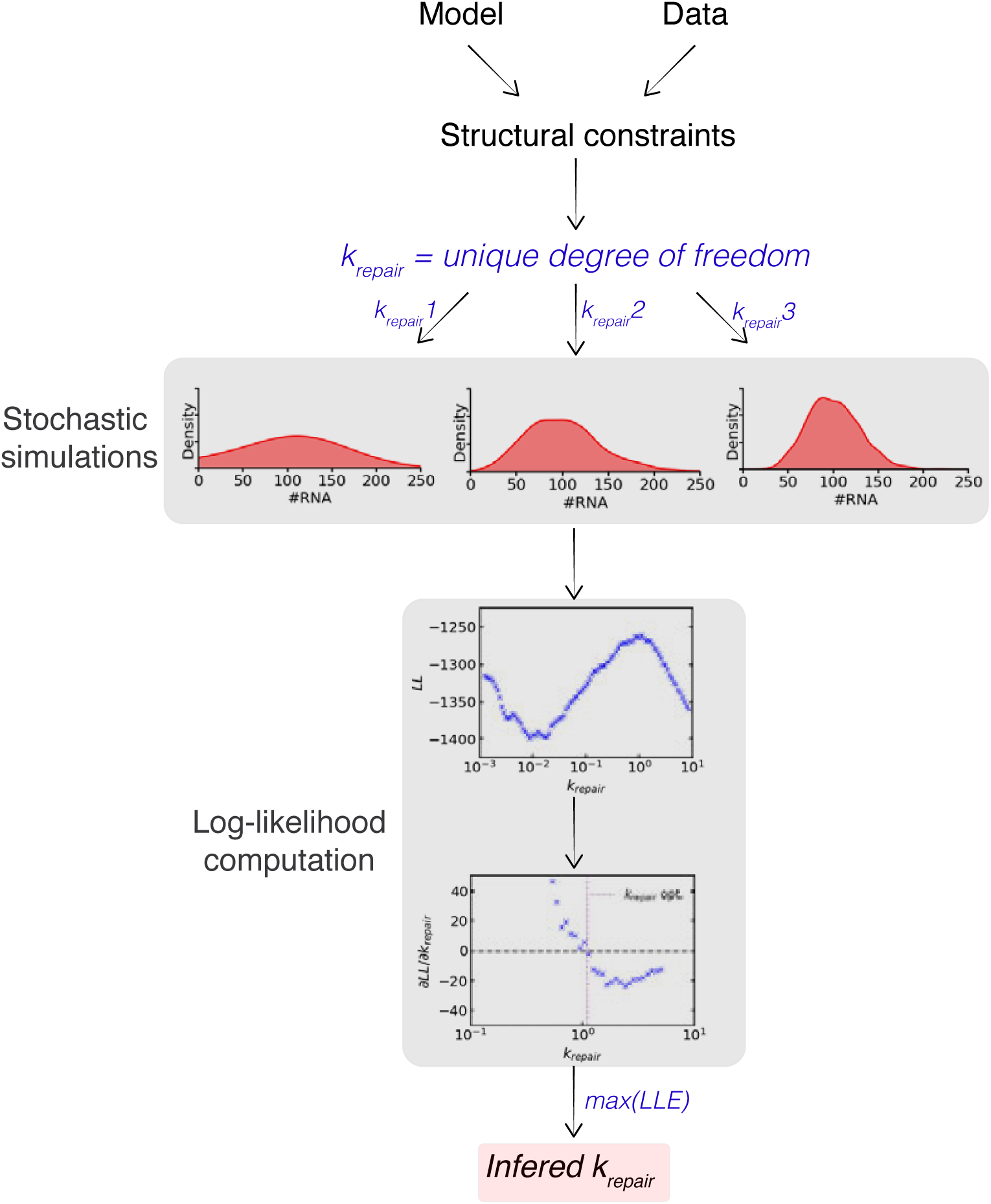

##### 5.1.2 Method of Log-likelihood and AIC computation

For each of the 500 chosen values (logarithmically spaced) of our unique degree of freedom (*k*_*repair*_), we ran 50,000 simulations for a total duration of 200h. Only the last point of each simulation was stored. Then the steady-state distribution of RNA was computed using the same binning as in the experimental data. The computation of the log-likelihood is as follows:

Let **X** be the vector of *n* empirical observations and let 𝒳_Δ_ : {*x*^Δ^, *x*^Δ^ ∈ [*x, x* + Δ]}, where Δ is the size of a bin and 𝒳_Δ_ is the set of bins. We define 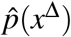 as the probability of observing an experimental value within a particular bin (*x*^Δ^). *P*(*x*^Δ^) is the probability of a given *x*^Δ^ ∈ [*x, x* + Δ] for a given model. The likelihood function ℒ is defined as :

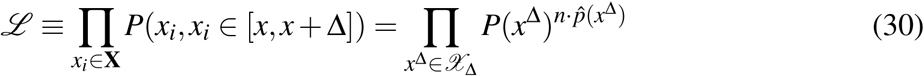

The log-likelihood is then given as:

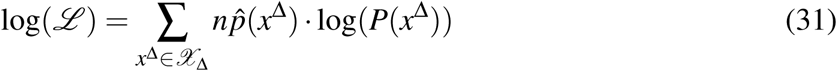

ℒ is a function of *k*_*repair*_. We then try to find the value of *k*_*repair*_ that maximizes log(ℒ):

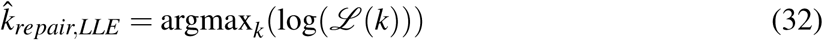

With *k*_*repair*_ ∈ [10^−4^, 10], by assumption. After computing log(ℒ ^(*k*)^) for each of the 500 values of *k*_*repair*_, we apply a smoothing (moving average) to the data, take the derivative, smooth the derivative, find the two points on each side of the abscissa, and then interpolate the point for which the derivative is equal to zero using a linear interpolation. The maximum log(ℒ) is computed after the first smoothing. Then, we compute the macroscopic behavior of the system using the inferred value of *k*_*repair*_.

The model selection is based on the *AIC* and the resulting measures Δ_*i*_*AIC* and *w*_*i*_ *(94)*. Because 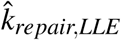 is dependent on the empirical distribution, we can assume that it is only an estimate of the *true* value, which we’ll call 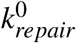. Thus we want to reduce as much as possible the distance between 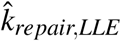 and 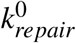. This optimization problem allows us to derive the so-called Akaike’s information criterion (AIC) as a measure to compare models. *AIC* is an estimate of the expected relative distance between the fitted model and the unknown true mechanism that actually generated the observed data:

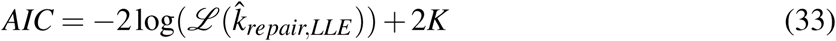

with *K* the number of degrees of freedom of the system.

##### 5.1.3 Results

As shown in Figures S20A-B, Model 5 is selected on the basis of AIC. Model 4 is second-best. Model 5 qualitatively and quantitatively matches experimental data with its inferred *k*_*repair*_.

##### 5.1.4 Parameter identifiability

We used bootstrapping to assess the quality of the inference of *k*_*repair*_ for Model 5 using MLE estimator. This method allows the computation of the confidence/credible interval (CI) for *k*_*repair*_, and, in the framework of Bayesianism, a posterior distribution *P*(*Data* | *k*_*repair*_) using a non-informative prior *(94)*. The distribution of the MLE is peaked around a particular value, suggesting parameter identifiability (Figure S20C).

#### 5.1 APE-based approach

##### 5.2.1 Workflow

The second approach for model selection that we employed involves comparison of the macroscopic behavior of simulation results to experimental data. After deriving constraints on the phase space spanned by *k*_*repair*_ we simulate each of the models over a range of *k*_*repair*_ values. From these stochastic simulations, we then extract the macroscopic behavior of the system for a given model. We computed the mean number of mRNA and the Fano factor at equilibrium, on the last 100 time points of the traces. *k*_*o f f*_ and *k*_*on*_ were computed using a non-linear curve fitting assuming exponentially distributed residence times (Poisson process). Both *ν*_*on*_ and the burst size were computed using their basic definitions. The density of mRNA population was computed using the last points of the simulations. A classic kernel density was used to represent the data.

For each model, we then infer the best value of *k*_*repair*_ based on the minimization of a loss function (absolute percentage error, a.k.a. APE). This quantitative approach is coupled with visual comparison of model behavior to experimental data. The model whose inferred *k*_*repair*_ value minimizes divergence between model and data behavior is thus chosen. A graphical representation of such a process is given below:

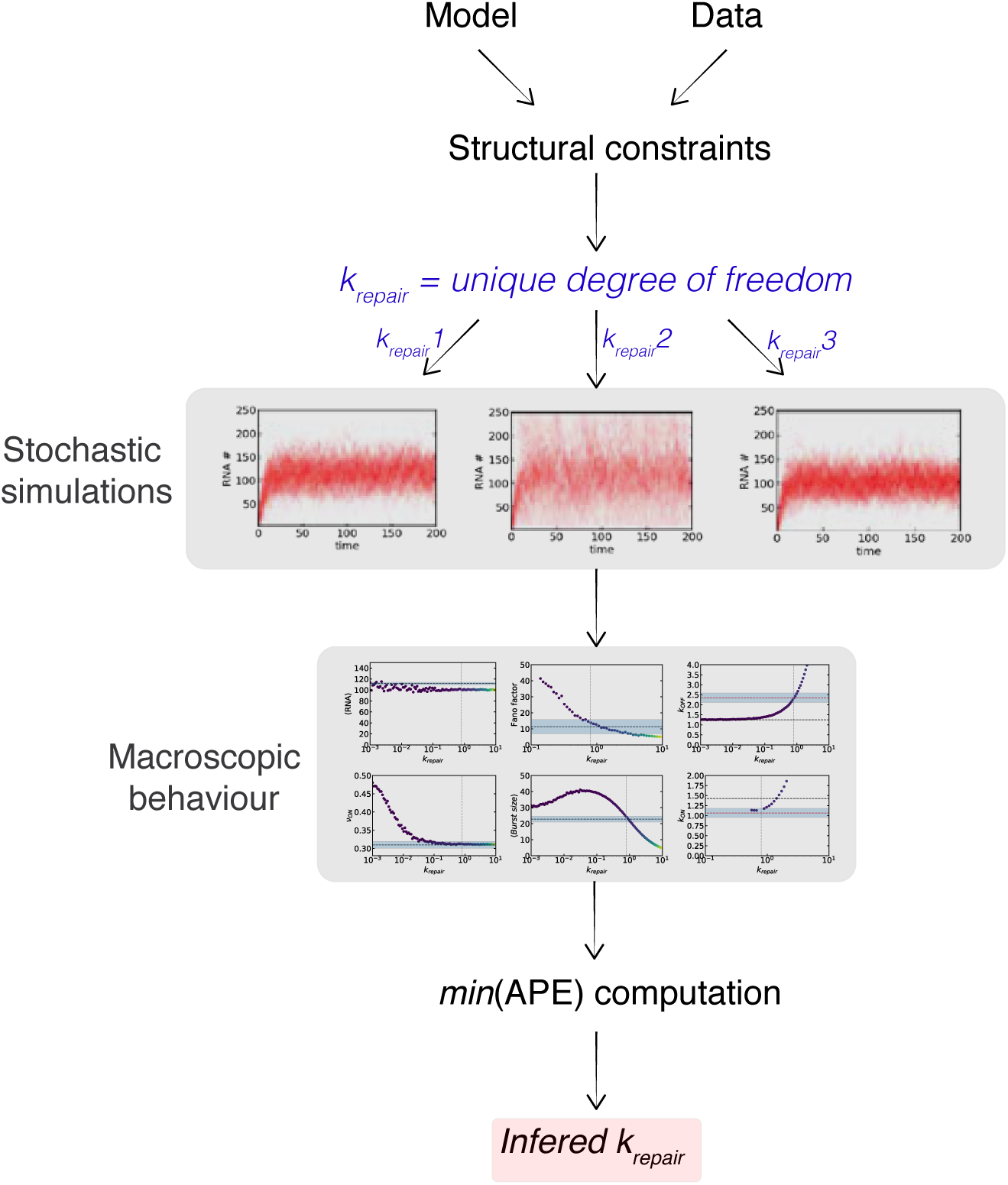

##### 5.2.2 Method of APE calculation

To discriminate between models based on their macroscopic behavior, we need a measure that quantifies the discrepancy between model-derived results and experimental data. Because the macroscopic observations (Fano factor, *k*_*ON*_, etc.) are of different orders of magnitude, we need a relative measure to avoid the largest parameters carrying the highest weight on the error. We chose to use the absolute percentage error (APE). The procedure is as follows:

Consider the vector **M** ≡ (*RNA, ν*_*ON*_, *BS, FF, k*_*ON*_, *k*_*OFF*_). Thus, **M**_*model*_ and **M**_*exp*_ contain all the macroscopic observations from modeling and experiment respectively. **M**_*model*_ is a function of *k*_*repair*_ or *k*_*incorpo*_ equivalently. 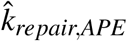,is the best inferred value of the degree of freedom for a particular model. It is given by:

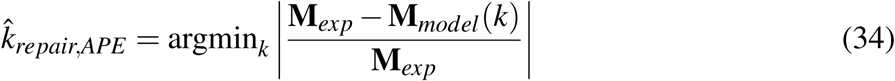

With *k* ∈ [10^−4^, 10], by assumption. This notation implies that we are minimizing the ℓ 1 norm (sum of the vector components). In an operative manner, we simulated each model for 250 logarithmic distributed values of *k*_*repair*_ using the previously described Gillespie algorithm.

To validate our approach, we devised an alternative loss function where the ℓ 1 norm is computed using a non-biased (i.e symmetric) measure of relative prediction accuracy: the absolute log accuracy (ALA). The procedure is as follows:

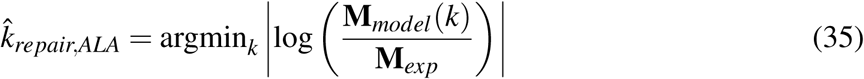

With *k* ∈ [10^−4^, 10], by assumption.

##### 5.2.3 Results

Using both the APE and ALA approaches, we obtained *exactly* the same results for parameter inference (except for model 1 for which 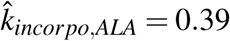) and model selection. According to the APE-based approach for parameter inference and model selection, model 5 best recapitulates the macroscopic behavior observed in the data (Figure S21).

#### 5.3 Selection of Model 5

Based on both the MLE- and APE-based approaches, model 5 best matches experimental FISH data and scRNA-seq data for Nanog. Model 5 is most similar to a transcription-coupled repair (TCR) mechanism in which repair events only occur while the gene is transcriptionally permissive. The key adjustment we make is the introduction of a higher transcription rate upon completion of repair. This modification is justified by our model selection process, as we find that the second-best model (model 4) also has an amplified transcription rate following Apex1 interaction and subsequent DNA repair. Interestingly, for the same value of *k*_*repair*_, transcription-coupled repair (model 5) leads to a less significant increase in noise and better maintenance of mean as compared to model 4 in which repair can also take place in the OFF state (model 4). This implies that coupling of repair to transcription is the most efficient method of DNA repair in terms of minimizing excess transcriptional variability.

Several molecular mechanisms can lead to the amplified transcription rate that we have incorporated into model 5. Apex1 binding may both momentarily silence RNA transcription while also inducing increased chromatin supercoiling and chromatin remodelling. This could lead to a subsequent increase in transcription efficiency through increased initiation and/or RNApol II processivity. This feedback mechanism may be perceived as a homeostatic process, allowing maintenance of the mean mRNA production despite a perturbation in template integrity. It is important to note that the dynamic binding and unbinding of Apex1 triggers noise enhancement and mean maintenance more than the repair per-se. One implication of this is that other protein-DNA dynamic interactions may lead to unavoidable noise modulation through structural constraints like supercoiling. The strength of such modulation will depend on the kinetic rates of interaction.

### 6 Sensitivity analysis of TCR model (Model 5)

#### 6.1 Modulation of *k*_*repair*_ and *k*_*incorpo*_ with fixed cooperativity

We have seen that a model incorporating Apex1 interaction with chromatin and an associated transcriptional amplification, can recapitulate an increase in noise without alteration of the mean number of mRNA produced by the Nanog gene expression system. We next asked how this behavior - increase in noise without dramatic modification of mean expression - is related to the dynamics of Apex1 binding and unbinding with chromatin. *k*_*repair*_ and *k*_*incorpo*_ are the kinetic rates describing such interaction.

We conducted a phase plane analysis of the system mean and Fano factor for both *k*_*repair*_ and *k*_*incorpo*_ (Fig S22A-B). We assume that the cooperativity is fixed and equal to the deduced cooperativity from the previous analysis using Nanog FISH data for 10*µM* IdU. As expected, the Fano factor increases as *k*_*incorpo*_ increases (for *k*_*incorpo*_ lower than ≈ 1). This suggests a positive dose-dependent relationship between IdU and noise (Fig S22B).

We observe an inverse relation for *k*_*repair*_, where noise increases as *k*_*repair*_ decreases. Experimentally, we use a small-molecule inhibitor of the Apex1 endonuclease domain (CRT0044876) to decrease *k*_*repair*_. It is interesting to highlight that when *k*_*incorpo*_ is higher than ≈ 1 the Fano factor starts to decrease. These observations can be understood looking at equation (36) for the Fano factor. For *k*_*incorpo*_ *>* 1, *ν*_*on*_ decreases slowly and the effective *k*_*RNA*_ starts to increase slowly as compared to when *k*_*incorpo*_ ∈ [0.1, 1] (Fig S22C-D). These changes are counteracted by a larger increase in *K*_*OFF*_. The behavior for the mean number of RNA produced with increasing *k*_*incorpo*_ *>* 1 can also be understood using the previous considerations: the decrease in mean corresponds to a decrease of the frequency in the ON state that is not counteracted by a strong enough cooperativity. All the results can be understood using the following formula for the Fano factor:

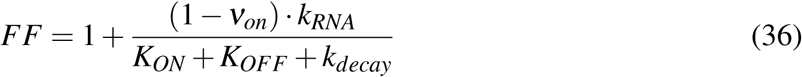

#### 6.2 Modulation of *k*_*ON*_ and *k*_*OFF*_ with fixed cooperativity

We next wanted to define the parameter regime for *k*_*ON*_ and *k*_*OFF*_ in which homeostatic maintenance of mean expression is possible with transcription-coupled repair (model 5). For this analysis, simulations were run with values of *k*_*ON*_, *k*_*OFF*_ ∈ [10^−3^, 10] for both the null model (standard 2-state model, DMSO condition) and model 5. For the same values of *k*_*ON*_, *k*_*OFF*_, the fold change in mean of mRNA counts was calculated by comparing results of model 5 to the null model. This provides insight into how IdU treatment may impact expression of genes with different bursting kinetics. When *k*_*OFF*_ *>> k*_*ON*_, the addition of IdU in Model 5 increases the average number of mRNA produced as compared to the null model (Fig S22E). This can be explained by a competition between the *OFF* and *ON*^∗^ states and by the fact that in this portion of the phase space *k*_*ON*_ *< k*_*OFF*_ *< k*_*incorpo*_ (and *k*_*incorpo*_ *< k*_*repair*_). Therefore, the probability of presence in the *ON*_2_ state (transcriptionally more productive state), increases. This implies that IdU treatment would increase the mean of very lowly expressed genes. This was seen experimentally in bulk RNA-seq measurements of transcript abundance in mESCs as the ≈ 100 genes that showed an increase in mean with IdU treatment were from the lowest expression regime (Fig S3). The exact inverse effect is observed in the upper left corner of the heatmap, where *k*_*ON*_ *> k*_*OFF*_ *> k*_*incorpo*_. Thus for highly expressed genes, the effect of IdU on mean expression is minimal as seen experimentally.

#### 6.3 Cooperativity and noise enhancement

When all the kinetics rates of the system are fixed, increasing the cooperativity, and thus the effective transcription rate, leads both to an increase in mean and Fano factor (Fig S22F).

### 7 Testing of TCR model for other noise-enhanced genes

In analyzing the Nanog gene expression we have found that IdU treatment leads to recruitment of DNA repair machinery while a gene is transcriptionally permissive (TCR model). This repair activity makes a second ON state accessible to the system. This state is characterized by an increased *k*_*RNA*_ which is sufficient to recapitulate the experimental observations of mean maintenance with increased noise strength (Fano factor). We next asked whether the inferred TCR model can explain the experimental data collected for other noise-enhanced genes within the scRNA-seq dataset of mESCs treated with 10*µM* IdU for 24 hours.

To simulate the TCR model for additional genes, estimates for the following parameters are needed: *k*_*ON*_, *k*_*OFF*_, *k*_*RNA*1_(basal transcription rate in DMSO), *k*_*decay*_, *and k*_*RNA*_ (effective transcription rate in IdU). To derive estimates of *k*_*ON*_, *k*_*OFF*_, *k*_*RNA*1_, *and k*_*RNA*_ we used a moments-matching technique described in (*(83, 84)*, where the first, second, and third exponential moments of the mRNA distributions in the DMSO and IdU conditions are used to calculate the parameters of a Poisson-beta distribution (describes 2-state model) that best fits experimental count data. The parameter estimates are derived in proportion to *k*_*decay*_. Values of *k*_*decay*_ were retrieved from an existing dataset of mRNA degradation rates in mESCs *(92)*.

Of the 945 genes classified as highly variable with IdU treatment, 314 genes remained for downstream analysis based on availability of mRNA degradation rates and high confidence parameter estimates (Fig. S24A). With the above parameter estimates, we next computed the constraints on the cooperativity term and *k*_*incorpo*_ as a function of *k*_*repair*_ using the relationships derived in sections 2.5 and 2.6. There is again one remaining degree of freedom in our model system: *k*_*repair*_. Using the MLE-based approach outlined in section 5.1, simulation results for a range of *k*_*repair*_ values were compared against scRNA-seq data to identify 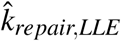 for each of the 314 genes.

Once 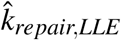 was identified, the macroscopic values of *K*_*ON*_ and *K*_*OFF*_ from simulation results were compared to experimentally derived estimates of *K*_*ON*_ and *K*_*OFF*_ from scRNA-seq data for each gene (Fig. S25). Overall, simulated values for macroscopic rates of promoter toggling in the IdU condition align with experimental results, indicating that the TCR model holds explanatory power for noise-enhanced genes beyond just Nanog. The use of a second, transcriptionally-enhanced ON state appears to be a unifying mechanism for maintenance of transcriptional homeostasis during DNA repair across a broad range of genes with different bursting kinetics. This suggests that the TCR mechanism for mean maintenance is robust to the initial bursting characteristics of a gene.

## Supplementary Figures S1-S27

**Figure S1:**
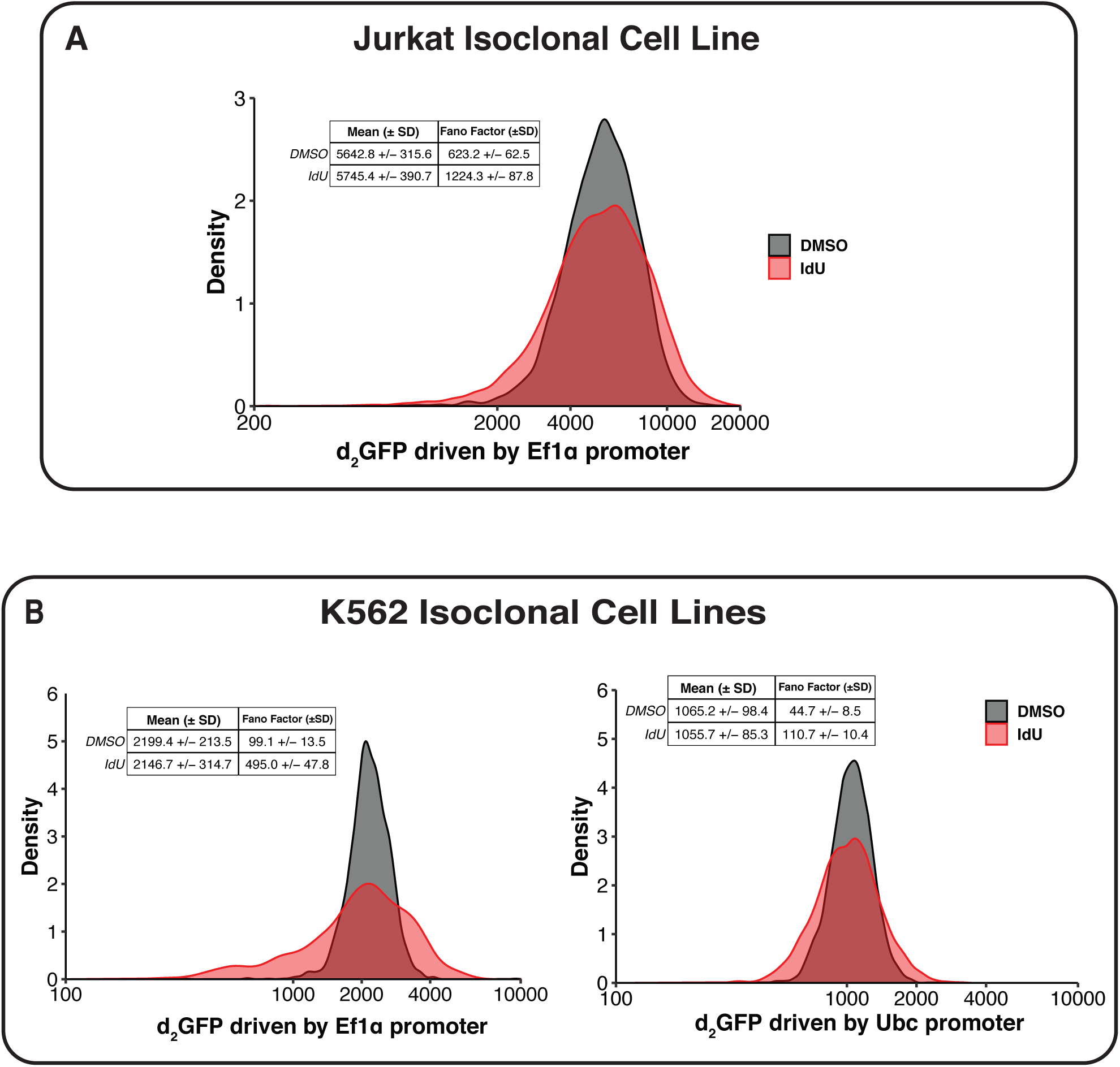
Nucleoside analog increases expression variability of housekeeping promoters in Jurkat and K562 cells. **(A)** Representative flow cytomtery distributions of d_2_GFP expression in an isoclonal population of Jurkat cells treated with either 20µM IdU or equivalent volume DMSO for 24 hours. Mean and SD are derived from 2 biological replicates. **(B)** Representative flow cytomtery distributions of d_2_GFP expression in isoclonal populations of K562 cells treated with either 20µM IdU or equivalent volume DMSO for 24 hours. Mean and SD are derived from 2 biological replicates.

**Figure S2:**
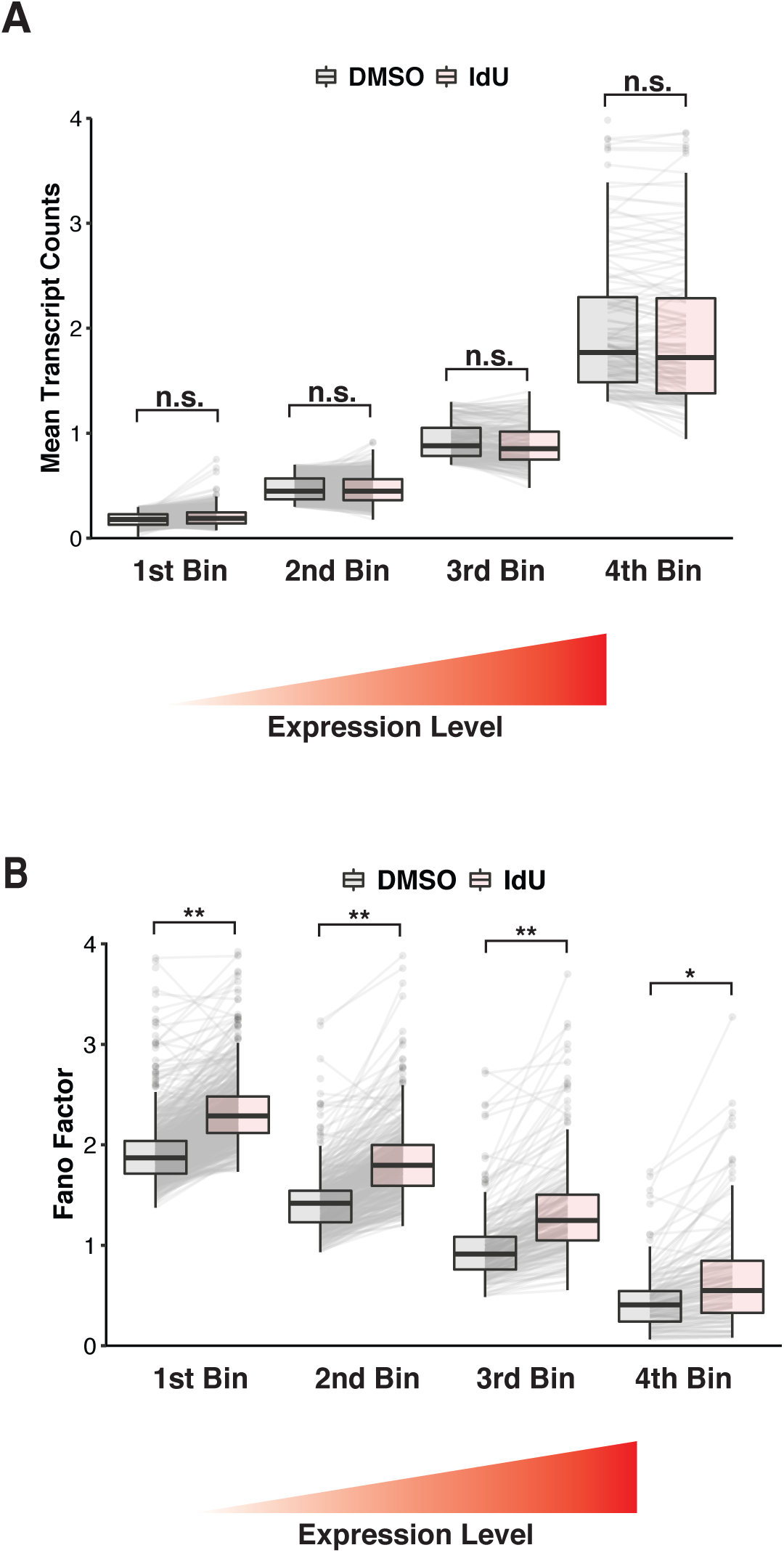
Noise enhancement occurs for genes across all expression levels. 4,578 genes from scRNA-seq dataset were binned into one of four groups (quartiles) based on mean expression level in DMSO condition. **(A)** Comparison of mean expression level for each gene in IdU and DMSO treatment groups. Boxplots show median *±* interquartile range of mean values for genes within each bin. Solid lines connect the same gene in the DMSO and IdU boxplots. P values were calculated using a two-tailed, paired Student’s t test. **(B)** Comparison of Fano factor for each gene in IdU and DMSO treatment groups. Boxplots show median *±* interquartile range of Fano factors for genes within each bin. Solid lines connect the same gene in the DMSO and IdU boxplots. P values were calculated using a two-tailed, paired Student’s t test. **p *<* 0.001, *p = 0.0016

**Figure S3:**
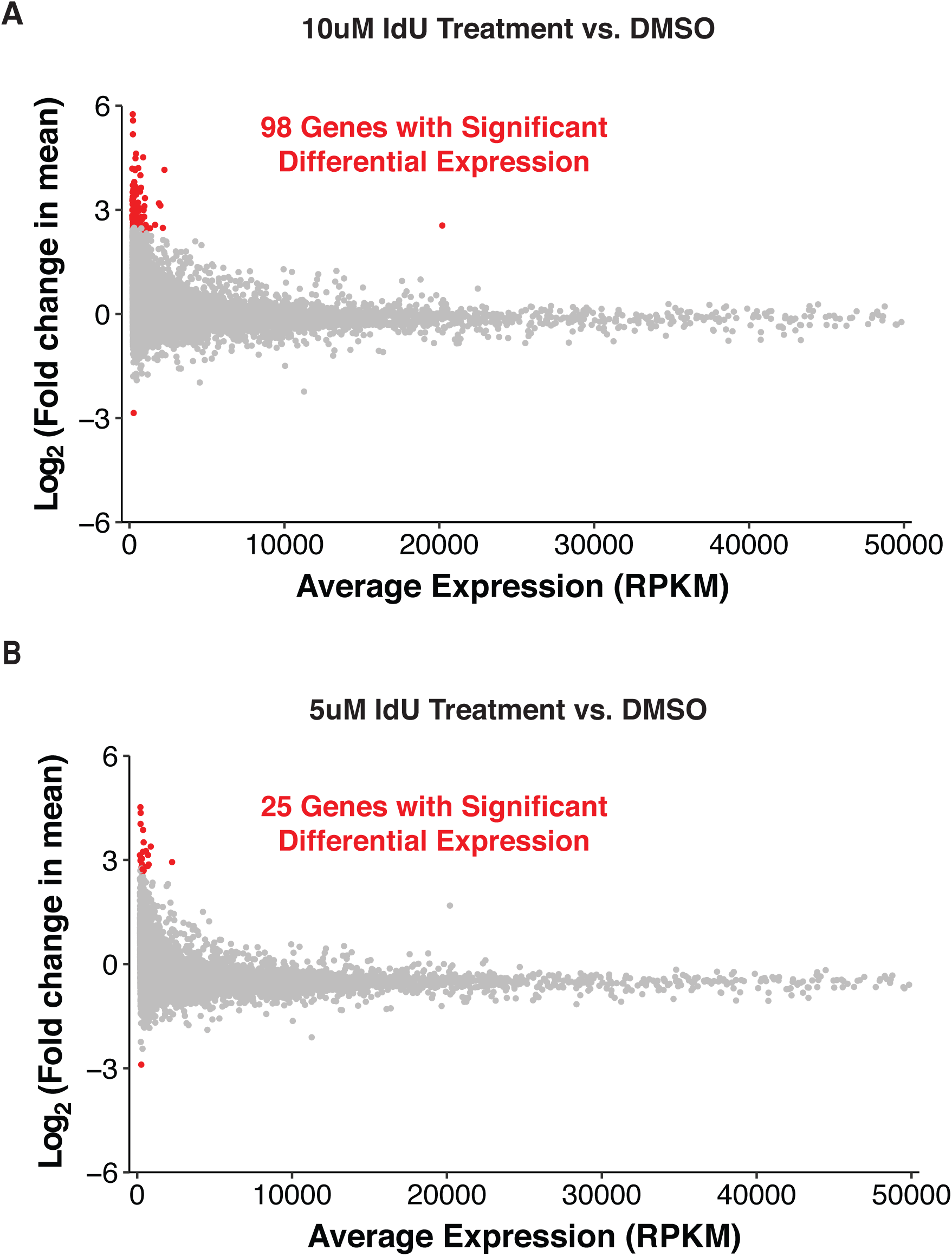
IdU causes minimal change in mean gene expression levels as measured by bulk RNA-seq. Transcript abundances were normalized using ERCC spike-in counts. Differential mean testing was conducted with a threshold of fold change *>* 2 and an FDR cutoff of 0.05. Genes considered differentially expressed are highlighted in red. **(A)** Mean transcript abundance vs. fold change (Log_2_) in mean for 12,502 genes. Comparison is between 10µM IdU and DMSO treatments. 98 genes were identifed as differentially expressed. **(B)** Mean transcript abundance vs. fold change (Log_2_) in mean for 12,054 genes. Comparison is between 5µM IdU and DMSO treatments.

**Figure S4:**
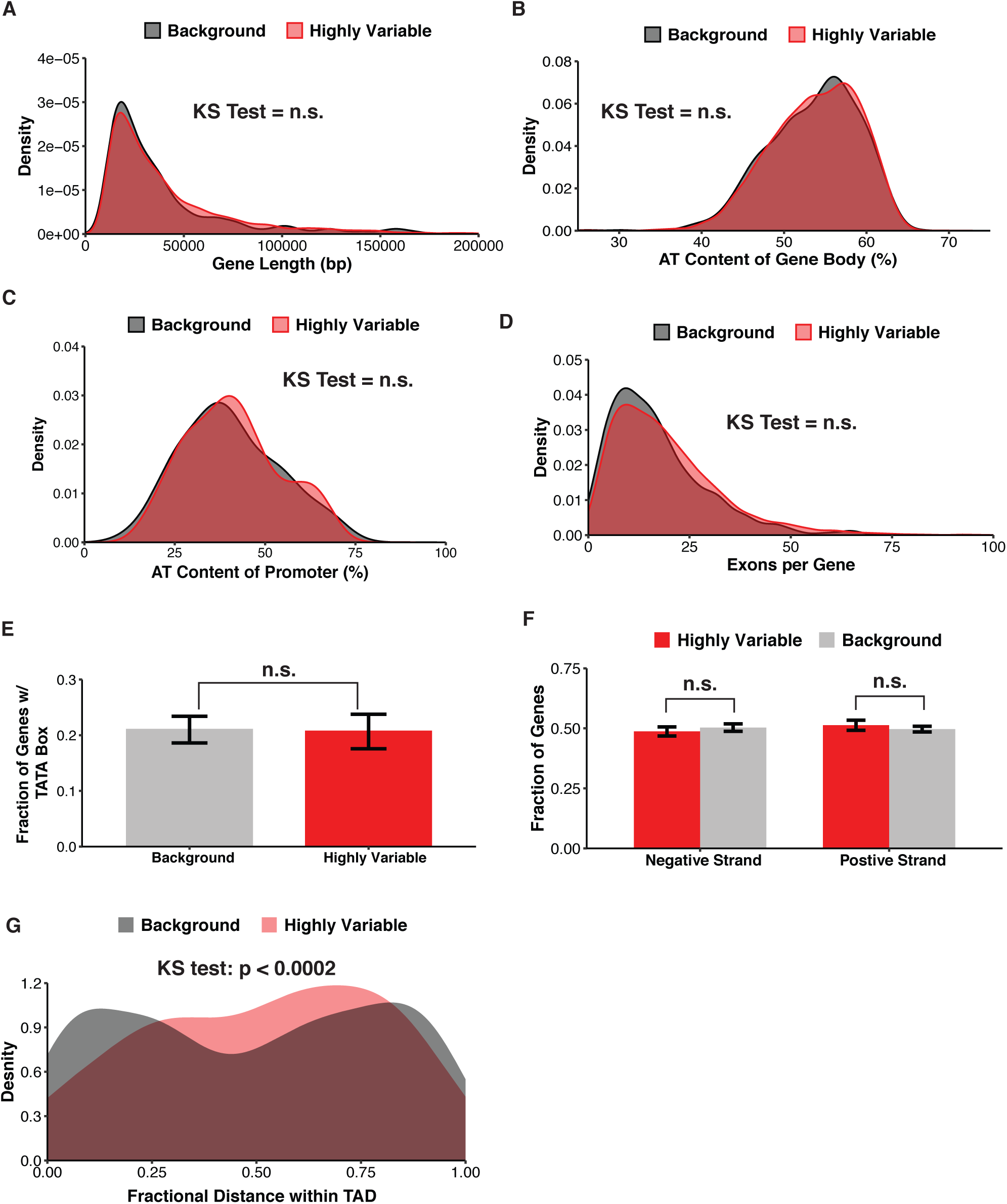
Noise-enhanced genes tend to be centrally located within topologically associating domains. Comparisons are between 945 genes classified as highly variable and 3,513 genes classified as non-variable (background) according to BASiCS algorithm. Gene characteristics and sequences were taken from Ensembl GRCm38 reference genome. **(A)** Distributions of gene lengths for highly variable and background genes. Length was calculated as distance between Ensembl gene start and end coordinates which correspond to outermost transcript start and end coordinates. **(B)** Percentage of base-pairs in gene body (based on gene start and end coordinates) that are A:T. **(C)** Percentage of base-pairs in 200bp region upstream of gene start that are A:T. **(D)** The number of exons was averaged over all transcripts associated with a gene. Distributions of average exon quantity for genes in the highly variable and background group were then plotted. **(E)** Fraction of genes with TATA sequence in 200bp region upstream of gene start. Data represent mean and SD from bootstrapping procedure with 10,000 resamplings of 100 genes from each group with replacement. **(F)** Fraction of genes whose coding sequence is located on negative and positive strands. Data represent mean and SD from bootstrapping procedure with 10,000 resamplings of 100 genes from each group with replacement. **(G)** Fractional distance of gene within TAD was calculated as (gene start coordinate -TAD start coordinate)/(TAD end coordinate - TAD start coordinate). Coordinates of TAD boundaries in mESCs were taken from Elphege et al. *(89)*.

**Figure S5:**
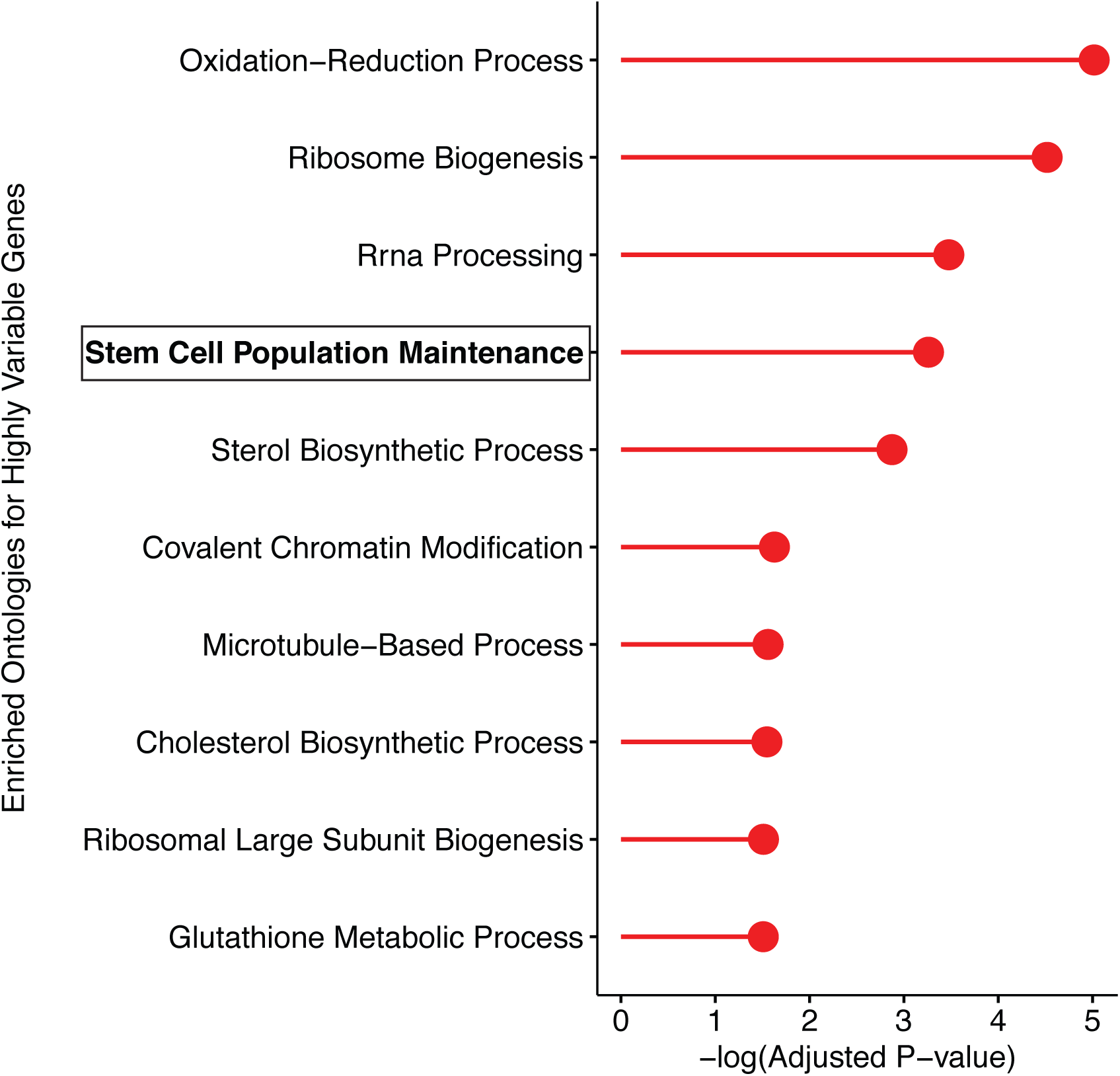
Ontology analysis of variably expressed genes shows enrichment for housekeeping and pluripotency maintenance pathways. DAVID v6.8 was used to identify enriched ontologies among the 945 genes classified as highly variable according to BASiCS algorithm.

**Figure S6:**
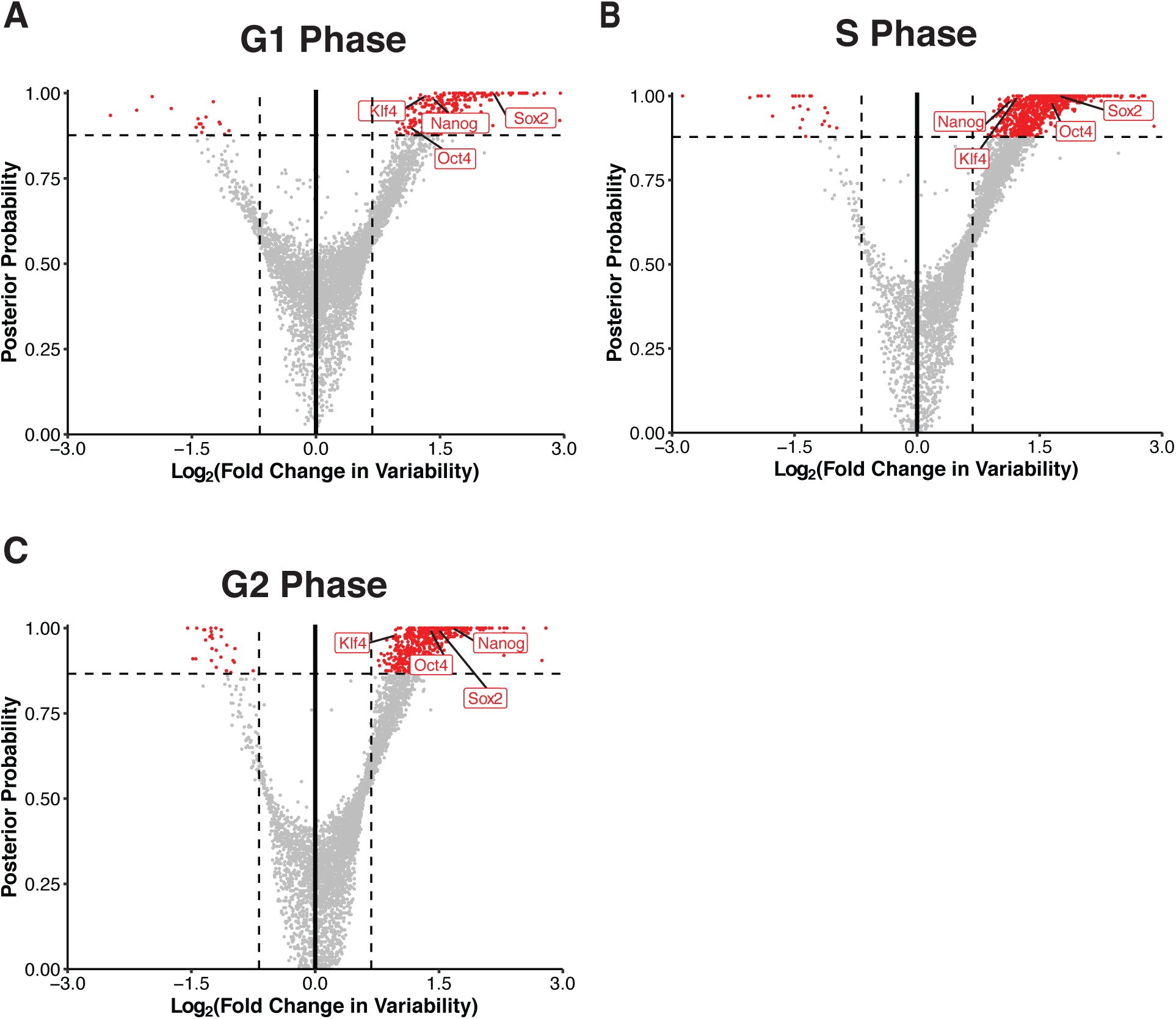
Noise-enhancement of pluripotency factors occurs in all three phases of the cell cycle. A total of 1556 cells in the scRNA-seq dataset were classified into one of three cell-cycle phases. Differential variability testing was then conducted between cells in the DMSO and IdU treatment groups with the same cycle classification.

**Figure S7:**
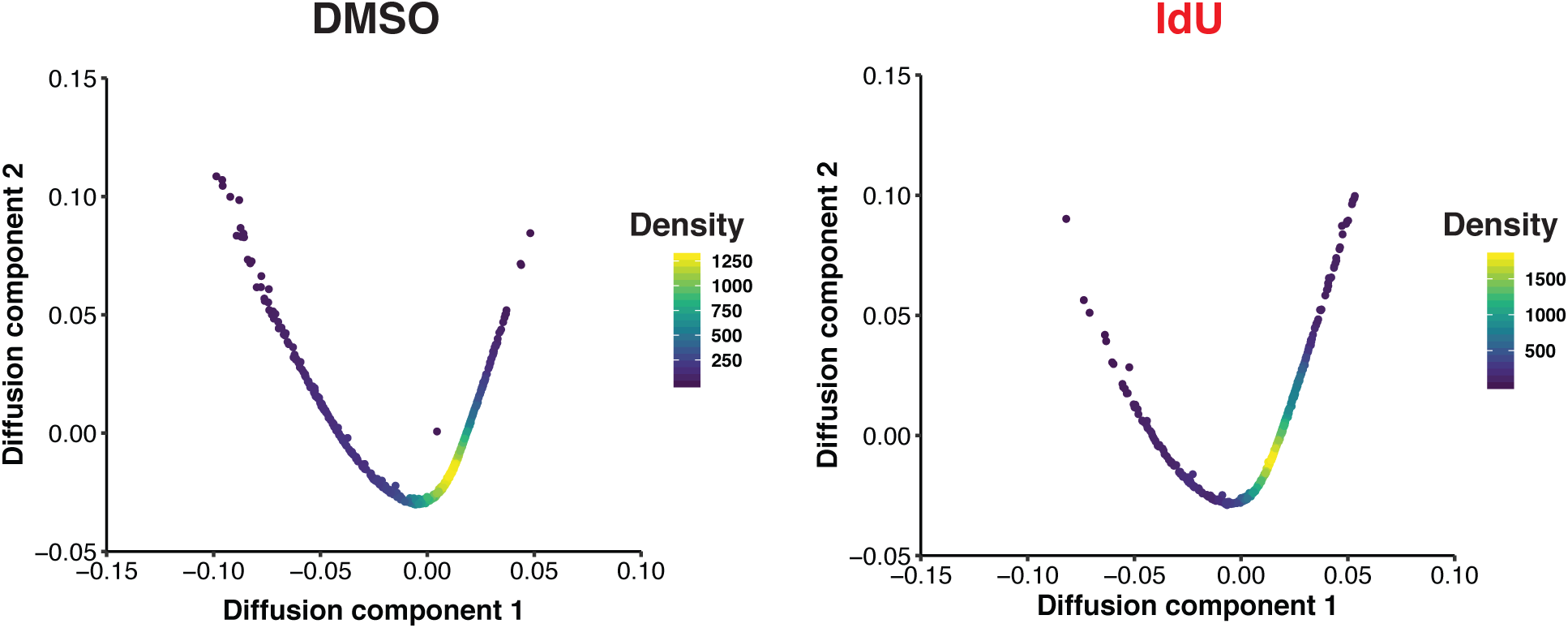
Transcript variability is not caused by bifurcation of mESCs into separate developmental lineages. Pseudotime analysis of IdU-treated cells shows no differentiation of mESCs into separate developmental lineages.

**Figure S8:**
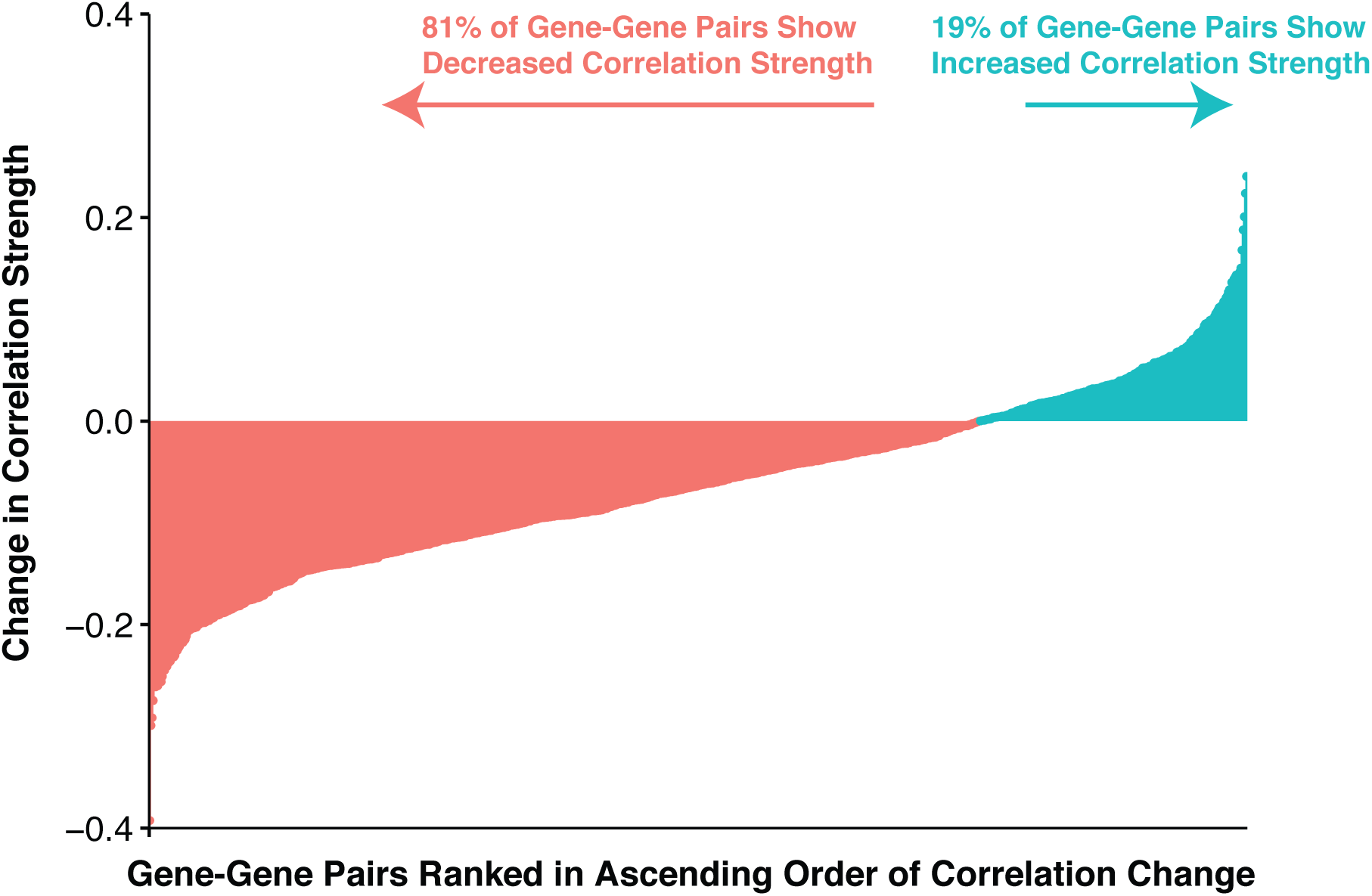
Majority of gene-gene pairs show a decrease in correlation strength. The Pearson correlation of expression for 923,521 (961 x 961) gene-gene pairs were compared between DMSO and IdU treatment groups. For each gene-gene pair, the absolute value of the correlation strength in DMSO was subtracted from the absolute value of the correlation strength in IdU. Negative values indicate loss of correlation in expression.

**Figure S9:**
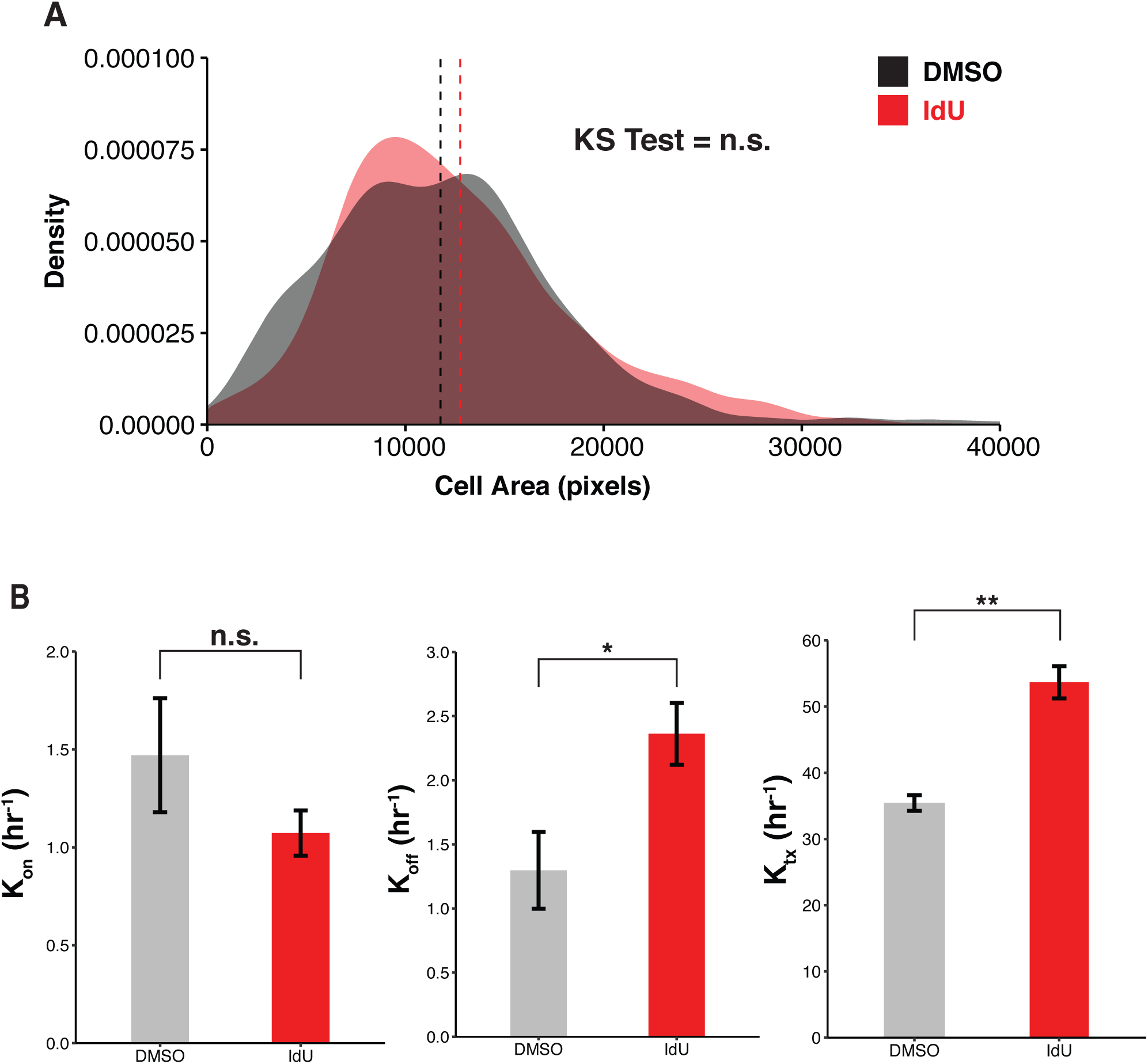
Shortened burst duration and increased transcription rate causes enhanced cell-to-cell variability in Nanog mRNA counts. **(A)** Distribution of cell sizes for analyzed cells in DMSO and IdU conditions. Cell size was calculated as number of pixels within segmented cell boundary. Dashed lines represent means of each distribution. Data represent pooling of cells from all four biological replicates for each condition. KS test shows no significant difference between cell size distributions. **(B)** Inference of parameters for 2-state model of transcription. P values were calculated using a two-tailed, unpaired Student’s t test. *p = 0.0017, **p = 0.0001

**Figure S10:**
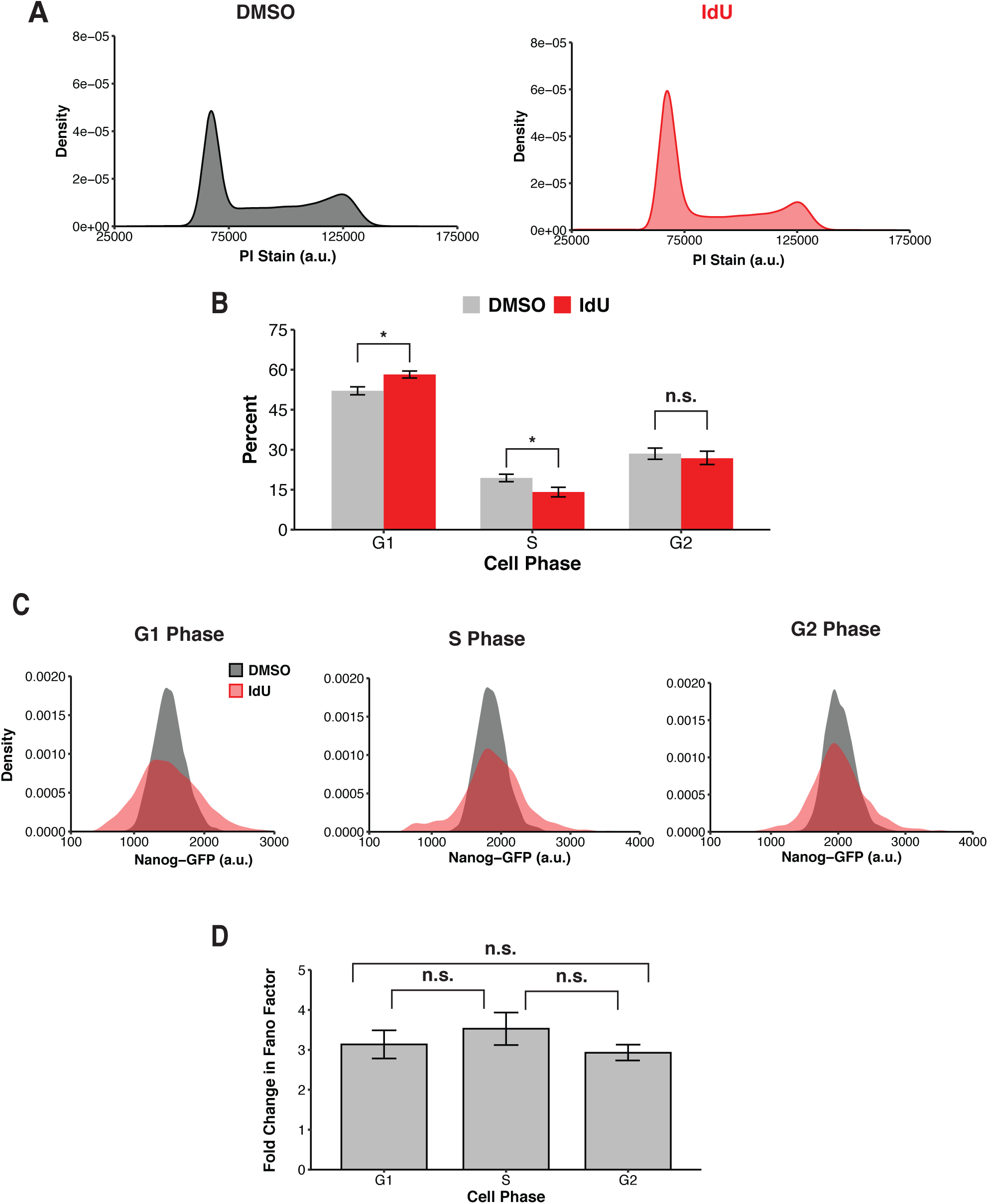
Noise-enhancement of Nanog protein expression is independent of cell-cycle state. **(A)** Representative flow cytometry distributions of propidium iodide staining for Nanog-GFP mESCs treated with either DMSO or 10µM IdU for 24 hours. No signs of aneuploidy are visible, indicating transcriptional variability is not due to cell-to-cell variability in gene copy numbers. **(B)** Percent of cells in each phase of the cell cycle for DMSO and IdU treatments based on propidium iodide staining. IdU treatment slightly slows entry into S phase. Data represent mean and SD of three biological replicates. P values were calculated using a two-tailed, unpaired Student’s t test. *p *<* 0.01 **(C)** Representative flow cytometry distributions of Nanog-GFP for mESCs within the G1, S and G2 phases of the cell cycle. mESCs were treated with 10µM IdU or equivalent volume DMSO for 24h followed by propidium iodide staining. **(D)** IdU-induced noise-enhancement of Nanog-GFP protein levels is unchanged across all three phases of the cell cycle. Nanog-GFP Fano factor with IdU treatment was normalized to DMSO control for calculation of fold change. Data represent mean and SD of three biological replicates. P values were calculated using a two-tailed, unpaired Student’s t test.

**Figure S11:**
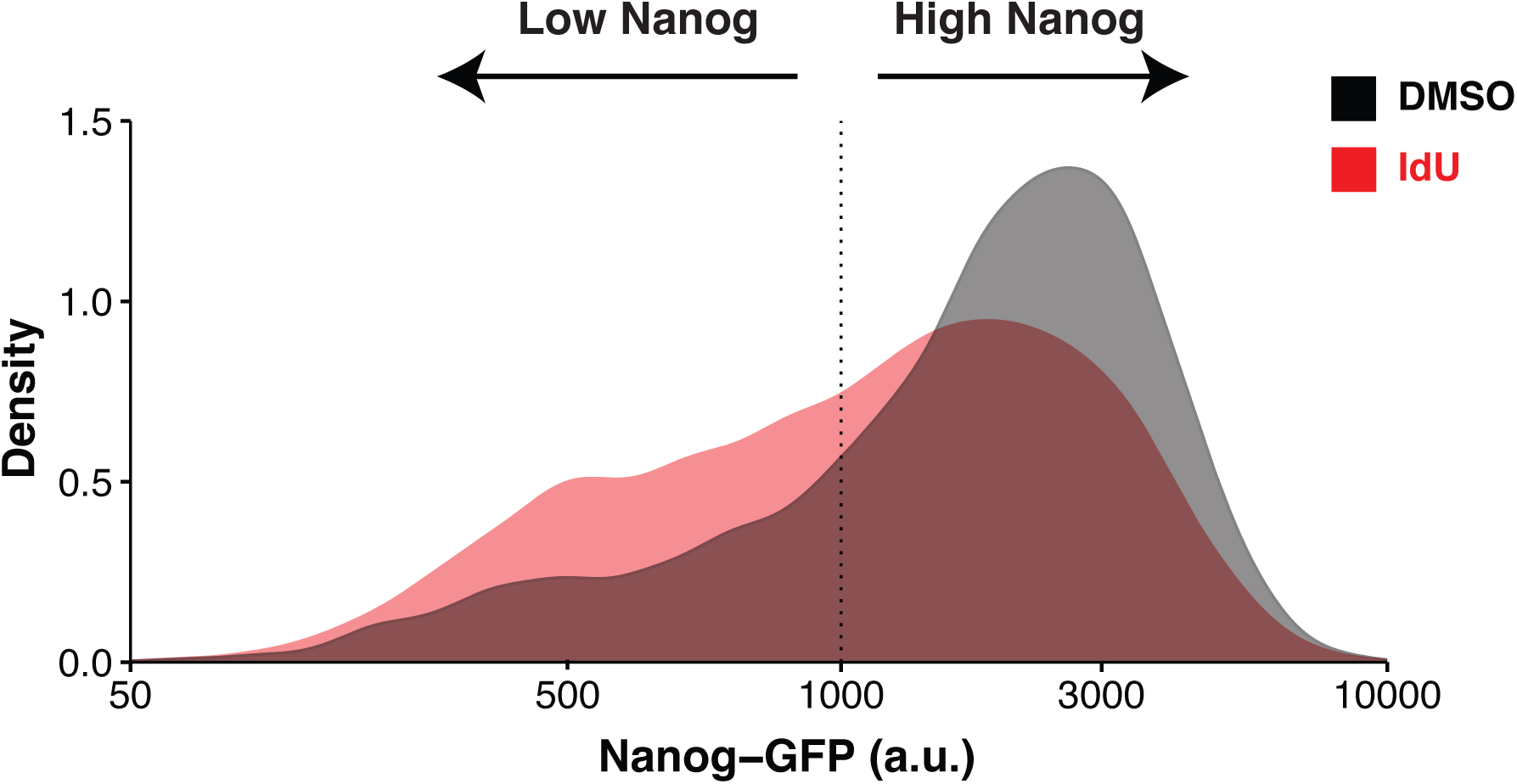
Increased transcriptional noise drives a greater number of mESCs into the low-Nanog state while cultured in serum/LIF. Flow cytometry distribution of Nanog-GFP expression for mESCs cultured in serum/LIF and treated with 10µM IdU or equivalent volume DMSO for 24h. Data is pooled from three biological replicates.

**Figure S12:**
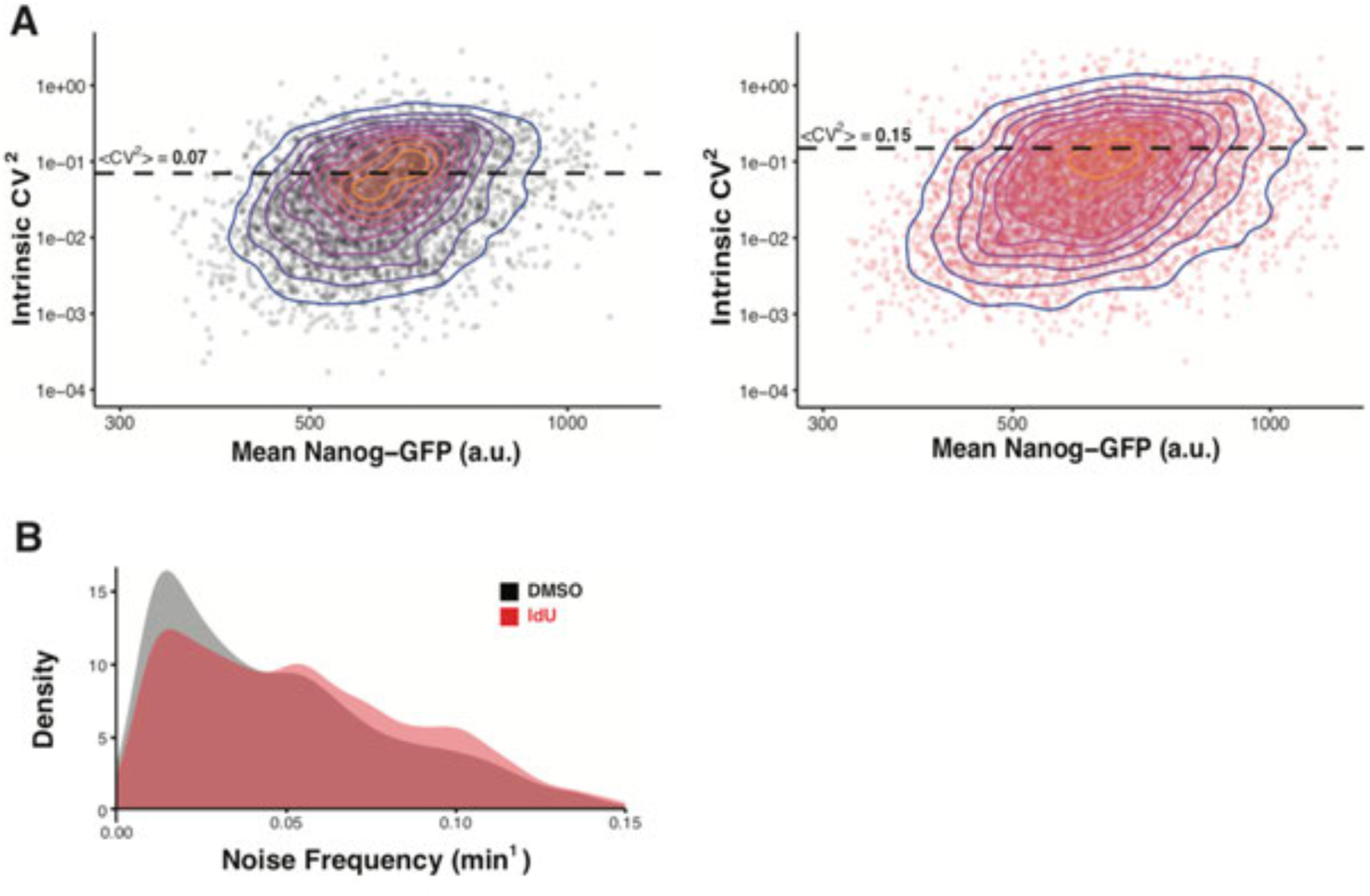
Time-lapse imaging demonstrates that altered kinetics of promoter toggling cause individual cells to experience larger fluctuations in Nanog protein expression. **(A)** Each point represents a single-cell fluorescence trajectory (DMSO on left, n = 1513; IdU on right, n = 1414). Single-cell fluorescence trajectories were detrended by subtracting time-dependent population average for Nanog-GFP fluorescence. The mean Nanog-GFP fluorescence for each raw trajectory is then plotted versus the CV^2^ of the detrended version of the trajectory to isolate intrinsic noise. The dashed lines represent the average intrinsic CV^2^ of all trajectories for each treatment group. Time-lapse imaging shows that for individual cells the magnitude of Nanog protein fluctuations increases with IdU treatment. **(B)** Distributions of noise frequencies from autocorrelation functions of each detrended trajectory. Noise frequency is calculated as the inverse of the autocorrelation time (*τ*_1*/*2_). Shorter but more productive transcriptional bursts with IdU treatment pushes the frequency content of Nanog-GFP fluctuations to higher spectra.

**Figure S13:**
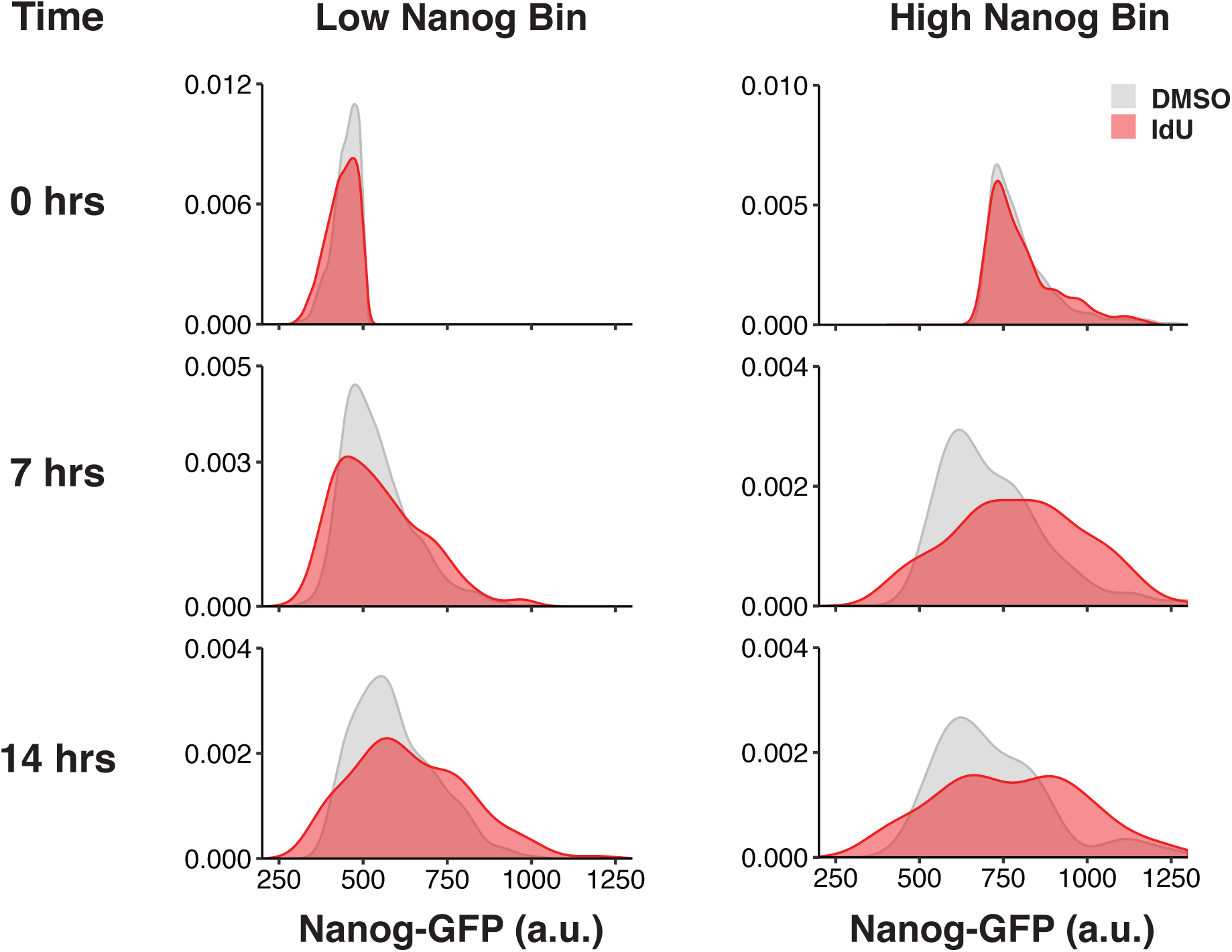
Amplification of expression fluctuations occurs independently of starting Nanog level. Single-cell trajectories whose starting fluorescence value was below 500 a.u. or above 700 a.u. were binned into low and high groups respectively. Only trajectories whose starting point coincided with addition of DMSO or IdU at time zero were used. Distributions of trajectory fluorescence values at zero, seven, and 14 hours into treatment conditions are shown. By 14 hours into IdU treatment, there is visible interconversion of cells between the low and high Nanog states, indicating that memory of initial Nanog expression level is erased. This precludes the possibility that noise enhancement is due to promoter mutations that create sub-populations of cells with stable expression of Nanog at low and high levels.

**Figure S14:**
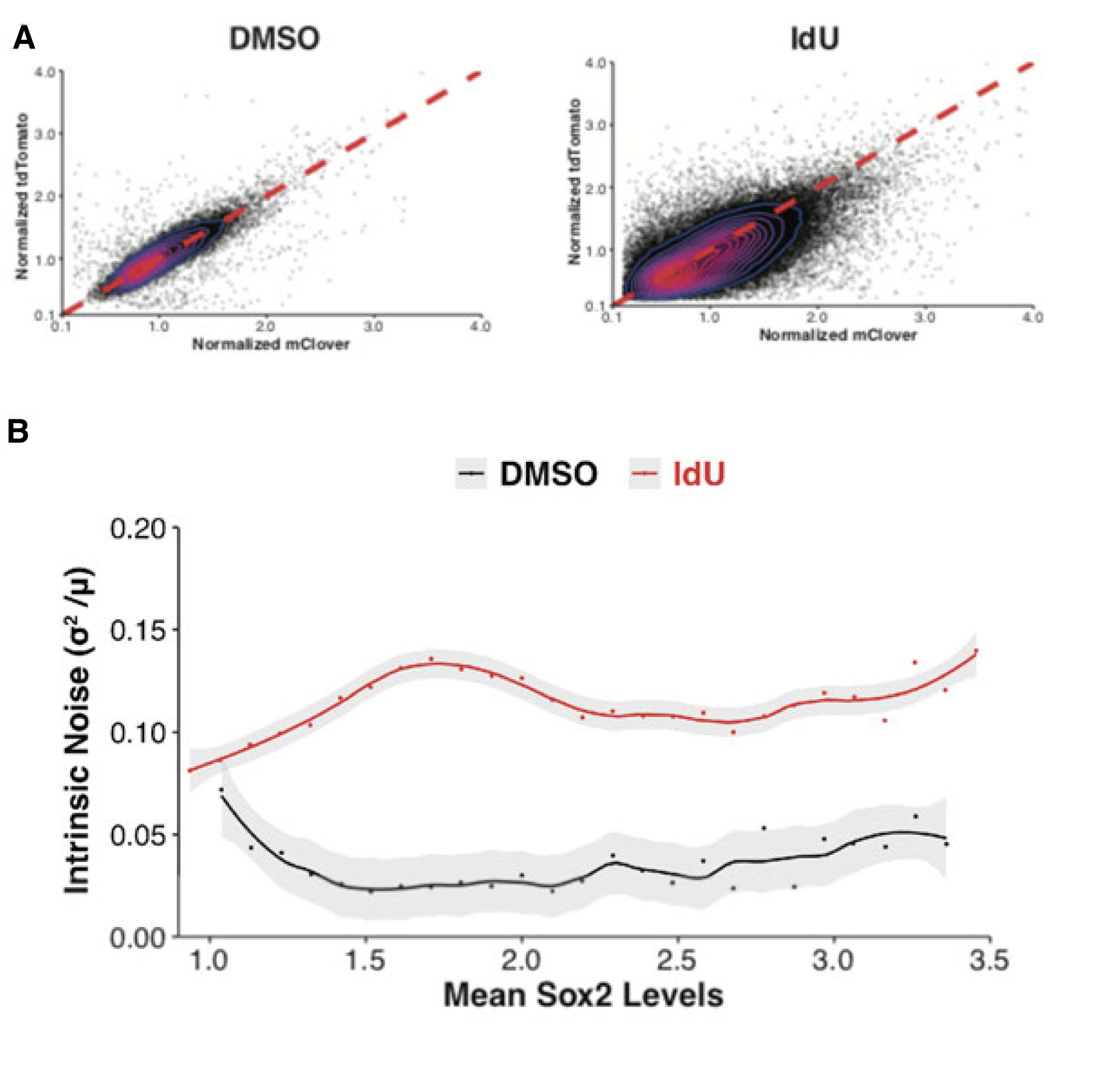
IdU treatment increases intrinsic noise of Sox2 expression. **(A)** Flow cytometry dot-plot of mESCs with Sox2 dual color tags. Dashed red line has slope of one. mCLover and tdTomato fluorescence values were normalized to population average. Data shown is pooled from three biological replicates. **(B)** Cells were binned according to total Sox2 expression from both alleles. Each point represents the intrinsic noise (Fano factor) of Sox2 expression for cells within a particular bin. Grey shadings represent 95% confidence intervals as determined by bootstrapping. Smooth lines are produced from loess regression. IdU increases Sox2 intrinsic noise across all expression levels.

**Figure S15:**
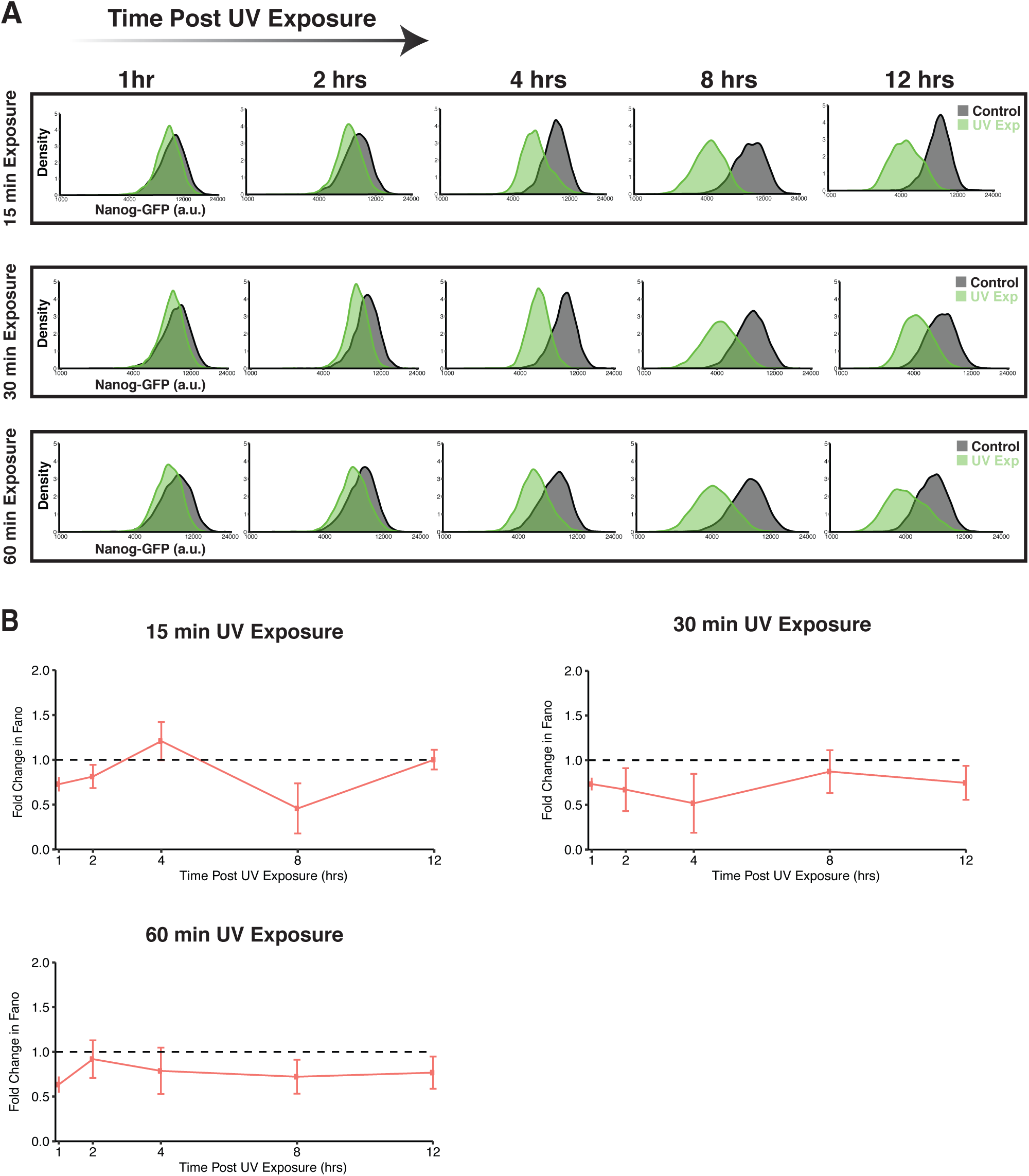
UV-stress reduces Nanog mean and Fano factor. **(A)** Representative flow cytometry distributions of Nanog-GFP expression from UV-exposed (green) and control (grey) cell populations. Cells were analyzed one, two, four, eight, and 12 hours post exposure. **(B)** For each exposure group (15, 30, and 60 minutes), the fold change in Fano factor is calculated as the Fano factor for Nanog in the UV-exposed population normalized to the Fano factor of its respective control population. Data points represent mean and SD of two biological replicates. Across all time points (except 4 hour point in 15 minute exposure group) UV stress reduces the Fano factor of Nanog.

**Figure S16:**
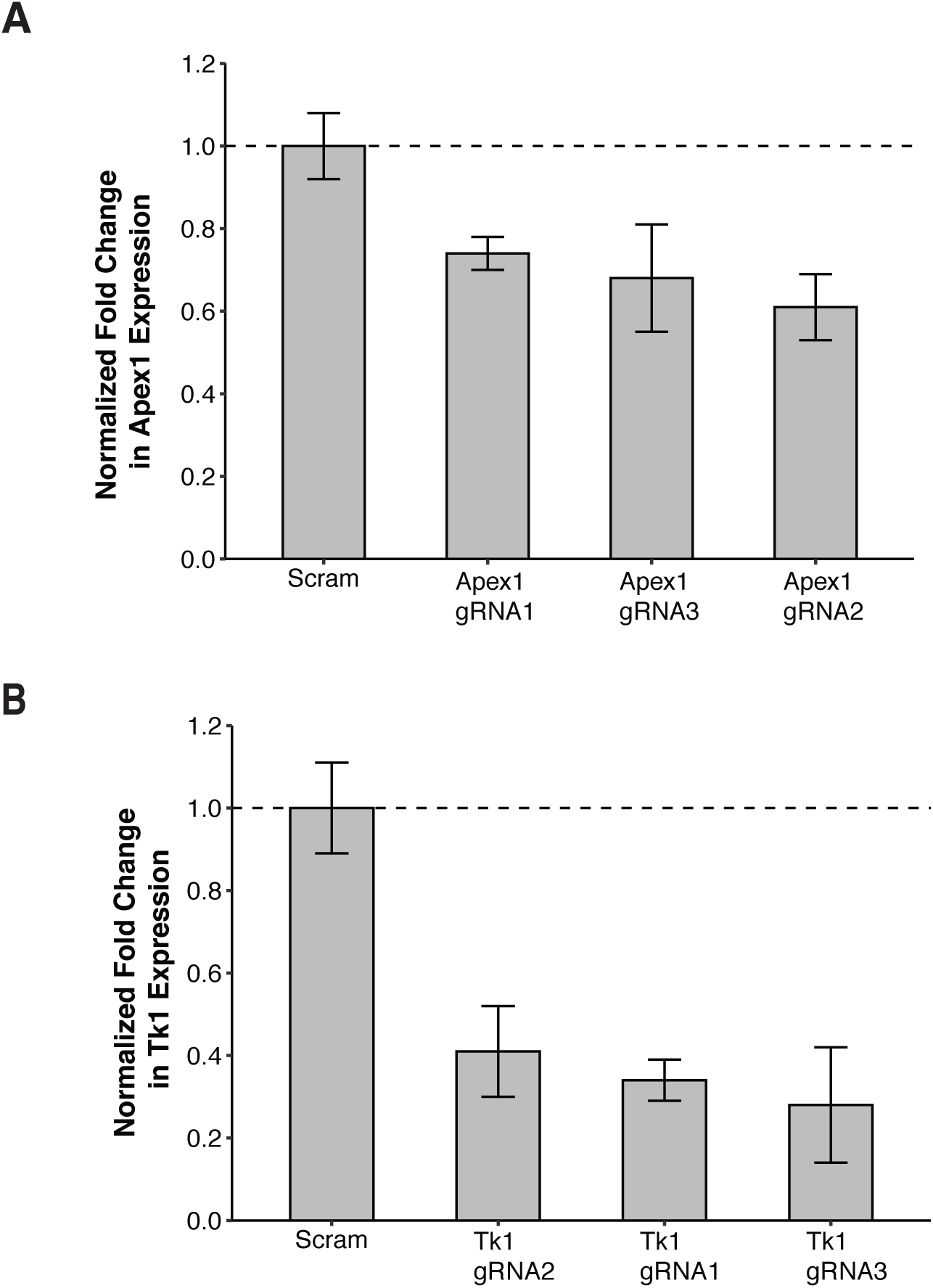
Validation of CRISPRi knockdown of Apex1 and Tk1 via qPCR measurements. ΔΔ*C*_*t*_ method was used with the empty-vector cell population as the control. Levels of Apex1 and Tk1 repression are relative to the non-targeting (scrambled) population. Data represent mean and SD of two biological replicates.

**Figure S17:**
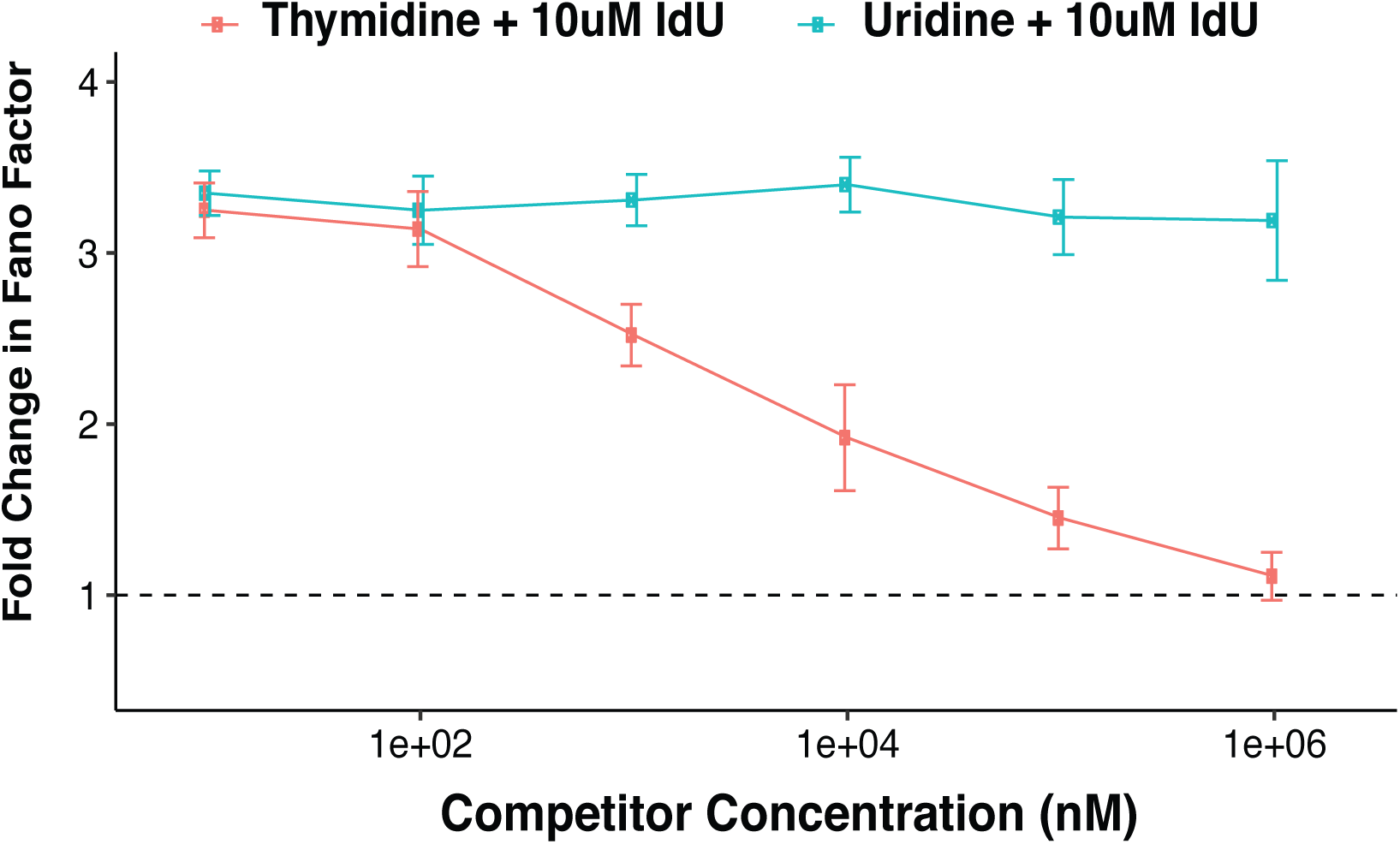
Thymidine competition ablates Nanog noise-enhancement from IdU. The Fano factor of Nanog for each concentration combination is normalized to DMSO control. For all treatment combinations, IdU concentration is held constant at 10µM. Concentration of thymidine (red) and uridine (blue) is reported on the x-axis. Combination of 100µM thymidine and 10µM IdU returns Nanog Fano factor to baseline level (DMSO control). Uridine, which is not a substrate of Tk1, fails to ablate IdU-induced noise-enhancement. Data points represent mean and SD of three biological replicates.

**Figure S18:**
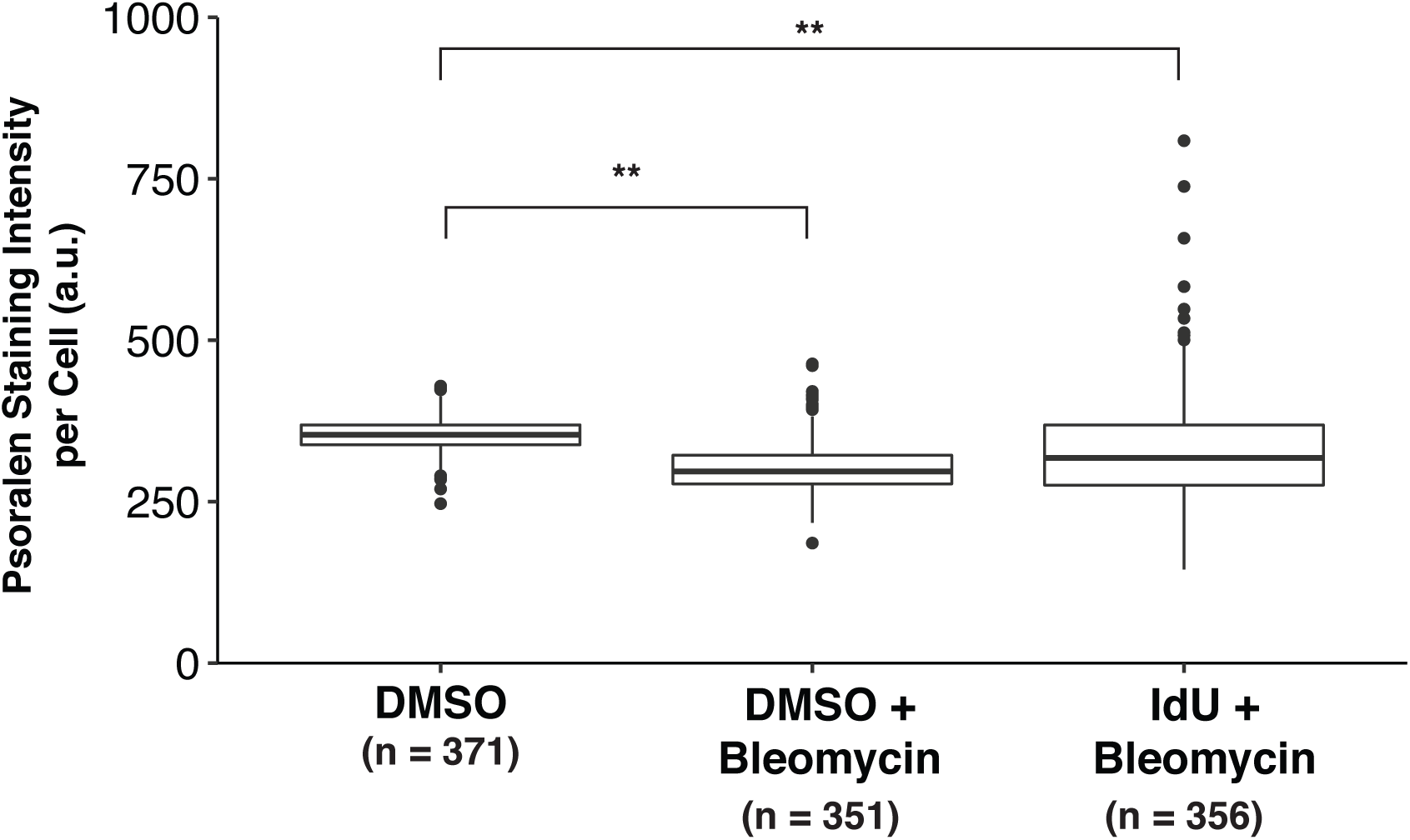
Bleomycin treatment reduces bTMP intercalation into DNA, validating assay sensitivity for negative supercoiling levels. Boxplots show median *±* interquartile range of single-cell bTMP staining intensities. Treatment of mESCs with 100µM bleomycin was performed for 1 hour just prior to bTMP incubation. Bleomycin reduces the mean bTMP staining intensity for cells treated with DMSO or 10µM IdU as compared to DMSO control with no bleomycin treatment (**p *<* 0.0001). The reduction in bTMP staining when IdU is coupled with bleomycin indicates that IdU alone in uncoiled DNA does not increase bTMP intercalation. Data shown are pooled from two biological replicates. P values were calculated using Kruskal-Wallis test followed by Tukey’s multiple comparison test.

**Figure S19:**
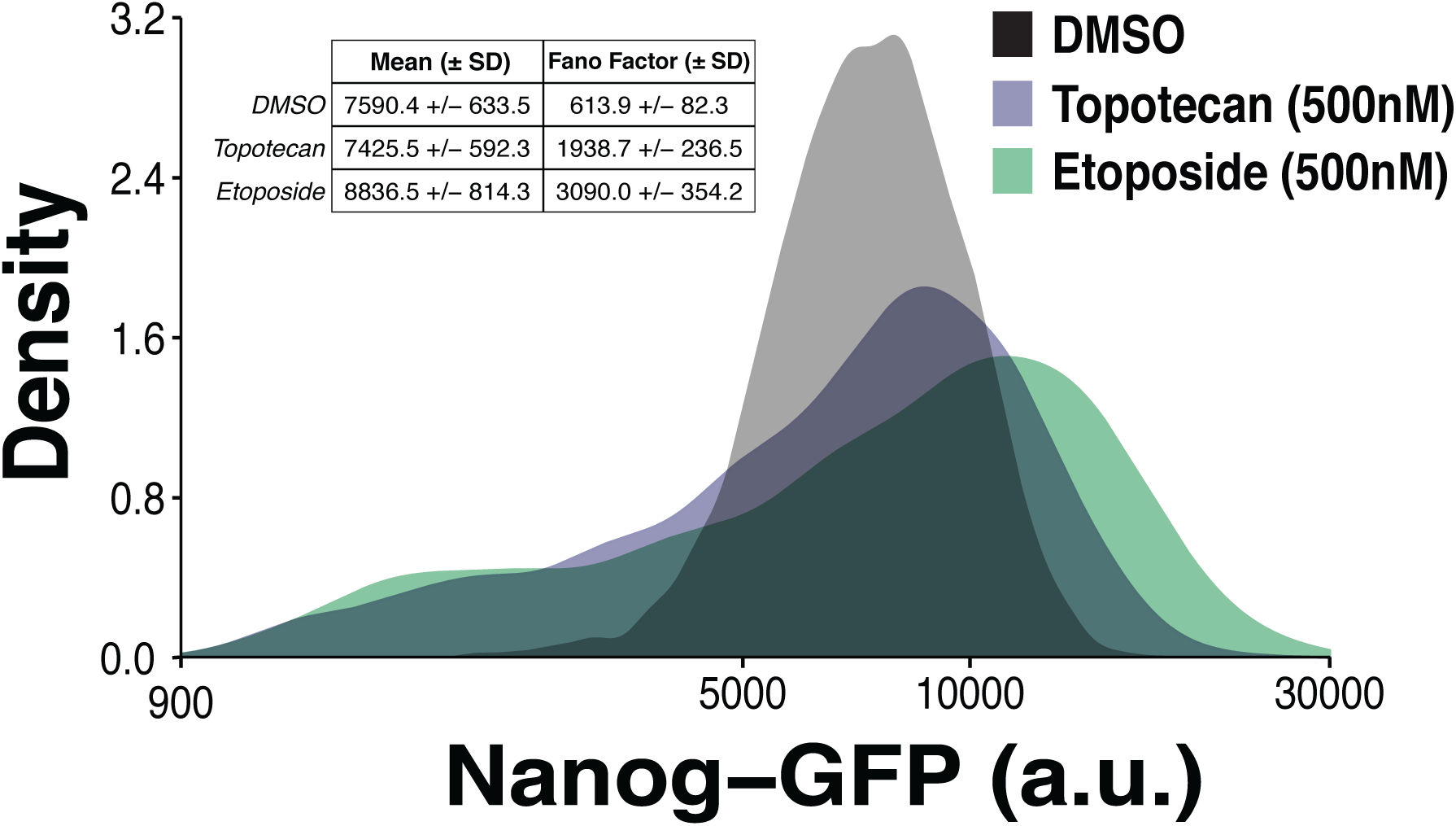
Small-molecule inhibition of Topoisomerase I and II increases Nanog expression variability. Representative flow cytometry distributions of Nanog-GFP expression in mESCs treated with DMSO, 500nM topotecan or 500nM etoposide for 24 hours in 2i/Lif media. Extrinsic noise filtering via cell-size gating was performed prior to calculation of Nanog Fano factor. Table inset shows mean and Fano factor (*±*SD) of Nanog expression averaged over three biological replicates of each treatment.

**Figure S20:**
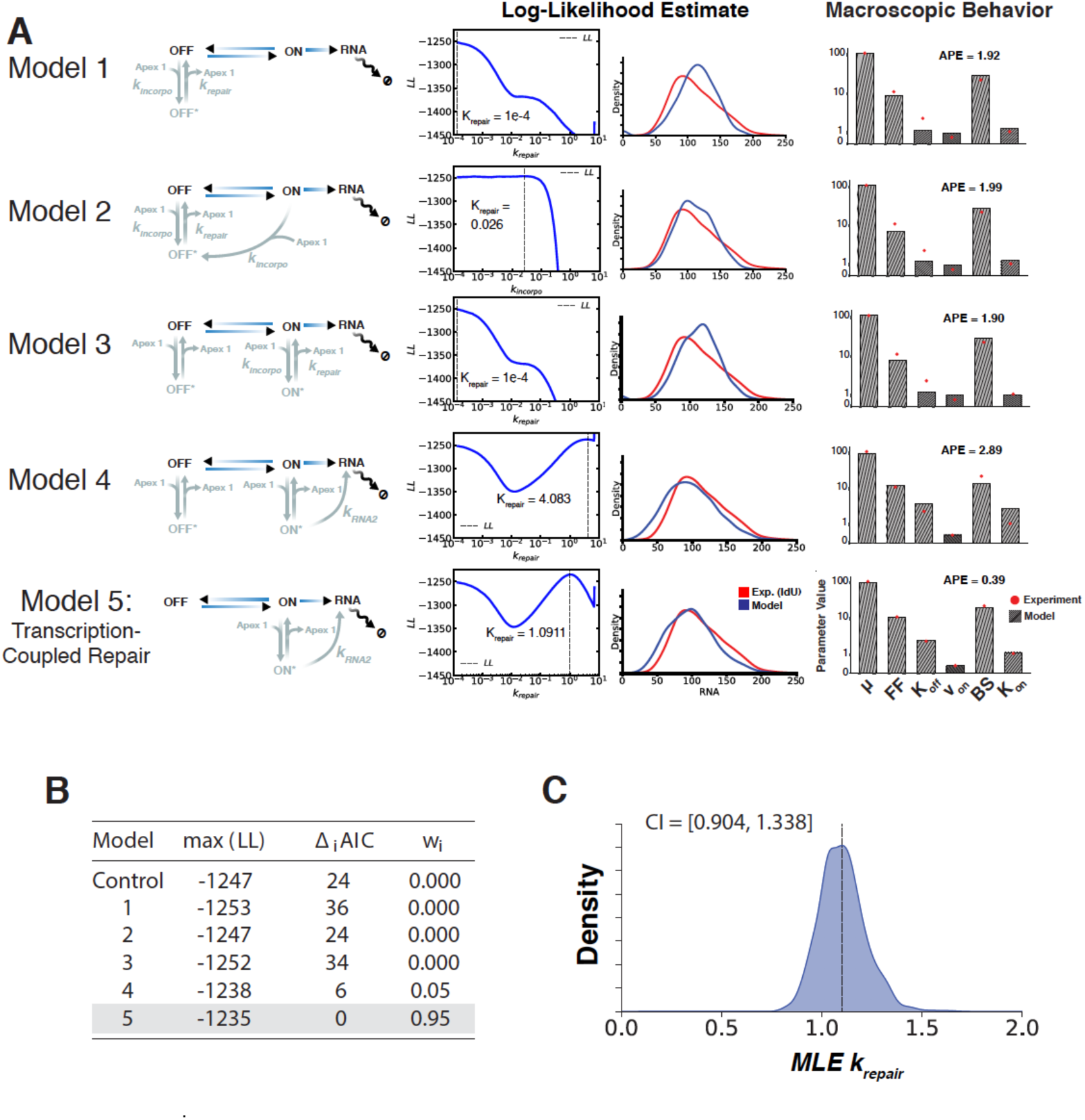
MLE-based approach for model selection reveals transcription-coupled repair mechanism best recapitulates experimental data. **(A)** (First Column) Schematic of simulated models incorporating Apex1 into standard 2-state model of transcription. (Second Column) For each model, 500 logarithmically-spaced values of *k*_*repair*_ ∈ [10^−4^, 10] were simulated. For each simulated value of *k*_*repair*_, log-likelihood is calculated as described in supplementary text 5.1.2 and plotted. Dashed vertical line in each plot denotes value of *k*_*repair*_ that maximizes log-likelihood estimate. (Third Column) Comparison of experimental Nanog mRNA distribution (red) to simulated distributions of Nanog mRNA (blue) for each model using value of *k*_*repair*_ that maximizes log-likelihood. (Fourth Column) Macroscopic behavior of simulation results (using value of *k*_*repair*_ that maximizes log-likelihood estimate) are compared to experimental data (supplementary text 5.2). Bars represent simulation values of Nanog gene expression system while red points with vertical line represent experimental data on Nanog expression from smRNA-FISH of mESCs treated with 10µM IdU. **(B)** For each tested model, the maximum log-likelihood value is listed along with the associated Δ_*i*_*AIC*. Model 5 (transcription-coupled repair) best describes experimental data based on these metrics. **(C)** Distribution and confidence interval (CI) of inferred *k*_*repair*_ values (based on MLE) for Model 5 using bootstrapping method in which the empirical distribution of Nanog mRNA counts from smRNA-FISH data was re-sampled 1000 times with replacement (supplementary text 5.1.4). Bootstrapping results show a well peaked distribution indicating practical parameter identifiability for *k*_*repair*_.

**Figure S21:**
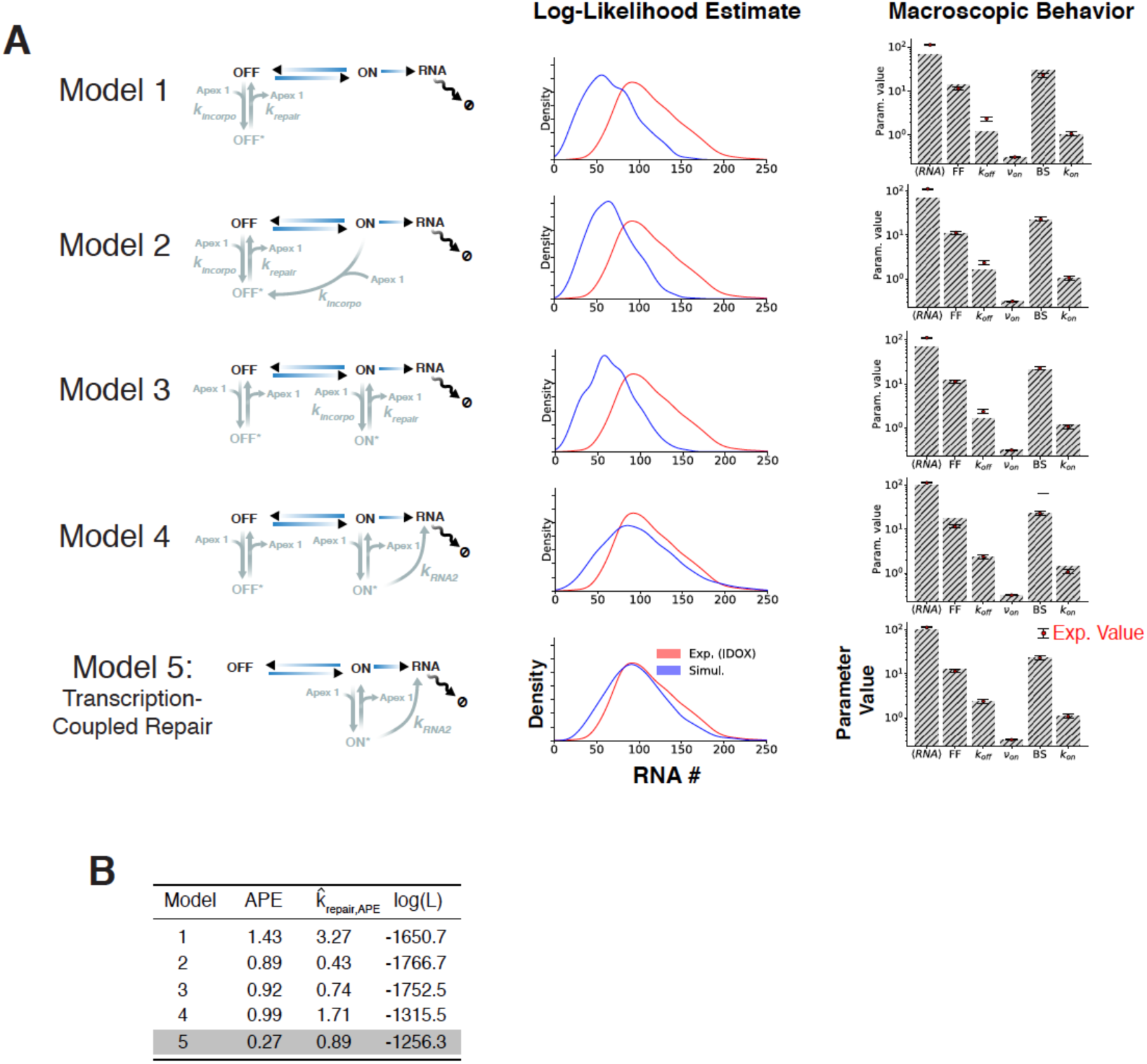
APE-based approach for model selection concurs with MLE-based approach, identifying TCR model as best match to experimental data. **(A)** (First Column) Schematic of simulated models incorporating Apex1 into standard 2-state model of transcription. (Second Column) Comparison of experimental Nanog mRNA distribution (red) to simulated distributions of Nanog mRNA (blue) for each model using value of *k*_*repair*_ that minimizes absolute percentage error. (Third Column) Macroscopic behavior of simulation results (using value of *k*_*repair*_ that minimizes absolute percentage error) are compared to experimental data (supplementary text 5.2). Bars represent simulation values of Nanog gene expression system while red points with vertical line represent experimental data on Nanog expression from smRNA-FISH of mESCs treated with 10µM IdU. **(B)** Values of *k*_*repair*_ that minimize the absolute percentage error for each model are listed. Model 5 (TCR model) yields the smallest APE and the largest log-likelihood.

**Figure S22:**
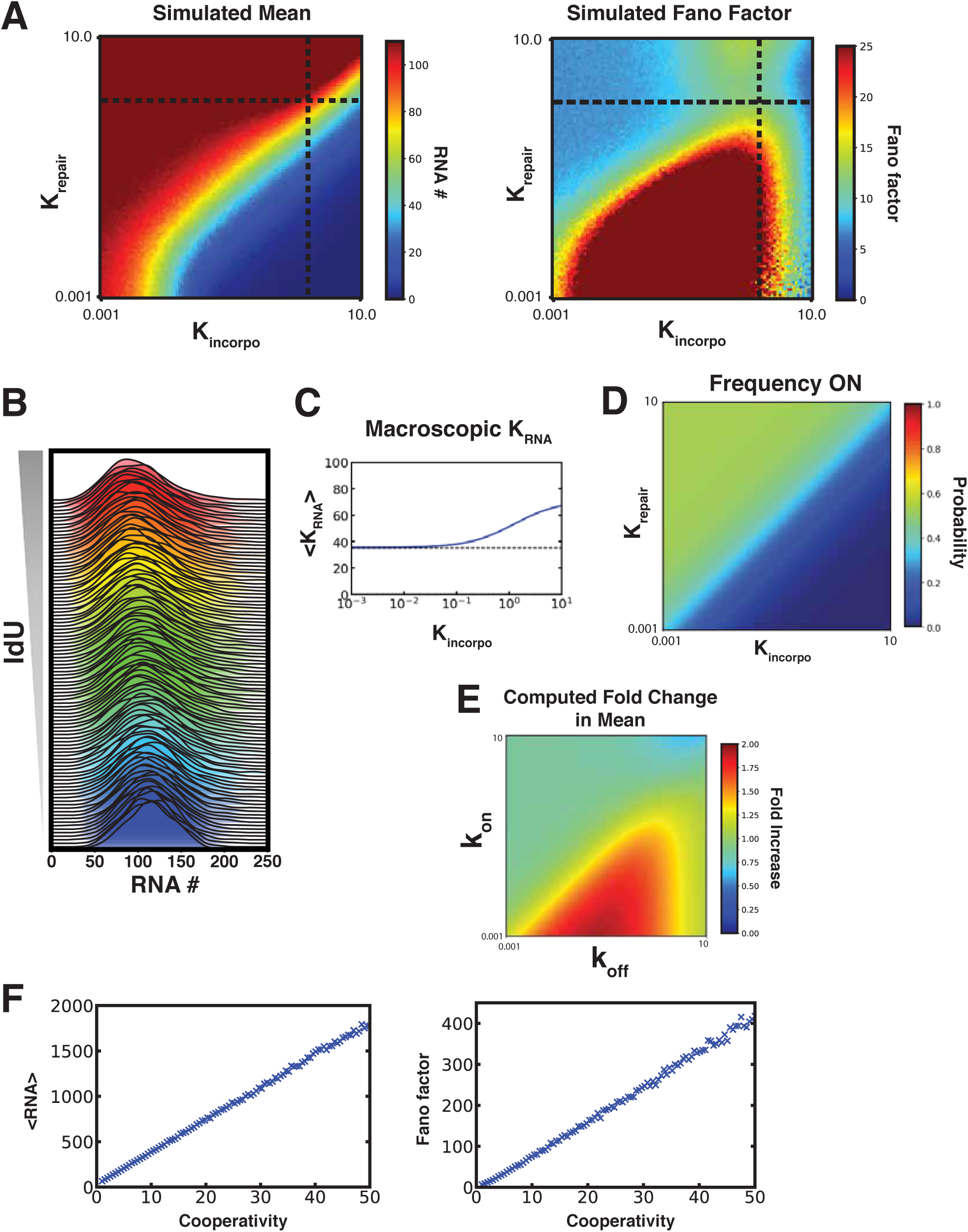
Sensitivity analysis of model parameters reveals phase-space for modulation of Nanog variability independently of mean. **(A)** Heatmaps displaying mean (left) and Fano factor (right) of Nanog mRNA from simulation results of TCR model as a function of *k*_*repair*_ and *k*_*incorpo*_ values spanning four orders of magnitude (supplementary text 6.1). Dashed horizontal and vertical lines represent inferred values of *k*_*repair*_ and *k*_*incorpo*_ that match experimental Nanog gene expression system in the presence of 10µM IdU. Multiple regions of the parameter phase-space exhibit constant mean output with unique levels of variability (Fano factor) demonstrating how mean and variability are tuned independently. **(B)** Simulated distributions of Nanog mRNA with increasing concentration of IdU which increases *k*_*incorpo*_. Simulation results demonstrate how TCR model allows for maintenance of mean output with increasing variability as concentration of IdU is increased. **(C)** Effective transcription rate of Nanog gene expression system as a function of *k*_*incorpo*_. As IdU incorporation and subsequent Apex1 recruitment increases, the effective transcription rate increases as well. This represents the compensatory mechanism of model 5 allowing for maintenance of mean output with increasing incorporation of IdU. **(D)** Heatmap displaying fraction of time that the Nanog gene expression is in the macroscopic ON state as a function of *k*_*repair*_ and *k*_*incorpo*_ values. **(E)** Heatmap displaying fold change in mean as a function of microscopic *k*_*o f f*_ and *k*_*on*_ values (supplementary text 6.2). Fold change is calculated as the output of Model 5 relative to model 0 (canonical 2-state model) for the same set of *k*_*o f f*_ and *k*_*on*_ values. For a gene whose *k*_*o f f*_ >> *k*_*on*_, addition of IdU to the system increases the mean output. **(F)** Mean mRNA and Fano factor of Model 5 output as a function of the cooperativity term which describes how strongly the transcription rate is amplified following completion of repair (supplementary text 6.3).

**Figure S23:**
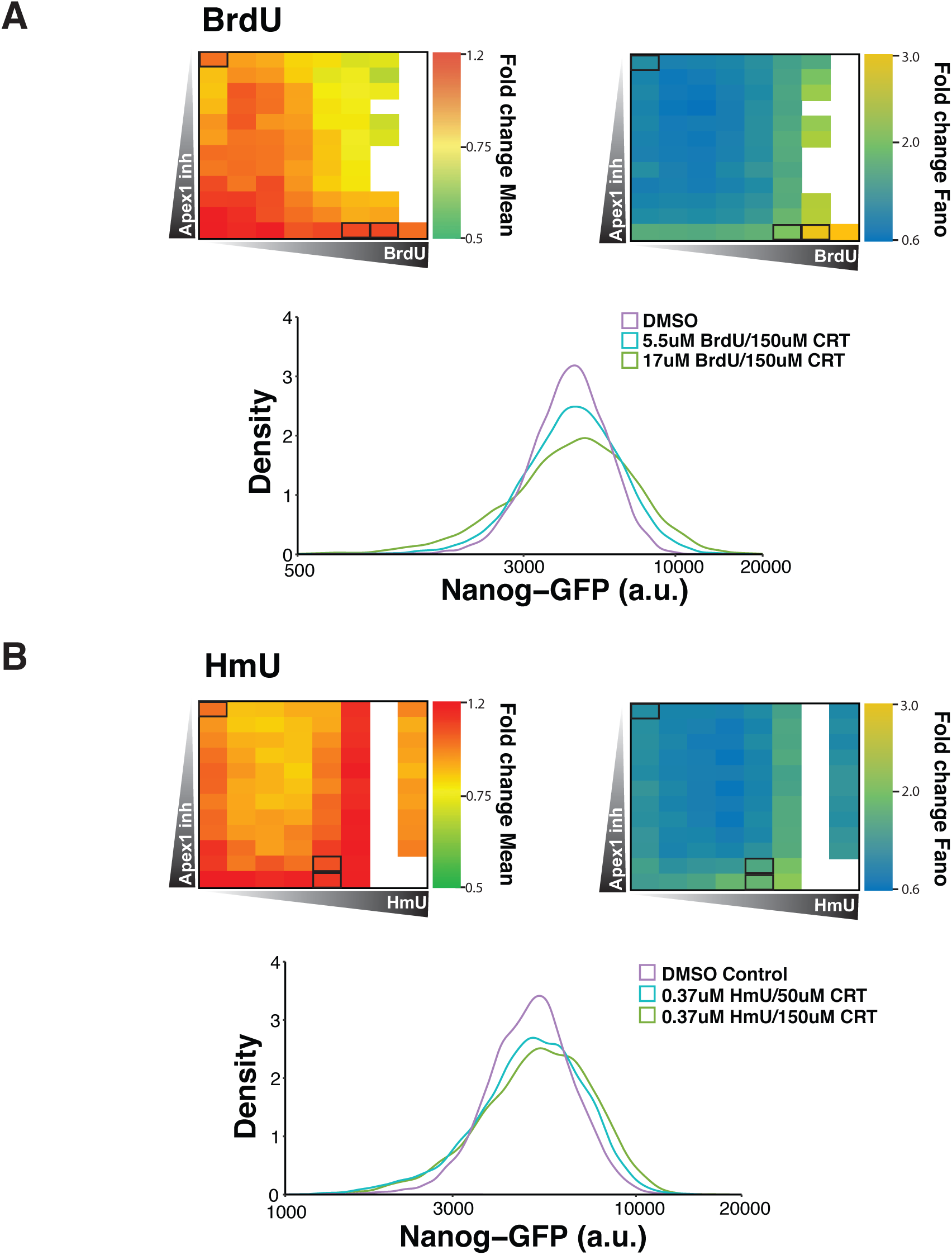
Treatment with BrdU or HmU in combination with CRT0044876 allows for tuning of Nanog variability independently of the mean. **(A)** Testing of 96 concentration combinations of BrdU and CRT0044876 (apex1 endonuclease domain inhibitor) to validate tunability of Nanog variability. BrdU and CRT0044876 were used to increase binding and decrease unbinding of Apex1 respectively. Nanog-GFP mESCs grown in 96-well plates were treated with 12 concentrations of CRT0044876 ranging from 0 to 150uM in combination with 8 concentrations of BrdU ranging from 0 to 50uM. Data represent average of two biological replicates. (Top left and top right panels) 96-well heatmaps displaying fold change in Nanog mean and Fano factor for each drug combination as compared to DMSO (top-leftmost well). Insufficient number of cells (*<*50,000) for extrinsic noise filtering were recorded from white wells. (Bottom Panel) Representative flow cytometry distributions from highlighted wells (black rectangles). Nanog variability increases independently of the mean. **(B)** Testing of 96 concentration combinations of HmU and CRT0044876 (apex1 endonuclease domain inhibitor) to validate tunability of Nanog variability. HmU is a naturally found, Tet-induced oxidation product of thymine. Nanog-GFP mESCs grown in 96-well plates were treated with 12 concentrations of CRT0044876 ranging from 0 to 150uM in combination with 8 concentrations of HmU ranging from 0 to 10uM. Data represent average of two biological replicates. (Top left and top right panels) 96-well heatmaps displaying fold change in Nanog mean and Fano factor for each drug combination as compared to DMSO (top-leftmost well). Insufficient number of cells (<C50,000) for extrinsic noise filtering were recorded from white wells. (Bottom Panel) Representative flow cytometry distributions from highlighted wells (black rectangles). As with IdU and BrdU, Nanog variability increases independently of the mean.

**Figure S24:**
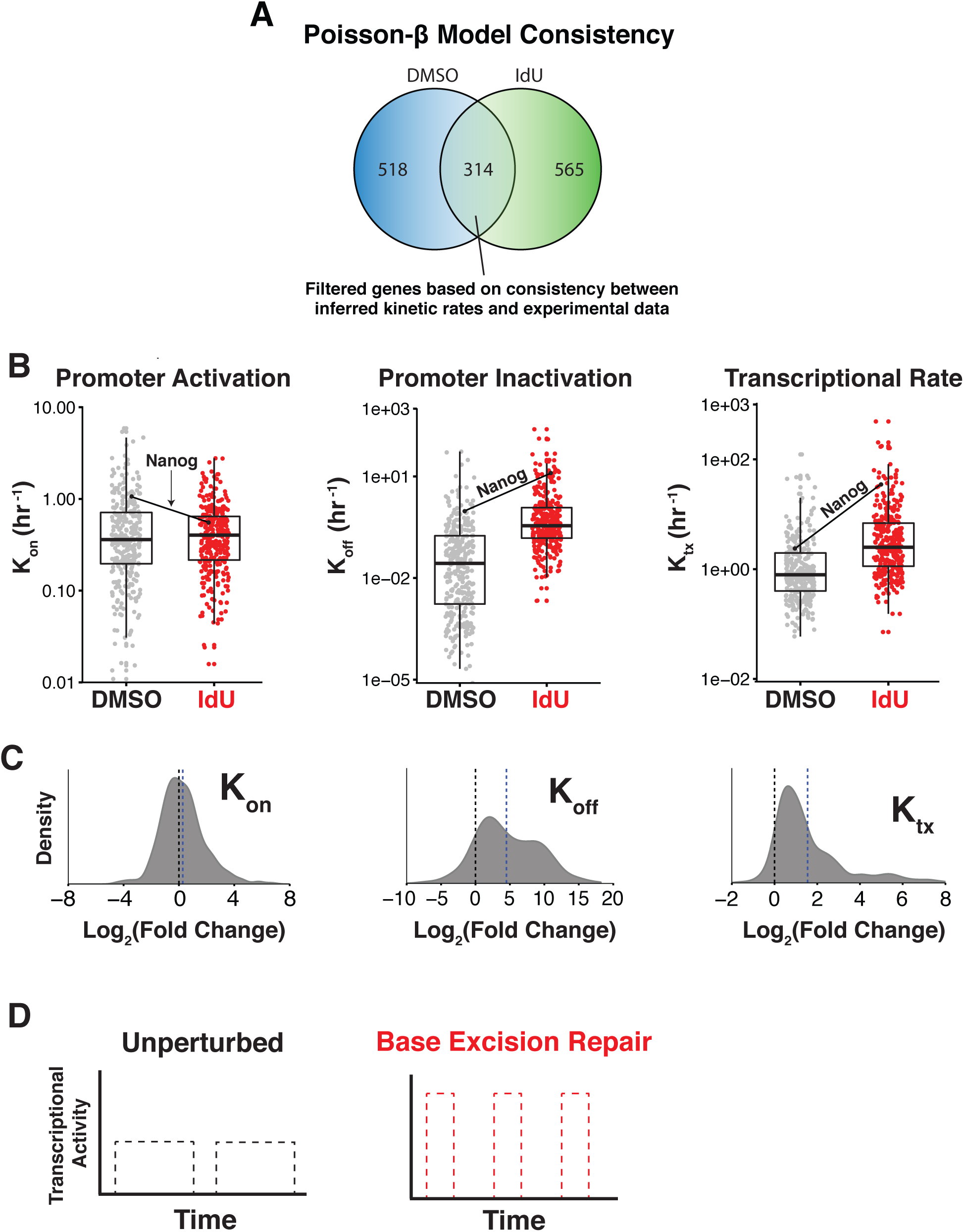
Highly variable genes exhibit shorter but more intense transcriptional bursts. **(A)** Of the 945 genes classified as highly variable with IdU treatment, we were able to estimate parameters of the 2-state model for 314 of these genes (supplementary text 7). **(B)** Boxplots show median ±interquartile range of parameter estimates with each point representing a gene. **(C)** Distributions of fold change in bursting kinetics between IdU and DMSO conditions for 314 highly variable genes. Dashed blue line signifies mean of distribution. Majority of highly variable genes exhibit increased *K*_*OFF*_ and *K*_*tx*_, which is consistent with TCR model. **(D)** Base-excision repair orchestrates shorter but more intense transcriptional bursts to maintain mean expression for genes with diverse bursting kinetics.

**Figure S25:**
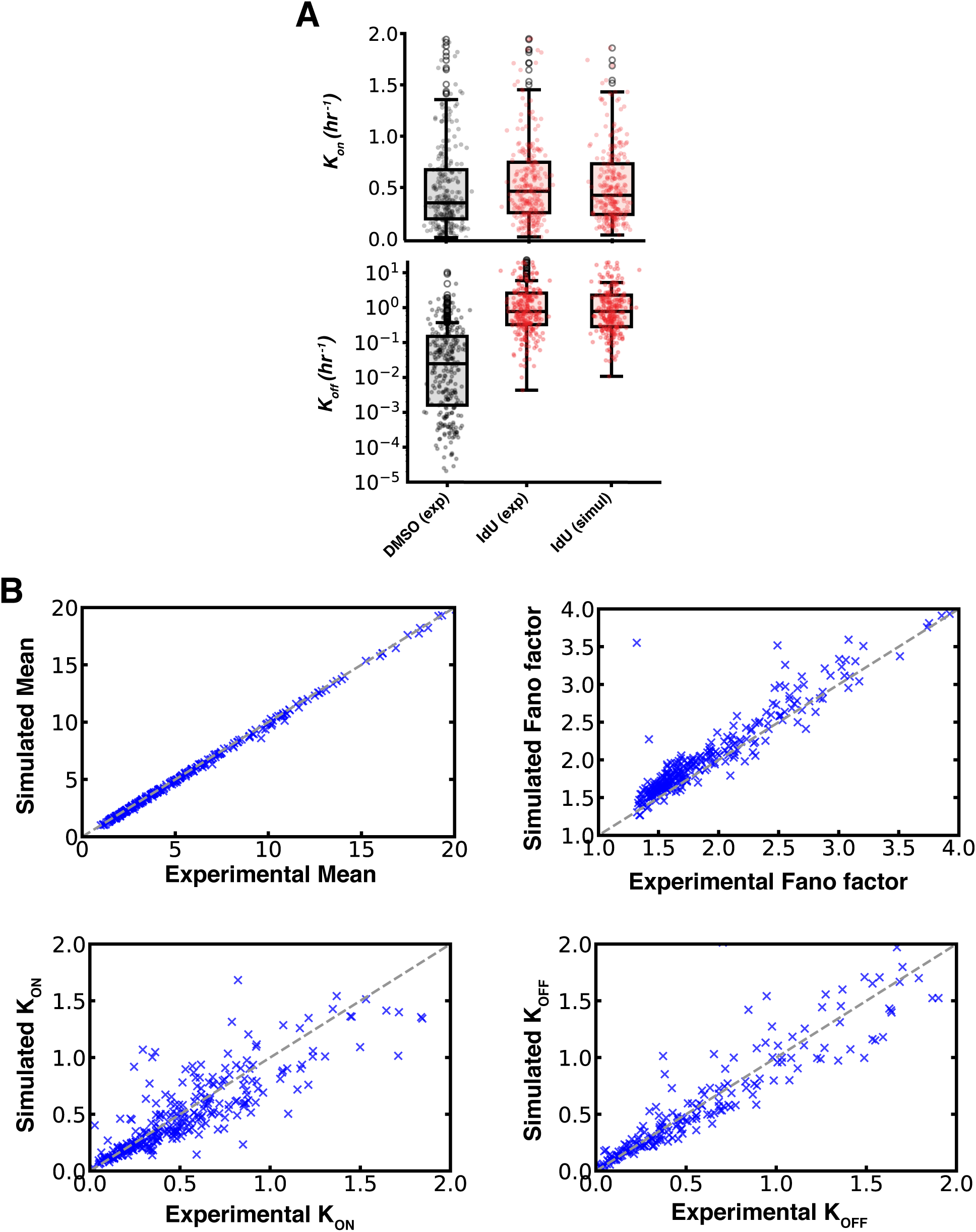
TCR model provides unifying mechanism for noise-enhancement of genes with different bursting kinetics. **(A)** Experimental values (exp) of macroscopic *K*_*ON*_ and *K*_*OFF*_, as derived from a moments-matching technique applied to scRNA-seq data, are compared to predicted values (simul) derived from simulations of TCR model. Each point represents a gene. Boxplots show median *±* interquartile range of parameter values. **(B)** Experimental values of mean, Fano factor, *K*_*ON*_, and *K*_*OFF*_ (based on scRNA-seq data) are compared to simulated values derived from TCR model.

**Figure S26:**
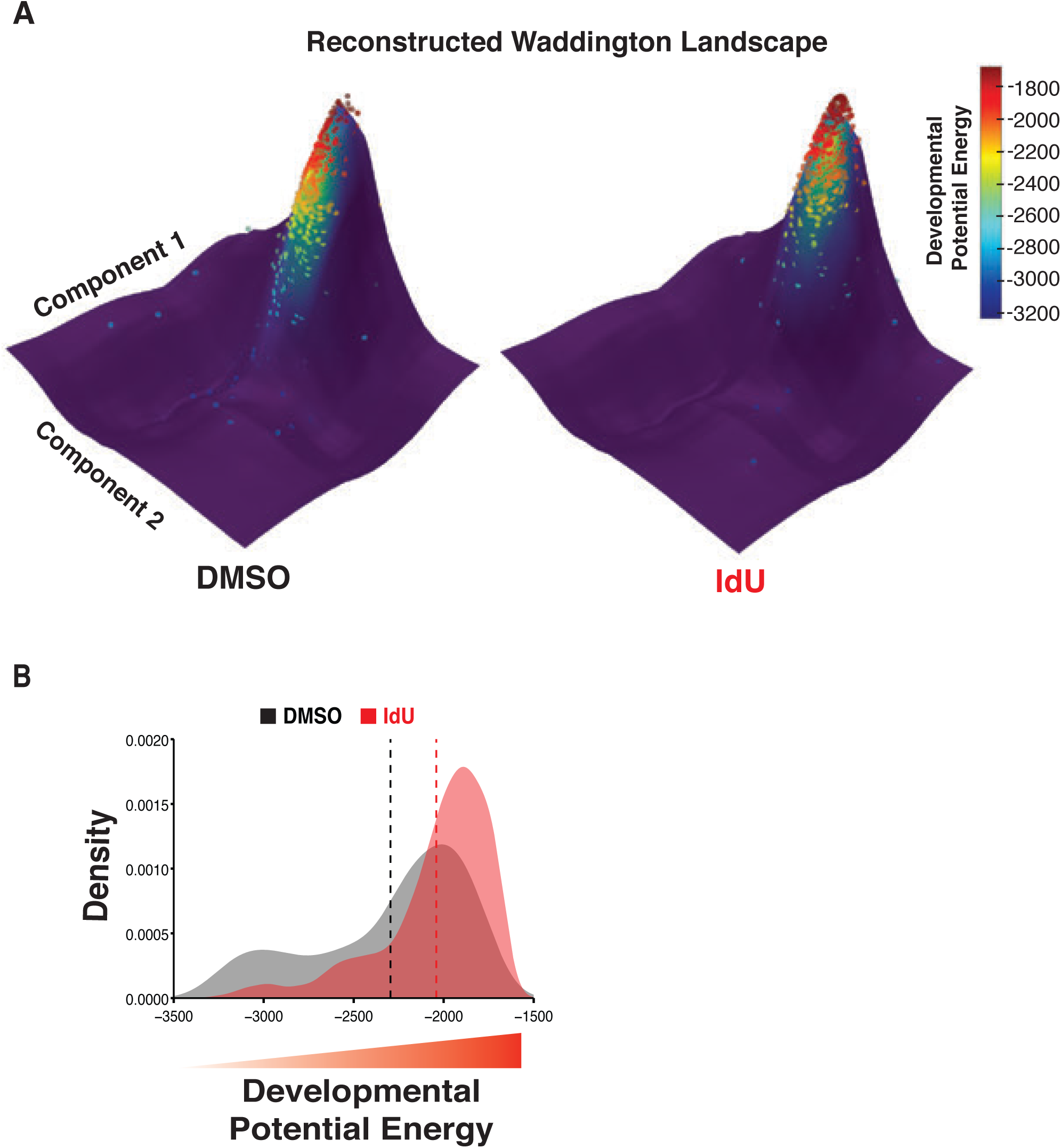
Amplification of transcriptional fluctuations destabilizes cellular identity resulting in greater cellular plasticity. **(A)** Reconstruction of Waddington’s landscape using scRNA-seq data of mESCs treated with DMSO (left) or IdU (right). Each point represents a cell. A Gaussian process latent variable model (GP-LVM) was used for dimensionality reduction to create a 2-D map of cell clustering, represented by component 1 (y-axis) and component 2 (x-axis). The z-axis represents the calculated potential energy (distance from an attractor) of a cell’s gene expression state with lower values indicating greater proximity to an attractor and thus lower developmental potential. Cells are colored according to their height (developmental potential energy, z-axis value) on the landscape as denoted by the associated color bar. Underlying shading of landscape represents density of points with purple being the least dense and yellow being the most dense. **(B)** Distributions of developmental potential energy (z-axis values from Waddington landscape in panel A) for mESCs treated with DMSO or 10µM IdU for 24 hours. Dashed vertical lines signify the mean of each distribution, with IdU-treated cells demonstrating greater potential energies.

**Figure S27:**
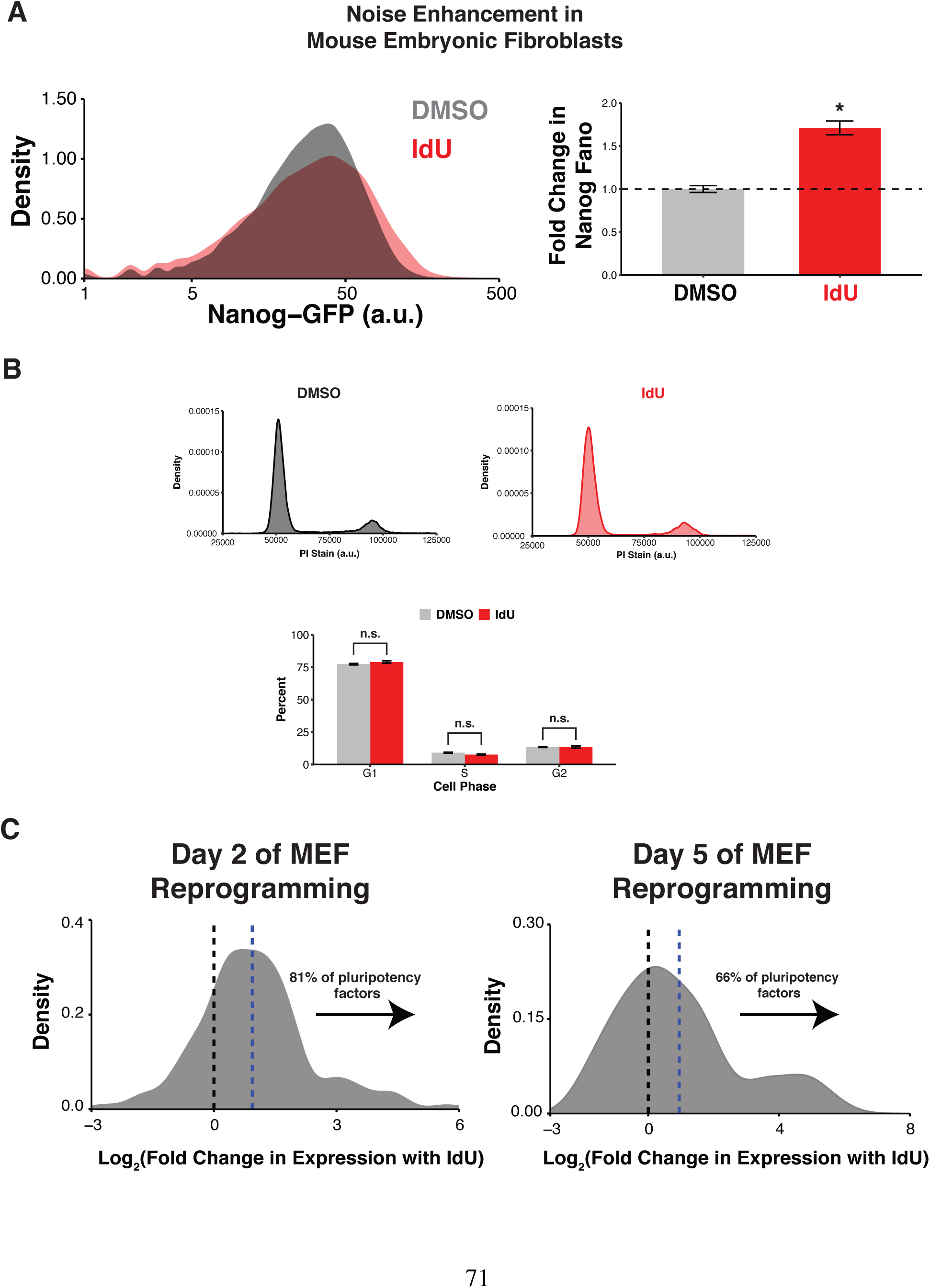
IdU treatment enhances conversion of mouse embryonic fibroblasts (MEFs) into induced pluripotent stem cells (iPSCs). **(A)** Secondary MEFs with the endogenous Nanog locus tagged with GFP were treated with 4µM IdU or equivalent volume DMSO for 48 hours in MEF media. (Right) Representative flow cytometry distributions of Nanog-GFP expression in secondary MEFs after 48 hour treatment with IdU or DMSO. (Left) Quantification of Nanog Fano factor demonstrates that IdU treatment increases expression variability as compared to DMSO control (*p = 0.003, by a two-tailed, unpaired Student’s t test). Data represent mean and SD of three biological replicates. **(B)** (Top) Representative flow cytometry distributions of propidium iodide staining for Nanog-GFP secondary MEFs treated with either DMSO or 4µM IdU for 48 hours in MEF media. No signs of aneuploidy are visible, indicating Nanog expression variability is not due to cell-to-cell variability in gene copy numbers. (Bottom) Percent of cells in each phase of the cell cycle for DMSO and IdU treatments based on propidium iodide staining. IdU treatment does not alter cell-cycle progression, indicating enhanced reprogramming is not due to accelerated cellular division. Data represent mean and SD of three biological replicates. P values were calculated using a two-tailed, unpaired Student’s t test. **(C)** Bulk RNA-seq was conducted on days 2 and 5 of doxycycline-induced reprogramming of secondary MEFs supplemented with 4µM IdU or equivalent volume DMSO for the first 48 hours. Distributions of fold change in expression for 129 pluripotency genes (taken from Mouse Genome Informatics, gene ontology term: 0019827) in the IdU condition as compared to the DMSO control are shown. Dashed blue line represents mean of distribution. 81% and 66% of the pluripotency factors show increased expression with the addition of IdU as compared to DMSO control at days 2 and 5 of reprogramming, respectively. Noise amplification during early stages of reprogramming accelerates activation of pluripotency network.

## Captions for supplementary tables S1-S6

**Table S1** (attached separately)

Sequences of smRNA-FISH oligonucleotide probes for first intron of Nanog and GFP.

**Table S2** (attached separately)

Inferred macroscopic kinetic rates of 2-state random telegraph model for Nanog in DMSO and IdU conditions.

**Table S3** (attached separately)

List of nucleoside analogs that were screened for ability to increase to Nanog protein variability.

**Table S4** (attached separately)

Gene targets and sequences of gRNAs used in CRISPRi screen.

**Table S5** (attached separately)

Sequences of primers used for RT-qPCR verification of Apex1 and Tk1 knockdown.

**Table S6** (attached separately)

Concentrations and layout of compound plates used for testing of IdU, BrdU, or hmU in combination with CRT0044876.

## Notes

### Competing Interest Statement

The authors have declared no competing interest.

